# Cancer Exacerbates Chemotherapy Induced Sensory Neuropathy

**DOI:** 10.1101/667105

**Authors:** Stephen N. Housley, Paul Nardelli, Dario Carrasco, Emily Pfahl, Lilya Matyunina, John F. McDonald, Timothy C. Cope

**Affiliations:** School of Biological Sciences, Georgia Institute of Technology, Atlanta, Georgia 30332; W.H. Coulter Department of Biomedical Engineering, Emory University and Georgia Institute of Technology, Georgia Institute of Technology, Atlanta, Georgia 30332; Integrated Cancer Research Center, Parker H. Petit Institute for Bioengineering and Bioscience, Georgia Institute of Technology, 315 Ferst Drive, Atlanta, GA, 30309, USA

## Abstract

For the constellation of neurological disorders known as chemotherapy induced peripheral neuropathy, mechanistic understanding, and treatment remain deficient. Here we present the first evidence in preclinical investigation of rats that chronic sensory neuropathy depends on non-linear interactions between cancer and chemotherapy. Global transcriptional profiling of dorsal root ganglia revealed differential expression, notably in regulators of neuronal excitability, metabolism and inflammatory responses, all of which were unpredictable from effects observed with either chemotherapy or cancer alone. Systemic interactions between cancer and chemotherapy also determined the extent of deficits in sensory encoding and ion channel protein expression by single mechanosensory neurons, with the potassium ion channel Kv3.3 emerging as a potential contributor to sensory neuron dysfunction. These original findings identify novel contributors to peripheral neuropathy, and emphasize the fundamental dependence of neuropathy on the systemic interaction between chemotherapy and cancer.

## Introduction

Chemotherapy can achieve high rates of survival in patients with common cancers^1^ but often causes severe side effects, including neuropathy, that can limit its use^2-4^. Debilitating sensory disorders, including pain, paraesthesia, and somatosensory loss, reduce quality of life for many patients and can persist for months or years after discontinuing chemotherapy^5-7^. These neurological disorders occur for up to 80% of patients receiving commonly used antineoplastic agents, notably antitubulins and proteasome inhibitors, as well as platinum-based compounds^7,8^, which are the prescribed adjuvant treatment in 50% of cancer cases worldwide^9,10^. There are no options available for the prevention of sensory disorders and the few available pharmacological treatments that focus on symptomatic management are largely ineffective^11^. Thus, there is an urgent need to better understand the pathogenesis of sensory dysfunction to aid development of mechanism-based therapies.

Preclinical studies of chronic neuropathy in experimental models of cancer are unavailable, possibly because of presumption that chemotherapy alone is sufficient to explain the neuropathology. A recently published meta-analysis identified 341 preclinical studies of chemotherapy induced neuropathy, and none assessed interaction between chemotherapy and cancer^12^. Omitting cancer from preclinical study produces a fundamental gap in knowledge that may explain why treatments for neuropathic side effects of chemotherapy have been unsuccessful in patients with cancer. Cancerous tumors located outside the nervous system can induce cognitive disability and peripheral nervous system dysfunction independent of chemotherapy, surgery, or co-morbidities^13,14^. Moreover, cancer provokes dysregulation in immune, metabolic, oxidative and neuronal excitability^15,16^, all of which are identified as targets through which chemotherapy produces neuropathy^2^. Convergence of cancer and chemotherapy on the same biological processes seems likely to yield non-linear interactions, leading us to hypothesize that clinically relevant neuropathy emerges from codependent actions of cancer and chemotherapy.

We tested our hypothesis by uncoupling the independent and combinatorial effects of cancer and chemotherapy on sensory neurons in studies of rats that are not feasible in human patients. Here we present original findings to show that sensory neuropathy depends on complex systemic interactions between cancer and chemotherapy. By combining cancer and chemotherapy, we reproduce the relevant clinical condition in preclinical study of rats to produce the first transcriptional profile of nervous tissue, to discover recurrent metabolic reprogramming and perturbed inflammatory and ion channel processes distinct from the independent effects of cancer or chemotherapy. Parallel codependency is found in novel neuropathic responses of mechanosensory neurons, both in their immunohistochemistry for selected voltage-gated potassium ion channels and in electrophysiological measures of encoding naturalistic stimuli *in vivo*. Collectively, our data show for the first time that sensory neuropathy emerges from codependencies between the systemic effects of cancer and chemotherapy. Apart from identifying new molecular targets for restoring function to mechanosensory neurons, our findings provide the fundamental insight that accounting for chemotherapy-cancer combinatorial effects on the somatosensory nervous system is prerequisite for developing meaningful treatment or prevention of neuropathy.

## Results

### Transcriptional profiling of sensory neurons

We first profiled the transcriptomes of lumbosacral (L4-S1) dorsal root ganglia (DRG), where sensory neurons (Fig. 1b) are vulnerable to chemotherapy^2^. We studied wild-type (*Apc*^*WT*^) rats and rats carrying an *Apc gene* mutation (*Apc*^*Pirc/+*^)^17^ associated with development of colon cancer (Extended Data Fig. 1^18^), not only in these rats, but also in 80% of patients with colorectal cancer^18^). Rats in each group were randomized to receive injections of oxaliplatin (OX), a platinum based compound known to induce neuropathy in patients with cancer and shown in our previous studies to induce chronic movement disorders^19^ or control. OX was administered by intraperitoneal injection on a human-scaled dose schedule (Fig. 1a)^19^. The OX treatment schedule resulted in four experimental groups used for all studies: *Apc*^*WT*^+control, *Apc*^*WT*^+OX, *Apc*^*Pirc*/+^+control and *Apc*^*Pirc*/+^+ OX (Fig. 1a).

**Figure 1.**
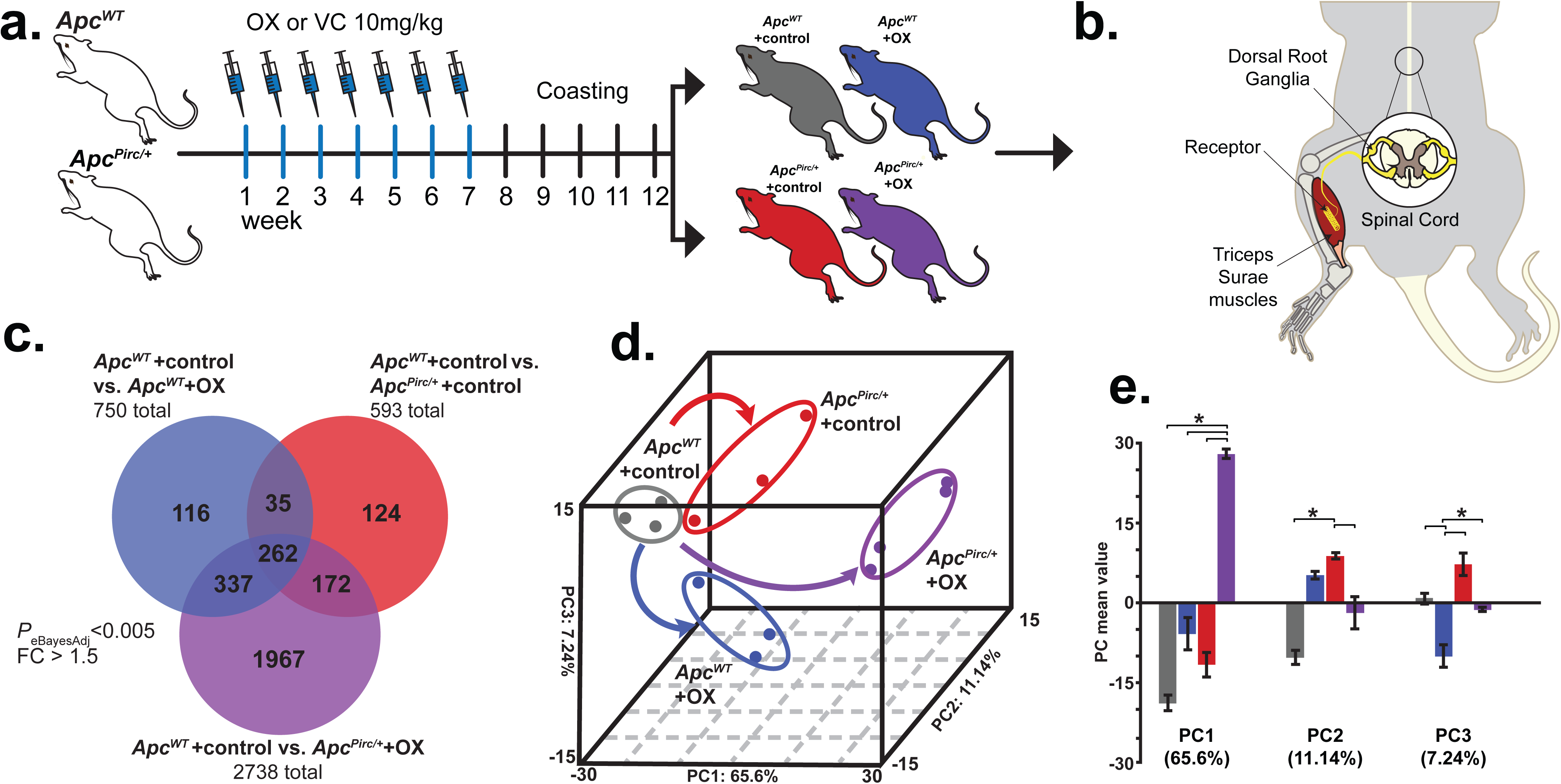
Cancer-chemotherapy codependence amplifies transcriptional dysregulation in rat sensory neurons. **a**, Schematic of treatment schedule of oxaliplain (OX) and vehicle control (VC) generating four independent experimental groups in wild-type (*Apc*^*WT*^) and autosomal dominant mutation of the adenomatous polyposis coli gene (*Apc*^*Pirc/+*^): *Apc*^*WT*^+control, *Apc*^*WT*^+OX, *Apc*^*Pirc/+*^+control, and *Apc*^*Pirc/+*^+OX rats. **b**, Diagram of experimental targets of lumbosacral dorsal root ganglia for assaying transcriptomes and cell-specific protein expression in rat models. **c**, Venn-diagram distribution of significantly differentially expressed genes from the lumbosacral dorsal root ganglia among all groupwise contrasts. Significance determined by both *P*<0.005 and fold-change (FC) >1.5 using a Bayesian moderated linear fixed effects model (eBayesAdj). **d**, Group correlation determined by principal-component (PC) analysis of significantly differentially expressed genes. Genes were visualized in the new 3D composite space created by PC1-3. Effects of independent chemotherapy (OX) or cancer (*Apc*^*Pirc/+*^) and their combinatorial (*Apc*^*Pirc/+*^+OX) treatment are indicated by curved arrows; dots represent individual animals. **e**, Bar graphs of the mean values of PC1, PC2, and PC2. Error bars represent s.e.m. * indicates statistically significant differences between experimental groups as empirically derived from hierarchical Bayesian model (*stan_glm*): 95% highest density intervals do not overlap between groupwsie contrasts.

We employed an empirical Bayesian (eBayes) linear model^*20*^ to modify residual variances on a total of 31,042 (22,850 named) genes and identified 3,013 differentially expressed genes (DEG) across all groupwise contrasts (*P*_eBayesAdj_ < 0.005). Subsequent *post hoc* analysis identified DEG from all possible contrasts. Overlap between DEGs retrieved from individual comparisons is illustrated in Figure 1c. In keeping with the notion that chemotherapy and cancer act through similar molecular mechanisms, we found significant overlap for 297 DEG affected independently by cancer and chemotherapy (Fig. 1c). Of note, we found that 65% (n=1,967) of all DEG were uniquely affected when cancer and chemotherapy interacted (*Apc*^*Pirc/+*^+OX) (Fig. 1c). This finding demonstrates the first evidence that cancer exacerbates genetic dysregulation induced by chemotherapy, unmasking a more than three-fold increase in DEG.

Next, we performed two independent downstream analyses (Extended Data Fig. 2) to test for interaction and determine which biologic processes might be involved in pathogenesis of sensory disorders. First, we subjected the eBayes filtered database to unsupervised principal component (PC) analysis. DEG were visualized in the new 3D composite space created by PC1-3 (Fig. 1d) that explained 83.98% of the variance and clearly segregated the four experimental groups. Inspection of individual PCs revealed significant evidence of non-linear interactions between cancer and chemotherapy captured in PC1 (65.6% of explained variance), whereas PC2 (11.14%) and PC3 (7.24%) separated the effects of cancer and chemotherapy, respectively (Fig. 1e). Second, we drove unsupervised hierarchical clustering onto the PC1 solution to identify groups of coregulated genes that express interaction^21^ (Fig. 2a). Analysis identified two clusters of genes displaying distinct, but internally consistent expression profiles clearly representing patterns of interaction (Fig. 2a and Extended Data Table 1).

**Figure 2.**
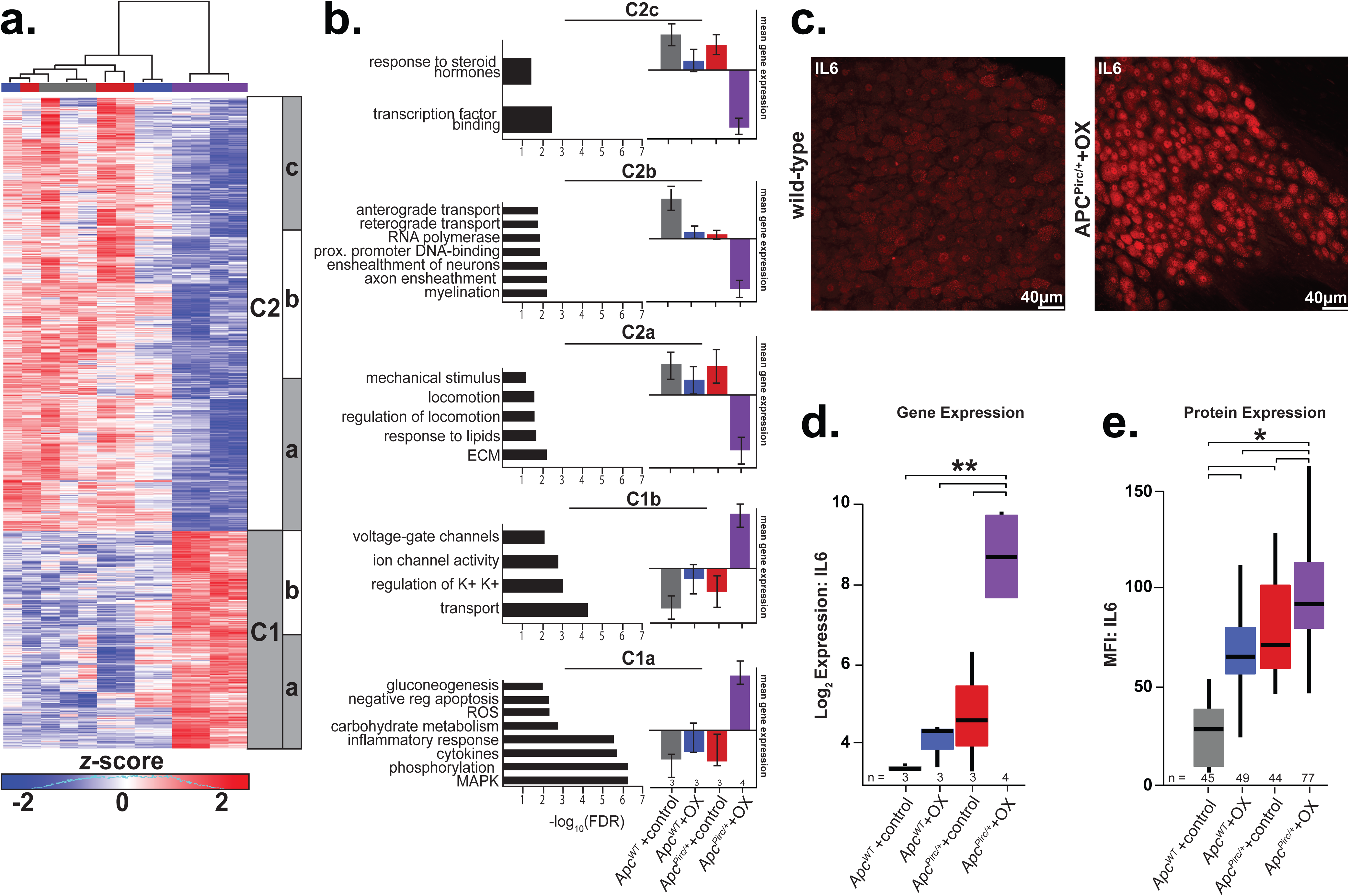
Cancer-chemotherapy interaction manifests as disruption of distinct biological processes in rat sensory neurons. **a**, Heatmap of the top differentially expressed genes in the sensory neurons identified by PC1 driven hierarchical-clustering (biological triplicates n= 3 for *Apc*^*WT*^ +control, *Apc*^*WT*^+OX, *Apc*^*Pirc/+*^+control and four biologic replicates for *Apc*^*Pirc/+*^+OX rats). Data were mean-centered and s.d. normalized before clustering; upregulated and downregulated genes (row) are shown in red and blue, respectively. White and grey bars (C1-2) display the main clusters from gene level hierarchical-clustering and were chosen for clarity, while subclusters (a-c) highlight unique biologic characteristics within the main clusters. **b**, Clusters showing distinct up- or downregulation of key biological processes (Gene Ontology (GO) terms) and pathways (Kyoto Encyclopedia of Genes and Genomes (KEGG)). Selected pathway and gene ontology terms significantly associated (false discovery rate [FDR] ≤ 0.25) with misregulated transcripts in individual clusters are listed on the left of each gene cluster with significance indicated by horizontal bars (-log10(FDR)). The vertical bar graphs on the right compare mean±s.d. of gene expression levels in each subcluster (**Extended Data Table 1**). **c**, Confocal image of a dorsal root ganglia immunostained against interleukin 6 (IL6). Scale bar, 40 μm. Gene expression (**d**) and receptor protein expression (**e**, mean fluorescence intensity (MFI)) of IL6 as determined by transcriptional profiling and cell-specific immunohistochemistry of large diameter sensory neurons (>30μm). Group sample size inset below plot. * indicates statistically significant differences between experimental groups as empirically derived from hierarchical Bayesian model (*stan_glm*): 95% highest density intervals do not overlap between groupwsie contrasts. ** indicates *P*_eBayesAdj_<0.005 and fold-change (FC) >1.5 using a Bayesian moderated linear fixed effects model. Data presented as mean±s.d.

We then subjected clusters of named genes (Extended Data Table 1) to pathway-enrichment analysis^22,23^ and identified unique subclusters (Fig. 2a) expressing dysregulation in distinct biological processes (Fig. 2b). In keeping with the notion that combinatorial effects of cancer and chemotherapy (OX) reflect convergence onto shared signaling pathways, genes mediating inflammatory response, e.g. interleukin 6 (*Il6)* (*P*_eBayesAdj_=7.82× 10^-6^), *Ptgs2* (*P*_eBayesAdj_=0.0028) and *Cxcr4* (*P*_eBayesAdj_=3.74× 10^-7^), and reactive oxygen species processes, e.g. *Hmbox1* (*P*_eBayesAdj_=0.001), were found to be among the most upregulated in cluster 1a (C1a) (Fig. 2b). Notably, we observed substantial induction of genes encoding glycolytic and carbohydrate metabolic processes (FDR ≤ 1.83×10^-3^, C1a; Fig. 2b) in the *Apc*^*Pirc/+*^+OX neurons. Coincident with these changes was a statistically significant down-regulation of genes mediating lipid metabolism (C2a), suggestive of pervasive metabolic dysfunction beyond the small subset of dysregulated mitochondrial related processes that we found in response to *Apc*^*WT*^+OX or *Apc*^*Pirc*/+^+control alone (Fig. 2b; Extended Data Fig. 3a,b) and similar to dysfunctional effects previously suggested^2^. Among additional unique transcripts downregulated in *Apc*^*Pirc/+*^+OX neurons, Fig. 2b highlights multiple deficits in processes related to extracellular matrix control, mechanical stimulus transduction, (C2a), DNA-binding, RNA polymerization, (C2b), transcription factor activity, hormone signal transduction (C2c and Extended Data Table. 1). Furthermore, we observed genes in C1b and 2a that represent specific dysregulation of neuronal and sensorimotor function, e.g. ion-channel genes determining neuronal excitability along with coincident down-regulation of locomotion. These data provide the first evidence that codependent neuropathy induces broad metabolic reprogramming, e.g. increased glycolytic and decreased lipid metabolism, consistent with cancers capacity to corrupt metabolic processes^24,25^, which extends knowledge beyond the limited impairment observed in mitochondria.

**Figure 3.**
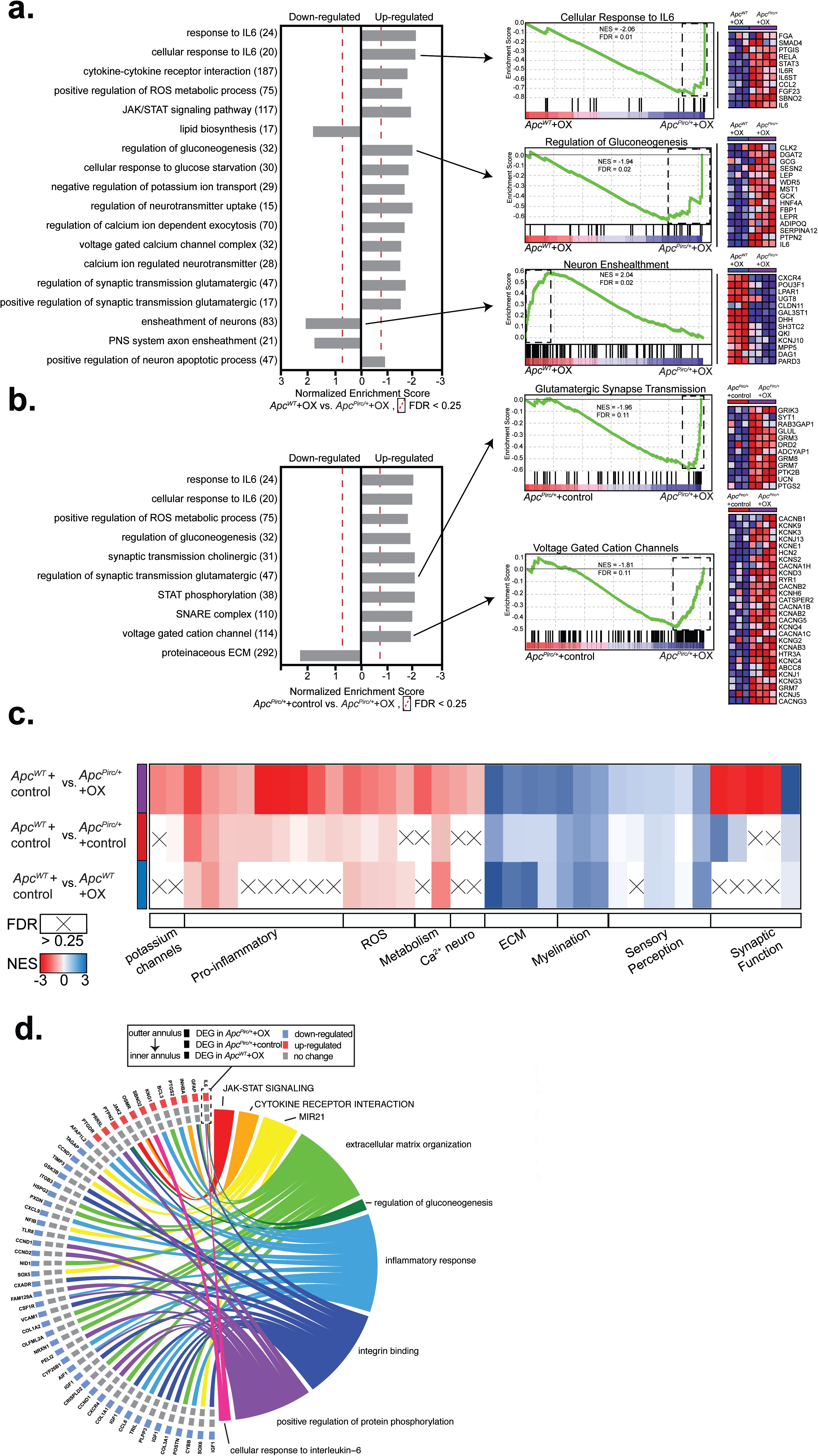
GSEA and pathway analyses test codependence of cancer and chemotherapy. Enriched gene sets in sensory neurons identified by gene set enrichment analysis (GSEA: MSigDB, C2-CP: canonical pathways; C3-MIR: microRNA targets; C3-TFT: transcription factor targets; C5-BP: GO biological process; C5-CC: GO cellular component; C5-MF: GO molecular function C7: immunologic signatures gene sets from the comparison of **a**, *Apc*^*WT*^+OX and *Apc*^*Pirc/+*^+OX and **b**, *Apc*^*Pirc/+*^+control and *Apc*^*Pirc/+*^+OX groups. Individual frames (horizontal grey bars) identify functional related (internally consistent) groups of gene sets expressing distinct up- or downregulation as compared with *Apc*^*WT*^+OX. Number of overlapping genes displayed in parentheses to the right of the corresponding GO term. Representative enrichment plots are shown in the middle column. Heat-map representation of leading-edge genes is shown on right. **c**, Heat-map representation of enriched individual gene sets, identified by GSEA, in sensory neurons for chemotherapy (bottom rown), cancer (middle row), and the combination of cancer (*Apc*^*Pirc/+*^) and chemotherapy (OX) (top row). Gene sets are grouped into functionally related cohorts. Gene sets not reaching statistical significance indicated by (X). **d**, Chord diagram shows enriched pathways (Database for Annotation, Visualization, and Integrated Discovery; DAVID) on the right, and genes contributing to enrichment are shown on the left. Squares on left indicate genes differentially expressed in *Apc*^*WT*^+OX, *Apc*^*Pirc/+*^+control, or *Apc*^*Pirc/+*^+OX groups compared to control (see the key for gene differential expression). Statistical significance determined by permutation testing with normalized enrichment score (NES) and Benjamini–Hochberg false discovery rate (FDR) < 0.25. DEG, differentially expressed genes; ROS, reactive oxygen species: ECM, extracellular matrix.

Markers for nerve degeneration, inconsistently observed in patients^26^ treated with antineoplastic drugs exhibited mixed results. While we observed few significantly downregulated genes associated with myelination processes (C2b; *P* ≤ 7.12×10^-3^, Fig. 2b), we also found significant induction of processes involved in the preservation of neurons (e.g. enrichment of negative regulation of apoptosis *P* ≤ 5.96 × 10^-3^) in C1a. Mixed results from these data suggests heterogeneous neuronal populations in the DRG may contain discrete subpopulations with selective vulnerability whereas others are resistant to degeneration, which is consistent with mixed results seen in human studies^26^ and highlights the need for future cell-type specific investigations that are currently unavailable.

### Dysregulation is conserved at protein level

Because inflammatory pathways play multiple roles in regulating neuron function^27^, we focused attention on their significant response to codependent neuropathy. We selected the most dysregulated gene in the inflammatory pathways, *Il6*, for molecular validation by assessing gene—protein correspondence. Sensory neurons in dorsal root ganglia were immunolabeled with monoclonal antibodies targeting IL6 (Fig. 2c). As predicted from the transcriptional profiling (Fig. 2d), the IL6 protein was constitutively expressed in *Apc*^*WT*^+control rats at lower levels than in *Apc*^*Pirc/+*^ control animals or Apc^WT^+OX animals, and at significantly higher levels in *Apc*^*Pirc/+*^+OX neurons (Fig. 2e), demonstrating that codependent neuropathy was conserved in protein expression (Fig. 2e and Extended Data Fig. 4). Previous studies implicate IL6’s capacity to mediate hyperexcitability and morphological changes associated with neuron death^28^. However, studies restricted investigation to acute neurotoxicity, which limits generalizability to symptoms that persist long after treatment. Our data present the first evidence that *Il6* (IL6) expression levels remain elevated long after treatment cessation without observable morphological changes. We show that codependent neuropathy significantly increases *Il6* (IL6) when compared to chemotherapy or cancer alone. In light of IL6’s capacity to decrease excitability following chronic exposure^28^, our data suggest that neuronal dysfunction as a result of codependent neuropathy may be expressed as hypo-excitability, in contrast with previous work^7^.

### Broad dysregulation of neuron transcriptomes

We then used gene-set enrichment analysis (GSEA) as a second independent tool to investigate the cellular processes underlying codependent neuropathy. We initially focused on the comparison of *Apc*^*WT*^+OX and *Apc*^*Pirc/+*^+OX (Fig. 3a) and *Apc*^*Pirc/+*^+control and *Apc*^*Pirc/+*^+OX (Fig. 3b) and interrogated entire data sets against the Molecular Signatures Database (Extended Data Table 2). Two novel conclusions can be drawn from the GSEA findings. First, GSEA independently validated codependencies of cancer and chemotherapy in the DEG identified by transcriptional profiling using paired unsupervised analysis (Figs. 3a and 3b). GSEA corroborated downregulation of lipid metabolic pathways (normalized enrichment score (NES)=1.73) and upregulation of glycolytic pathways (NES=-1.94) in *Apc*^*Pirc/+*^+OX sensory neurons when compared with either chemotherapy (Fig. 3a) or cancer alone (Fig. 3b). Focusing analysis on pathways directly involved in neuronal signaling confirmed dysfunction in peripheral neuron ensheathment in the absence of evidence supporting neuron specific apoptotic response (FDR>0.83) (Fig. 3a). Second, GSEA exposed dysregulation of genes relevant to neuronal excitability particularly potassium ion channels and transporters, and others participating in synaptic communication, e.g. SNARE and glutamatergic transmission (Fig. 3a,b). Dysregulation of these and other gene sets (e.g. proinflammatory chemokines) was not predicted from the independent effects of cancer or chemotherapy alone. (Fig. 3c,d and Extended Data Table 2). Unlike previous studies that consistently report dysregulated voltage-gated sodium channels, we find little evidence of differentially expressed sodium channels or disrupted regulatory pathways^29^. Instead, we find targeted DEG related to potassium ion channels unlike those previously identified. We draw two conclusions from these data. First, ion channel clusters are differentially vulnerable to codependent neuropathy. Second, chronic neuropathy may be mechanistically linked to dysregulation of ion channels distinct from those identified in acute preparations.

Taken together, our analyses indicate significant dysregulation of genes and proteins mediating core cellular processes of *Apc*^*Pirc/+*^+OX sensory neurons. Our integrated transcriptomics and protein level findings confirmed that cancer and chemotherapy effect common processes^15,30,31^. Moreover, our findings support our central hypothesis that clinically relevant neuropathy depends on systemic interaction between cancer and chemotherapy by showing for the first time that codependence exacerbates neuropathy and revealing it operates through distinct mechanistic pathways, unpredicted from cancer or chemotherapy alone.

### Novel ion channel dysfunction in Apc^Pirc/+^+OX

Sensory deficits, e.g. somatosensory and hearing loss, are commonly reported by patients long after cessation of chemotherapy, but knowledge of the combined effects of cancer and chemotherapy is missing. Mechanosensory systems, e.g. cochlear hair cells, and various somatosensory neurons in skin and muscle are responsible for encoding this information and exhibit deficits following chemotherapy^8^. We therefore next focused on identifying the combined effects of cancer and chemotherapy on a representative member of mechanosensory neurons, called muscle spindles. Muscle spindles share many of the molecular mechanisms^32^ underlying mechanotransduction and encoding found in this broad class of sensory neurons (e.g. hair cells^33^, Merkel^34^). Further, signaling by these neurons encodes sensory features of muscle mechanics necessary for perceiving (proprioception) and coordinating body position and movement, functions which, when impaired, have the potential to explain persistent disorders in patients^35^. We performed targeted analyses of our transcriptome database by querying DEGs identified in *Apc*^*Pirc/+*^+OX rats compared to all other groups against genes encoding proteins known to be constitutively expressed in mechanosensory neurons that mediate unique contributions to neuronal signaling (Extended Data Fig. 5)^32,36,37^. We found no evidence for independent nor combinatorial effects of cancer or chemotherapy at the genetic level (Fig. 4a and Extended Data) for channels mediating mechanotransduction (e.g. *Asic2, Piezo2, ENaCs*), signal amplification, and spike encoding (e.g. *Scn1a* (Nav1.1), *Scn8a* (Nav1.6), *Scn9a* (Nav1.7), *Cacna1s,-c,-d, -f (*Cav1.1-4), *Kcnn2* (SK2), *Kcna1* (Kv1.1))

**Figure 4.**
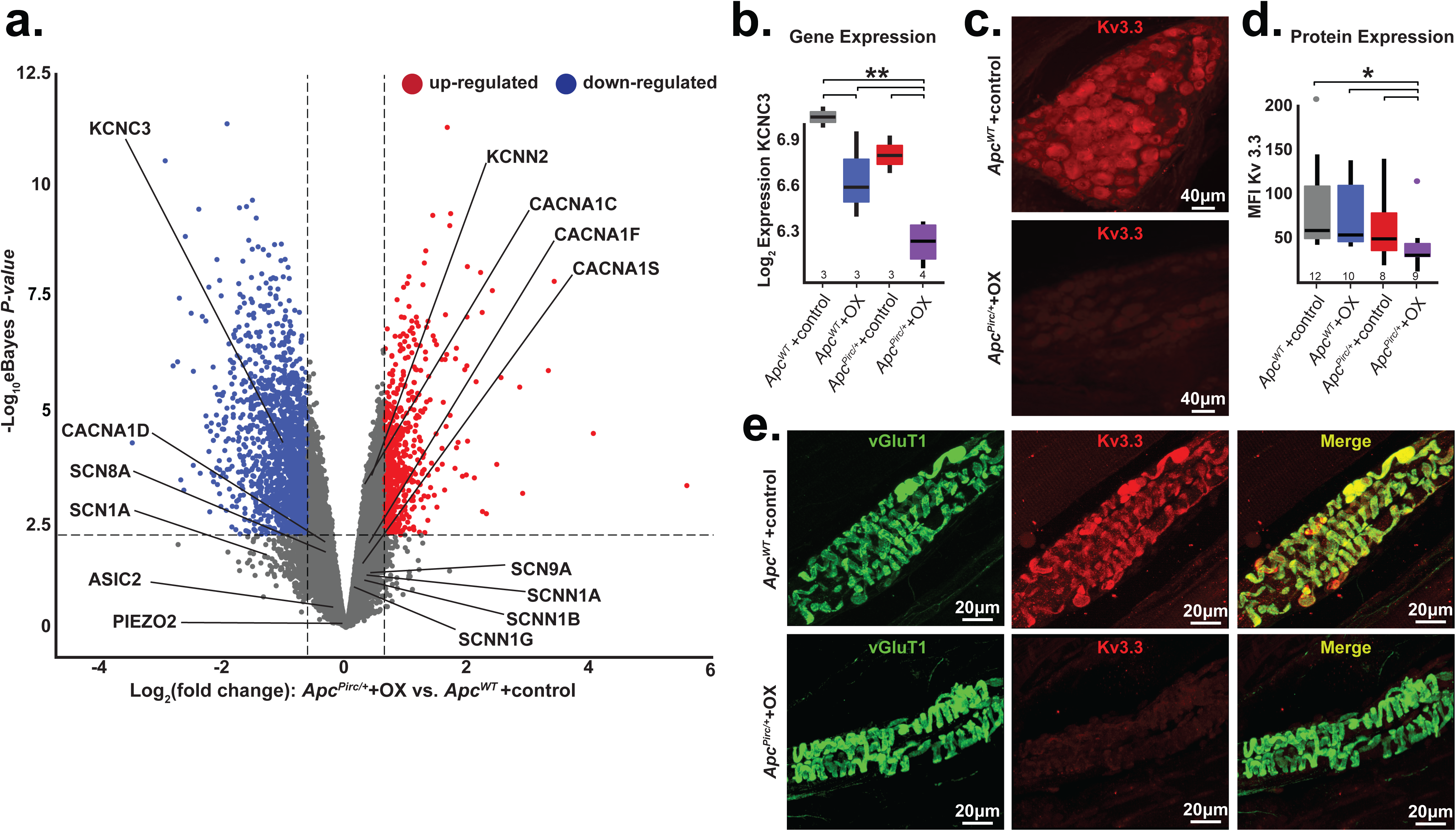
Targeted transcriptional analysis reveals mechanisms for cancer-chemotherapy interaction. **a**, Volcano plot comparing up- and downregulated genes in *Apc*^*Pirc/+*^+OX and *Apc*^*WT*^ +control sensory neuron transcriptomes. Annotation of genes involved in mechanotransduction, signal amplification, action potential generation and maintenance is shown. Significance was determined as Benjamini–Hochberg FDR < 0.01 and log_2_fold change 2: 1. **b**, Gene expression of *Kcnc*3 encoding Kv3.3, as determined through transcriptional profiling (Kcnc3: 1369133_a_at). **c**, Confocal images of dorsal root ganglia immunolabeled against Kv3.3 illustrates expression of *Apc*^*Pirc/+*^+OX relative to *Apc*^*WT*^ +control neurons.**d**, Quantification of receptor protein expression (mean fluorescence intensity (MFI)) of Kv3.3 determined through averaging over all annulospiral endings (617 from 12 *Apc*^*WT*^ +control neurons, 434 from 10 *Apc*^*WT*^ +OX neurons, 459 from 8 *Apc*^*Pirc/+*^+control neurons, and 362 from 9 *Apc*^*Pirc/+*^+OX neurons) for each neuron per experimental group. **e**, Confocal images of mechanosensory receptor ending of proprioceptive sensory neurons immunolabeled against VGluT1 (green), Kv3.3 (red), and merged (yellow) illustrates protein expression levels and distribution in *Apc*^*Pirc/+*^+OX relative to *Apc*^*WT*^ +control. * indicates statistically significant differences between experimental groups as empirically derived from hierarchical Bayesian model (*stan_glm*): 95% highest density intervals do not overlap between groupwsie contrasts. ** indicates *P*_eBayesAdj_<0.005 and fold-change (FC) >1.5 using a Bayesian moderated linear fixed effects model. Data presented as mean±s.e

Next, we performed cell-specific immunohistochemical analyses to test gene–protein correspondence. We found no evidence to suggest dysregulated protein expression nor did evidence emerge to suggest altered protein distributions (Extended Data Figs. 6-8) for most proteins, including those mediating mechanotransduction (ASIC2), signal amplification (Nav1.1, Nav1.6, Nav1.7), spike encoding (Kv1.1), and mechanosensory sensitivity (VGLUT1). These findings demonstrate that many of the molecular mechanisms responsible for mechanotransduction and determining excitability of mechanosensory neurons were preserved in the *Apc*^*Pirc/+*^+OX rats. These findings narrow the field of candidate mechanisms for which treatment might reasonably restore normal function.

Having exhausted examination of proteins known to be expressed in mechanosensory neurons, we extended our search to include ion channels yet to be identified, but known to regulate neuronal signaling in other classes of neurons^38-41^. As one of the most dysregulated voltage-gated ion channels, *Kcnc3* (encoding protein Kv3.3) emerged as a novel candidate (Fig. 4a, b, Extended Data Fig. 9). Pharmacologic and genetic perturbation of Kv3.3 (*Kcnc3*) impairs neuronal signaling^40,41^ and is causally linked to ataxias^42^ that are consistent with functional deficits observed in patients with cancer long after cessation of chemotherapy. We then immunolabeled sensory cell bodies in dorsal root ganglia and their receptor endings taken from *Apc*^*WT*^+control with monoclonal antibodies targeting Kv3.3, which revealed the first evidence of this ion channel in all large diameter cell bodies in the dorsal root ganglia that supply mechanosensors (Fig. 4c Extended Data Fig. 10) and endings (Fig. 4e). Results of comparison across experimental groups indicate that Kv3.3 expression was significantly lower in *Apc*^*Pirc/+*^+OX compared with *Apc*^*WT*^+control (Fig. 4d,), *Apc*^*Pirc/+*^+control, or *Apc*^*WT*^+OX (Extended Data Fig. 10), demonstrating that codependent dysfunction is conserved in cell-type specific protein expression. Moreover, conserved Kv3.3 expression in neuromuscular junctions suggests that Kv3.3 dysfunction was constrained to sensory neurons (Extended Data Fig. 11). This is the first evidence implicating Kv3.3 in the development of neuropathy and the first data, to our knowledge, demonstrating a channelopathy persists long after treatment cessation. The capacity of Kv3.3 to drive fast spiking and to enhance transmitter release – processes that are required for mechanotransduction and spike encoding– suggests that this ion channel might provide both a potential novel mechanism for the dysfunction of mechanosensory neurons and a potential target for therapy.

### Apc^Pirc/+^+OX neuronal signaling is impaired

Next, we examined whether the codependent neuropathy we discovered at gene and protein levels had negative consequences on mechanosensory function in living animals. (Extended Data Fig. 12). Applying electrophysiological methods to rats in vivo, we recorded spiking activity from single mechanosensory neurons responding to naturalistic mechanical stimuli (Fig. 5a). From these spiking responses, we collected 31 measured and derived parameters (average of four trials; n = 11 *Apc*^*WT*^+control, n=19 *Apc*^*WT*^+OX, n=20 *Apc*^*Pirc/+*^+control, n = 10 *Apc*^*Pirc/+*^+OX; Extended Data Fig. 13) from which we extract four functional features encoded by these neurons, sensitivity, dynamic, static, and history-dependent signaling information (Extended Data Fig. 13). In *Apc*^*WT*^+control rats, we found that muscle stretch elicited the spiking expected from sensory neurons in normal animals^43^ and humans^44^, specifically high frequency initial bursting at stimulus onset, increasing dynamic firing during increasing stimulus, sustained static firing with slow accommodation during the hold phase of stretch (Fig. 5b), and history-dependent reduction in spike number (Extended Data Fig. 14 and Fig. 6h).

**Figure 5.**
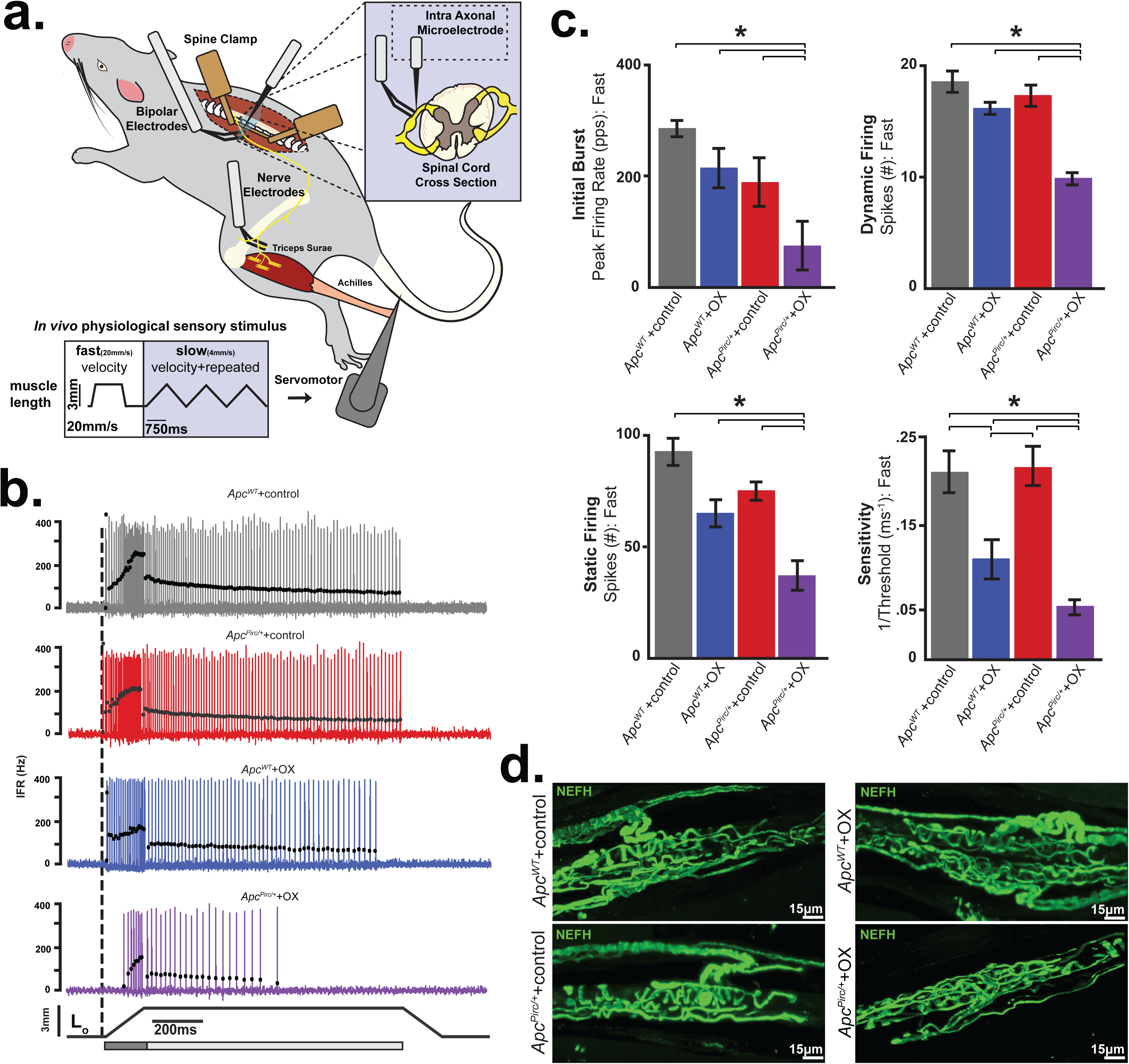
In vivo electrophysiology indicates that neuronal signaling is degraded in the cancer–chemotherapy rat model. **a**, Schematic of experimental approach for *in vivo* recordings centrally from intact proprioceptive sensory neurons firing action potentials in response to fast (20mm/s) and slow (4mm/s) naturalistic mechanical stimuli delivered in the periphery. **b**, Representative cases of spiking activity in *Apc*^*WT*^ +control, *Apc*^*WT*^+OX, *Apc*^*Pirc/+*^+control, and *Apc*^*Pirc/+*^+OX as a measure of sensory encoding. Black circles plot instaneous firing rates (IFRs) of corresponding spike (action potential) intervals. Dashed vertical line marks onset of muscle stretch (3mm) from resting length (Lo). Dynamic and static phases of naturalistic stimuli indicated by dark grey (150 ms duration after stretch command onset) and light grey (1 s duration after the dynamic phase) bars. **c**, Neuronal spiking parameters (n=31 averaged from four trials in each neuron, four shown from fast ramp stimulus) representing different features of sensory stimulus (*Apc*^*WT*^ +control (n = 11), *Apc*^*WT*^+OX (n = 19), *Apc*^*Pirc/+*^(n = 20) and *Apc*^*Pirc/+*^+OX (n = 10)): mean initial burst frequency (pulses per second [pps]) signaling stretch onset; the number of spikes during dynamic sensory stimulation (shown as a dark grey bar in **b**); the number of spikes during static sensory stimulation (shown as a light grey bar in **b**); sensitivity assessed as the inverse of latency to stimulus detection (ms^-1^), i.e. lower sensitivity corresponds to longer latency and higher threshold. **d**, Confocal image of a neurofilament heavy-chain (NEFH, in green) with immunolabeling of the terminal axon and receptor structure in *Apc*^*WT*^+control, *Apc*^*WT*^+OX, *Apc*^*Pirc/+*^+control, and *Apc*^*Pirc/+*^+OX rats. Scale bar, 15 μm. * indicates statistically significant differences between experimental groups as empirically derived from hierarchical Bayesian model (*stan_glm*): 95% highest density intervals do not overlap between groupwsie contrasts. Data presented as mean±s.e.m.

**Figure 6.**
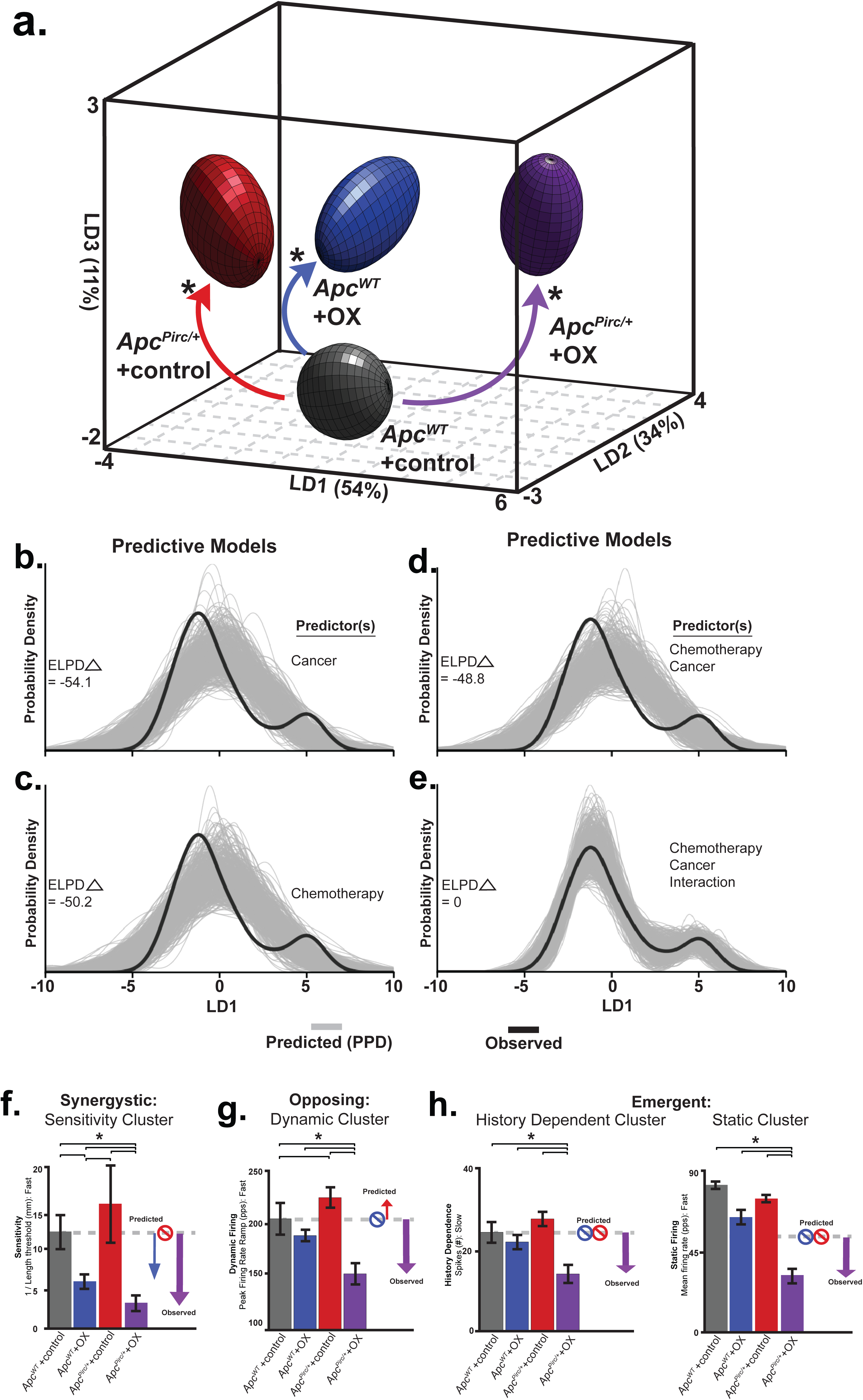
Cancer-chemotherapy codependence exacerbates sensory dysfunction beyond that predicted by cancer or chemotherapy alone. **a**, Neuronal spiking parameters (n=31, averaged from four trials, from each neuron (n=60)) representing different features of sensory stimuli were subjected to linear discriminant (LD) analysis. Neuronal signaling was visualized in the new 3D composite space created by LD1-3. 3D ellipsoids enclosing 68% of data were computed with least-squares elliptical fitting to emphasize differences between control and *Apc*^*Pirc/+*^+OX neurons. Effects of independent (OX or *Apc*^*Pirc/+*^) and combinatorial (*Apc*^*Pirc/+*^+OX) treatment are indicated by curved arrows. **b-e**, The hierarchical Bayesian model used to test for significant group differences in LD1 scores reconfigured to operate in a predictive fashion. Predictors included each model are listed to the right of each plot. Generative models in (**b-d**) utilizing one (**b**,**c**) or both independent (**d**) predictor(s), i.e. cancer (*Apc*^*Pirc/+*^) or chemotherapy (OX) for posterior prediction. The generative model in (**e**) utilizes both independent predictors and an interaction term for posterior prediction. Grey lines in both **b-e** represent 500 novel (generative) samples drawn from the posterior distributions. Black line illustrates experimentally observed mean LD1 score. Predictive accuracy was measured by calculating expected log predictive density (ELPD) for each model and benchmarked off of the highest performing model. ELPD () indicates difference from optimal model. Negative models represent worse predictive performance. **f**, Sensitivity assessed as the inverse of latency to stimulus detection (mm^-1^), i.e. lower sensitivity corresponds to longer latency and higher threshold. **g**, Peak firing rate (pps) achieved during the dynamic sensory stimulus (shown as dark grey bar in Fig. 5b). **h**, change in dynamic spike number in successive, slow stretch dynamic sensory stimulus (see Methods and Extended Data Fig. 9) and mean firing rate (pps) achieved during the static sensory stimulus (shown as light grey bar in Fig. 5b). Recordings were analyzed from *Apc*^*WT*^ +control (n = 11), *Apc*^*WT*^+OX (n = 19), *Apc*^*Pirc/+*^(n = 20) and *Apc*^*Pirc/+*^+OX (n = 10) in **b-g**, * indicates statistically significant differences between experimental groups as empirically derived from hierarchical Bayesian model (*stan_glm*): 95% highest density intervals do not overlap between groupwsie contrasts. Arrows in **f-h** indicate direction and significant differences from *Apc*^*WT*^ +control. Circles with slashes indicated no difference predicted for color-coded group. Data presented as mean±s.e.m.

We next tested whether *Apc*^*Pirc/+*^ or OX treatment alone induce signaling dysfunction. Although we find transcriptional dysregulation in *Apc*^*Pirc/+*^+control (Fig. 1c and Extended Data Fig. 3a), we found a remarkable degree of concordance with signaling in *Apc*^*WT*^+control (Fig. 5b). *Apc*^*Pirc/+*^+control neuronal signaling displayed comparable sensitivity to stimuli and showed similar static behavior across all neurons (n = 20; Fig. 5b,c) as the *Apc*^*WT*^+control. We then found that OX treatment of *Apc*^*WT*^ rats induced mild signaling deficits that were less pronounced, occurred in a small proportion of neurons (5/19), and were primarily restricted to sensitivity functional clusters (Fig. 5b,c and Extended Data Fig. 17d), validating our previous findings of restricted deficits observed in a different strain of healthy rats treated with OX alone^19^.

In contrast to the observations in *Apc*^*WT*^+control, *Apc*^*WT*^+OX, or *Apc*^*Pirc/+*^+control rats, we found drastically impaired neuronal signaling in *Apc*^*Pirc/+*^+OX rats (Fig. 5b). High frequency initial bursting was attenuated four-fold and sensitivity to stimuli was reduced by three-fold at both high (Fig. 5c) and low velocity stretch (Extended Data Fig.14, 17d). During dynamic stimuli, we found marked reduction in the number of spikes over both stretch conditions (Fig. 5c). Neurons in *Apc*^*Pirc/+*^+OX rats also failed to sustain firing (Fig. 5b). In addition, we observed fewer spikes (Fig. 5c) during rapid accommodation immediately preceding signal deletion, despite constant stimulus (Fig. 5b). We then simulated naturalistic compensation by subjecting *Apc*^*Pirc/+*^+OX neurons to repeated trials at 1, 2, and 3× background stimulus (L_o_ strain) intensity, which failed to completely rescue signaling back to *Apc*^*WT*^+control levels (Extended Data Fig. 16b,c,d). These data are the first, to our knowledge, to directly demonstrate a damaging functional interaction between the systemic effects of cancer and chemotherapy in sensory neurons in a living animal. While validating our previously findings of signaling deficits in chemotherapy alone^19^, these data indicate that restrictive signaling deficits induced by chemotherapy or cancer alone were insufficient to reproduce the magnitude, direction and number of dysfunctional parameters induced by codependent neuropathy.

### Dysfunction does not depend on degeneration

We next tested whether the impaired neuronal signaling of *Apc*^*Pirc/+*^+OX neurons might be explained by dying-back degeneration of sensory nerve terminals. While the underlying mechanisms for chemotherapy induced neuropathy are not understood, current data identify ‘dying-back’ axon degeneration as a major pathology in this disorder^45,46^. Blinded reviewers assessed the structural integrity of sensory afferents and receptor endings immunolabeled against neurofilament protein (NEFH)^37^. We then tested for functional evidence of degeneration by comparing differences in conduction delays, a physical measure of nerve degeneration that is used clinically^26^. Neither histological observation of sensory afferents and receptors (Fig. 5d) nor axon conduction tests (Extended Data Fig. 15) revealed evidence of dying-back degeneration of sensory nerve terminals (*Apc*^*Pirc/+*^+OX: 1.41±0.13ms, n=10. *Apc*^*WT*^+control: 1.52±0.14ms, n=11, Extended Data Fig. 15). Our findings demonstrate that transcriptional changes we observed for some markers of nerve degeneration (Fig. 3) were insufficient to yield nerve degeneration. While our results indicate selective resistance of mechanosensory neurons in *Apc*^*Pirc/+*^+OX rats, they are in line with the inconsistency of physical evidence for nerve degeneration reported in clinical studies.^26^ Furthermore, superimposed vibration immediately restores portions of static signaling, which further refutes the existence of degeneration’s role in signaling disorders (Extended Data Fig. 15). Overall, our findings suggest that the structure and core ability of mechanosensory neurons to produce action potentials remain unimpaired by codependent neuropathy and does not depend on dying-back degeneration.

### Dysfunction depends cancer-OX interaction

Data presented in Figure 5 suggests damaging cancer-chemotherapy interactions are conserved *in vivo* and have consequences in sensory function. In order to assess the extent of interaction quantitatively, we took an unbiased statistical approach by subjecting all neurons (n=60, 240 total trials) from all experimental groups to a machine learning algorithm (linear discriminant (LD) analysis). Our approach reduced complex feature space into canonical variables giving us a high-level understanding of where interaction emerges, without biased feature selection *a priori*. Our analysis yielded three canonical variables (Fig. 6a) that achieved overall 94.7% classification accuracy (Extended Data Fig. 17a). We then visualized neuronal signaling in the new 3D composite space created by LD1-3 (Fig. 6a). By projecting high dimensional parameter and feature data onto a simplified 3D canonical space, statistically significant non-linear interaction between cancer and chemotherapy clearly emerged in the first dimension (LD1; 54.23% proportion of variance; Fig. 6a and Extended Data Fig. 17b). Notably, dysfunction in LD1 induced by the cancer—chemotherapy interaction occurred in the opposite direction to that predicted by independent effects, and its magnitude was amplified greater than their sum (Fig. 6a and Extended Data Fig. 17b, c). To test the statistical significance of the cancer—chemotherapy interaction, we conducted Bayesian model comparison with full factorial and all restricted models using leave-one-out cross validation. We then quantified and validated each model’s predictive performance by computing the expected log predictive densities (ELPD; measure of a model’s out-of-sample predictive accuracy in Fig. 6b-e)^47^. We found decisive evidence in favor of a model including a cancer–chemotherapy interaction predictor (ELPD diff ≥ 48 SE <8.1; Fig. 6e). Moreover, our data and generative modeling conclude that codependent interaction (Fig. 6b-d) is necessary to accurately and reliability reproduce clinically relevant neuronal signaling deficits observed in *Apc*^*Pirc/+*^+OX rats (Fig. 6e)

We then asked which functional clusters are highly enriched in LD1. By examining the discriminant function coefficients, we found that static signaling and sensitivity functional clusters represent a large portion of the explained variance in LD1. These findings suggest the sensitivity and static signaling clusters are most susceptible to codependent neuropathy (Extended Data Fig. 17d).

While dynamic signaling provided relatively modest contributions to LD1 (lower coefficient weights in Extended Data Fig. 17d), examining the parameter level data allowed us to discover the expression of distinct classes of interactions. Independently, OX and *Apc*^*Pirc/+*^ induce highly conserved (87.5%) opposing effects exclusively within the dynamic functional cluster. We consistently found that *Apc*^*Pirc/+*^ mutation increased and OX treatment decreased dynamic firing. This led us to predict that their combination would nullify dysfunction and approximate *Apc*^*WT*^+control signaling. Instead, we observed opposing interactions that resulted in drastic reduction in dynamic firing properties. By contrast, for history-dependent functional clusters, we found that cancer–chemotherapy interaction emerged exclusively in *Apc*^*Pirc/+*^+OX neurons, since neither *Apc*^*WT*^+OX nor *Apc*^*Pirc/+*^+control neurons were disrupted.

Taken together, we found that 58% of the neuronal signaling parameters in *Apc*^*Pirc/+*^+OX rats showed high-confidence interaction effects, in that those deficits were greater than those observed in either *Apc*^*WT*^+OX nor *Apc*^*Pirc/+*^+control rats. Our findings demonstrate that complex systemic perturbations of chemotherapy and cancer result in synergistic interactions for sensitivity signaling characteristics (Fig. 6f), opposing interactions for peak firing rate during dynamic signaling (Fig. 6g), and emergent interactions for history dependent and static signaling characteristics (Fig. 6h). In summary, *Apc*^*Pirc/+*^+OX rats express widespread neuronal signaling dysfunction that targets all functional clusters, was independent of structural or physiologically detected degeneration and were not fully accounted for by a global decrease in sensitivity.

## Discussion

Here we present original evidence that cancer transforms the nature and magnitude of neuropathy induced by chemotherapy. Our preclinical study of rats is the first to compare the neuropathic effects of chemotherapy and cancer, both independently and in combination, currently impractical in human study. These comparisons for global transcriptional analysis of sensory neurons in dorsal root ganglia revealed dysregulation of genes uniquely induced, amplified or suppressed by the combination of cancer and chemotherapy. Codependence was conserved as impaired spike encoding of mechanosensory stimuli and novel ion channelopathy. While the present report is limited to rats, extensive conservation of core molecular and cellular processes across mammalian species^48^, leads reasonable expectation that cancer-chemotherapy codependency extends to humans. All considered, we conclude that inattention to co-dependencies necessarily prevents the development of mechanism-based treatments for sensory neuropathy, which remains unexplained and unabated in patients receiving chemotherapy for cancer.

Our transcription analyses expose two potential targets for treating sensorimotor disorders among the debilitating patient symptoms that persist following treatment of various cancers with platinum-based compounds and other antineoplastic agents, e.g. taxanes. Patients display deficits in the spatiotemporal parameters (speed and stride) of walking gait, in control of posture, and in balance relying on proprioception^49,50^. Acknowledging that these disabilities may arise from a variety of lesions in the nervous system, the deficits we observe here in the initial detection and encoding of muscle mechanics would necessarily impair movements and postures and cannot be fully compensated by other senses, e.g. vision. Furthermore, signaling by mechanosensory neurons in rats closely resembles that in human^44^. For these reasons, we assign special attention to depressed signaling by muscle spindles and to the associated decrease in expression of the voltage-gated ion channel Kv3.3 and its gene *Kcnc3*. Reports that *Kcnc3* gene knock out impairs firing responses of neurons and results in ataxia^42^ promote this gene and its ion channel as potential contributors to signaling deficits observed in the present study. At this time, we are uncertain about the sufficiency of Kv3.3 channelopathy to explain deficient mechanosensory signaling in cancer treated by chemotherapy. Uncertainty arises in part, because comprehensive understanding of the molecular mechanisms underlying the function of these sensory neurons is lacking, although advanced by our discovery of Kv3.3 in mechanoreceptors of healthy animals. Another candidate target for treating sensorimotor disorders is the inflammatory signaling molecule IL6, shown here to express large increases in both gene and protein expression. As a result of its effect in suppressing neuronal firing behavior^28^ IL6 has the potential to explain decreased signaling by mechanosensory neurons. We identify both Kv3.3 and IL6, therefore, as targets worthy of testing for their potential value in treating movement disorders.

Our study, being restricted to a single time point following chemotherapy, does not assess the stability of potential treatment targets that emerge from codependent neuropathy. It is reasonable to expect, however, that the codependence undergoes dynamic change. Cancer, presumably also its associated systemic effects, undergoes complex progression as subpopulations of cancer cells differentially resist, adapt, or succumb to chemotherapy. Moreover, biological systems themselves initiate dynamic responses to perturbations. In the nervous system, homeostatic regulation initiates compensatory mechanisms to offset perturbations in neuronal excitability^39^. It is thought provoking in this regard to consider the possibility that the chronic hypoexcitability which follows the acute hyperexcitability^2,45^ that develops during chemotherapy reflects the actions of a dysregulated compensatory mechanism. Understanding these non-linear interactions processes and their effects on potential treatment are both challenging and necessary for developing effective cancer treatment.

## Acknowledgements

We thank staff of the Physiological Research Laboratory at Georgia Institute of Technology. We are grateful to Dr. Thomas Burkholder for technical assistance, discussions and comments. We thank Dr. Mengnan Zhang and Evan Clayton for discussions and comments. This work is supported by NIH grant R01CA221363.

## Author contributions

T.C.C and S.N.H. designed this study. T.C.C and J.F.M. directed and coordinated. S.N.H., P.N., D.C., L.M., E.P. performed the experiments. S.N.H. and L.M. performed the bioinformatics analysis. S.N.H., T.C.C wrote the manuscript.

## Online Methods

### Animal Care

All procedures and experiments were approved by the Georgia Institute of Technology’s Institutional Animal Care and Use Committee. Adult rats (250-350g) were studied in terminal experiments only and were not subject to any other experimental procedures. All animals were housed in clean cages and provided food and water ad libitum in a temperature- and light-controlled environment in Georgia Institute of Technology’s Animal facility.

### Rat model of Colorectal Cancer

We adopted the rat model of colorectal cancer developed in Fisher 344 (F344) rats and known as polyposis in rat colon (*Apc*^*Pirc/+*^)^1^. Pirc rats generated from a germline mutation in the Apc gene (Apcam1137/+) approximate human colorectal cancer. *Apc*^*Pirc/+*^ rats exhibit: regional distribution of tumors that best represents that seen in human colorectal cancer^2^; histopathology and morphology closely resembling human tumors (adenocarcinomas)^3, 4^, up-regulation of well-established pro-growth factors, e.g. β-catenin and EGFR and proliferative makers Ki-67 seen in human carcinogenesis^1^, dramatic increases in Wnt signaling believed to be a key factor in human colorectal carcinogenesis^5^, presence of apoptotic resistant cells unique to carcinogenesis^1, 2^, oxaliplatin (OX)-induction of apoptosis in colon tumors, survivability (12-15 months) enabling long-term investigation of OX’s chronic effects^1^.

### Chemotherapy treatment

OX was injected i.p. once a week (10mg/Kg, 1 ml 5% dextrose in DMSO) to achieve a cumulative dose of 70 mg/Kg over 7 weeks, which scales to a human dose of 420mg/m2 (conversion based on rat body surface area and Km= 6)^6, 7^. This dose minimizes nerve degeneration in patients^8, 9^. The VC groups received i.p. injection of vehicle (1 ml 5% dextrose in DMSO). Body weight was measured 2x/week, which, combined with regular food monitoring, provided early detection of cancer related cachexia^10^. Throughout treatment, rats were frequently monitored for pain or distress. No individual rat reached set criteria established for early removal from the study, e.g. 20% weight loss, vocalization, failure to drink or groom, severe lethargy, self-mutilation, uncontrollable infection.

### Animal Groups

To determine the independent and combinatorial influence of cancer and OX on neuronal dysfunction, we included four groups of animals in all experiments based on the F344 genetic background (*Apc*^*WT*^). Age matched (4 months, tumors detected at 2 months of age) F344 rats were randomly assigned to vehicle control (*Apc*^*WT*^+control) and OX treatment groups (*Apc*^*WT*^+OX). Age matched, male Pirc rats were randomly assigned to VC (*Apc*^*Pirc/+*^+control) or OX treatment (*Apc*^*Pirc/+*^+OX) at 4 months of age when fully developed cancer is present^2^.

### Surgical Procedures

Terminal in vivo experiments were performed 5 weeks after achieving clinically relevant chemotherapy doses, 12 weeks total. They were designed to measure the firing of individual intact sensory neurons in response to physiologically relevant muscle contraction and stretch with electrophysiological techniques. All in vivo procedures are well established in our lab and have been extensively described in previous publications^11-15^; however, brief descriptions are provided. Rats were deeply anesthetized by inhalation of isoflurane (5% in 100% O_2_), intubated via a tracheal cannula, then maintained for the remainder of the experiment (up to 12 hours by 1.5–2.5% in 100% O2). Respiratory rate, pCO_2_, core temperature, pulse rate and pO_2_ were continuously monitored and maintained by adjusting anesthesia^16^ and adjusting heat sources. Dorsal roots (lumbar L4–6), muscles, and nerves in the left hindlimb were prepared for stimulation and/or recording with the rat fixed in a rigid frame at the snout, vertebral bodies, distal tibia, and distal femur (knee angle 120°). Triceps surae muscles (lateral and medial gastrocnemii and soleus) were partially freed of surrounding connective tissue and marked for their resting length (L_o_) at ankle angle 90° before their common Achilles tendon was severed at the calcaneus and tied directly to the lever arm of a force- and length-sensing servomotor (model 305B-LR; Aurora Scientific). Triceps surae nerves were loosely positioned in continuity on a unipolar silver stimulating electrode, and all other hindlimb nerves including common peroneal, sural, and posterior tibial nerves were crushed to reduce extraneous neuronal activity. Dorsal rootlets were carefully freed in continuity from overlying connective tissue and supported on bipolar hook electrodes positioned close to the rootlet’s entry into the dorsal spinal cord. Exposed tissues were covered with warm mineral oil in pools formed by attaching the edges of severed skin to the recording frame.

### *In vivo* Intracellular Recording

Dorsal rootlets positioned in continuity on bipolar recording electrodes were selected for sampling sensory neurons when they produced robust action potential activity in response to both electrical stimulation of triceps surae nerves and stretch of triceps surae muscles. Individual axons penetrated in these rootlets by glass micropipettes (∼30 MΩ filled with 2 M K+ acetate) were selected for study when electrical stimulation of triceps surae nerves produced orthodromic action potentials that were readily resolvable and had conduction delay of <2ms. Continuous intracellular recordings from sensory neurons we acquired with Spike2 software (version 8.02). Neurons were first classified on the basis of their responses to stimuli, e.g., muscle twitch contraction and vibration, and second to characterize the afferents’ sensory encoding of a range of (naturalistic) physiologically meaningful mechanical events. Sensory neurons were classified on the basis of binary scoring of three criteria previously described by our lab^15^ and in the cat^17^. Sensory neurons were classified as Ia by their perfect entrainment to 1-s bouts of high-frequency, small-amplitude vibration (100Hz, 80 µm), pause in firing during rising twitch force response (muscle shortening), and by responding with an initial burst of high-frequency firing (>100 pulses per second (pps)) at the onset of muscle stretch. Two stretch paradigms were used to characterize the firing responses of afferents to physiologically relevant mechanical stimuli. Stretches were delivered by the servomotor to triceps surae muscles, which were not engaged in active contraction. In both paradigms, triceps surae muscles were stretched by 3 mm from L_o_(7% strain) (Dashed vertical line marks the onset of muscle stretch Fig. 5 and Extended Data Figure 13-16). Ramp-hold-release stretches tested afferent encoding of both fast dynamic (20mm/s, 47% strain rate) and static stimuli; successive triplets of triangular stretch tested slow (4mm/s, 9% strain rate) dynamic and activity-dependent encoding in dynamic stretch known to be influenced by recent signaling history^12^. Strains and strain-rates fall within values expected for animals engaged in normal activities e.g. locomotion^18^, and have been previously used in our lab and others^12, 15, 17, 19^.

### Tissue Collection

At the conclusion of data collection, rats were overdosed with isoflurane inhalation (5%) then immediately transcardially perfused with cold vascular rinse (0.01 M phosphate buffer with 0.8% NaCl, 0.025% KCl, and 0.05% NaHCO3, pH 7.4) followed by room temperature fixative (2% paraformaldehyde in 0.1 M phosphate buffer, pH 7.4). Muscles and dorsal root ganglia were quickly dissected and post-fixed for one hour in the same fixative at room temperature. After a brief wash in a 0.1 M PBS, tissues were incubated in 0.1 M PBS containing 20% sucrose at 4°C overnight for cryoprotection. Three to four animals from each of four groups were selected for whole-transcriptome analysis alone and did not undergo further analysis. Surgical preparation to expose the lumbar spinal cord took place as described above. Dorsal root ganglia (DRG) from the lumbosacral (L5-S2) spinal cord were surgically removed (4-6 per animal), washed with sterile saline and immediately flash frozen (fresh) in 2-methylbutane (isopentane) pre-chilled in liquid nitrogen. DRG were then stored at −80°C for further analysis (see below). After DRG removal, colonic tumors from *Apc*^*Pirc/+*^ *and Apc*^*Pirc/+*^ +OX rats were dissected and counted prior to post fixation for immunohistochemical analysis

### Immunohistochemistry and Imaging

Detail immunohistochemistry procedures have been previously described^20^. Briefly, 50-µm thick sections of skeletal muscles, DRGs, and tumors were cut using a Cryostat (Leica). Tissues sections from all treatment groups were processed simultaneously. All tissue sections were incubated overnight in primary antibodies diluted in blocking buffer (5% normal goat serum, 0.3% Triton100 in PBS). The primary antibodies used were as follow: rabbit polyclonal anti-NaV1.6 (ASC-009, Alomone Laboratories, 1:200), rabbit polyclonal anti-NaV1.1 (ASC-001, Alomone Laboratories, 1:300), rabbit polyclonal anti-NaV1.7 (ASC-008, Alomone Laboratories,1:100), rabbit polyclonal anti-Kv3.3 (APC-102, Alomone Laboratories, 1:300), mouse monoclonal anti-Kv1.1 (K36/, NeuroMab, University of California Davis, 1:300), guinea pig polyclonal anti-VGLUT1 (135-104 Synaptic Systems, 1:300), chicken polyclonal anti-neurofilament protein (NF-H, Aves Laboratories, 1:200), mouse monoclonal anti-ASIC2, (E-20, Santa Cruz Biotechnologies,1:200), mouse monoclonal anti-EGRF (SC-120, Santa Cruz Biotechnolgy, 1:100), mouse monoclonal anti-KI67 (SC-23900, Santa Cruz Biotechnology, 1:100), mouse monoclonal anti-IL6 (SC-57315, Santa Cruz Biotechnology, 1:100). After washing with PBS, tissue sections were incubated with appropriate fluorescent-conjugated secondary antibodies (Jackson ImmunoResearch Laboratories), diluted in blocking buffer, for 1 hr. at room temperature. Following washes with PBS, slides were mounted using Vectashield containing DAPI (Vector Laboratories) in order to label cell nuclei.

### RNA Extraction and Amplification

RNA extraction and amplification were performed according to our previously described methods^21^. Briefly, RNAs were isolated and purified from sensory neurons of DRG (L5-S2) using the mRNeasy Micro KIT (Qiagen, Germantown, MD). Total RNA concentration and integrity was assessed on the Bioanalyzer RNA Pico Chip (Agilent Technologies, Santa Clara, CA). Labeling of RNAs was performed with the FlashTag Biotin HSR RNA Labeling Kit (Affymetrix, ThermoFisher). Gene expression was determined by microarray analysis on the Affymetrix platform (Rat Transcriptome Array 2.0).

### Microarray data analysis

In total, 13 (four groups with biologic triplicates per group (4 biologic replicates from *Apc*^*Pirc/+*^+OX group) global transcriptional expression data sets were generated in this study. Raw mRNA expression data were processed using Affymetrix Expression Console (EC) software Version 1.4. Raw data probes were normalized using SST-RMA algorithm. Differentially expressed genes (DEG) were determined through a linear fixed effects model (*limma*^*22, 23*^) with a robust empirical Bayes (eBayes) framework^24^ to moderate the residual variances (1.5 fold change and FDR <1% for multiple test correction across contrasts) *22, 23*. This has the effect of increasing the effective degrees of freedom by which gene-wise variances are estimated. This approach reduces the number of false positives for genes with small variances and improves power to detect DEG with larger variances^25^. In effect, this technique allows the borrowing of information across genes to gain statistical power.

### Unsupervised multivariate analysis

DEGs were subjected to paired multivariate analysis to discover high-level data structure. By serially linking two powerful analytic techniques, Principal Components Analysis (PCA) driven Hierarchical clustering, we stabilize clustering performance and reduced dimensionality to tractably few continuous variables containing the most important information on which clustering algorithms are focused which leads to a better clustering solution^26, 27^. PCA and visualization was applied to DEGs with the *factoextra*^*28*^ and *FactoMineR*^*26*^ packages. PCs capturing high-level structure were then employed to drive hierarchical clustering. Hierarchical clustering and heatmaps were visualization of the data were generated with *heatmap.2* command within the *gplots* package^29^ in the R environment (3.5.0)^30^. Hierarchical-clustering, using the ward.D2 method^31^ were applied to the eBayes-filtered gene set to obtain unsupervised visualization of the gene clusters coordinately expressed among the different experimental groups. Prior to hierarchical clustering, genes were de-meaned and standardized across the samples to a mean of zero and an s.d. of 1; *z* values were then used for clustering. Subclusters were derived using the R function *cutree*^*32*^.

### Pathway and Gene Set Enrichment Analysis

Hypergeometric testing was conducted on subclusters of named genes using DAVID (Database for Annotation, Visualization, and Integrated Discovery) and g:Profiler to pinpoint significantly enriched pathways or gene ontology terms^33, 34^. We increase statistical power by hypergeometric testing on subclusters which separates co-regulated (up or down) probe sets^35^. All biological processes with adjusted P < 0.05 were considered significantly enriched. g:Profiler 3-500 (min-max) Benjamini Hochberg FDR correction intersection terms (2) only annotated genes were searched.

Parallel identification of pathways significantly enriched in *Apc*^*WT*^+OX, *Apc*^*Pirc/+*^+control, *and Apc*^*Pirc/+*^+OX sensory neurons, as compared with *Apc*^*WT*^+control neurons, was performed by GSEA (gene-set enrichment analysis). GSEA enables us to uncover novel sets of functionally related genes by focusing on all detected (above-background) genes. GSEA is unbiased and more sensitive, thus allowing for the detection of even subtle enrichment signals. GSEA was also used to compare our gene-expression profiles with those of previously published, related studies. To this end, we used the Molecular Signature Database v 6.2 (C2-CP: canonical pathways; C3-MIR: microRNA targets; C3-TFT: transcription factor targets; C5-BP: GO biological process; C5-CC: GO cellular component; C5-MF: GO molecular function C7: immunologic signatures gene sets) with the permutation type set to ‘gene set,’ 15-500 (min-max), to calculate statistical significance, as suggested for fewer than seven replicates; default settings were applied to all the other options. GSEA contrasts were visualized as networks (enrichment maps) using the EnrichmentMap (version 3.1.0)^36^ plugin in the Cytoscape software (version 3.7.0)^37^. For GSEA, a false discovery rate (FDR) of <0.2 was considered statistically significant.

### Targeted Transcriptional Analysis

Complementary supervised analysis of DEGs leveraged knowledge of proteins known to be expressed at sensory neuron receptor endings that directly^20, 38, 39^ or indirectly^40, 41^ regulate excitability. In addition, due to incomplete knowledge of the full suite of proteins that are necessary and sufficient for normal function^41^, we expand targeted transcriptional analysis to incorporate gene families, including: ion channels, GPCRs, and channel regulators. DEGs and gene families were illustrated using volcano plots, generated by plotting fold-change differences against comparison −log_10_P-values.

### 3D Digital Reconstructions and Anatomical Quantifications

Z-axis stacks of images of muscle spindles receptor endings, DRGs and tumors were constructed by sequentially imaging using a confocal microscope (LSM 700A, Zeiss). Muscle spindles terminals z-stacks (1μm steps) were captured with a 40X oil immersion objective (N.A 0.6) at 0.6 digital zoom. DRG and tumors z-stacks (1μm steps) were captured with a 20X objective (N.A 0.4) at 0.6 digital zoom. Stacks of images were processed and analyzed using Amaris (Bitplane) or imageJ (NIH) imaging software. Figures present images as flat maximal projections of the sum of the z-axis optical slices. Analysis of neuron receptor endings (spindles) and soma (DRG) were performed by an investigator blinded to treatment group. Kv3.3 immunoreactivity in muscle spindle nerve terminals was determined by measuring the mean pixel intensity of Kv3.3 staining on each en-face portions of the terminal (visualized by the VGLUT1 staining) observed on every 1 um single image of the Z-stack. Thus, Kv3.3 mean pixel intensity for each muscle spindle is equal to the product of the sum of mean pixel intensity measured on each portion of the terminal divided by the total number of nerve terminal portions analyzed. Kv3.3 immunoreactivity in DRG was determined by measuring the mean pixel intensity of Kv3.3 staining on the medium and large size cell somas observed on the flat maximal projections of the z-stacks images.

### Intracellular Recording Pre-Processing

Intra-axonal recordings of action potentials together with records of muscle length and force were digitized (20 kHz) and were monitored online and stored on computer for later analysis with Spike2 and custom-written MATLAB scripts. From raw intracellular data of both paradigms, we extracted 31 measured and derived features (Extended Data Fig. 13) that provide a comprehensive quantification of neuronal signaling characteristics. We then categorized features into four broad clusters that represent functional features encoded by these neurons, containing sensitivity (Thr), dynamic (Dyn), static (Stat), and history-dependent (Hx) signaling information. Extended Data Fig. 13 identifies and describes the measured and computed features and functional clustering used for inference.

### Statistical Analysis of Intracellular Recordings

Linear discriminant analysis (LDA) was used to provide supervised dimensionality reduction for the multiple features of neuron signaling, in an attempt to find a linear combination of features that separated and characterized independent and combinatorial treatment effects. Data were first de-meaned, normalized to unit standard deviation, and tested by Bayesian one-way ANOVA (*stan_glm*)^42^. The derived covariance matrix was then normalized by within-group, pooled covariance. The eigenvectors of that modified covariance matrix defined three canonical variables that characterized and separated the four treatment groups identified *a priori* as *Apc*^*WT*^+control, *Apc*^*WT*^+OX, *Apc*^*Pirc/+*^+control, *and Apc*^*Pirc/+*^+OX. LDA and 10-fold cross validation of model performance (repeated holdout method) was performed with the *MASS* (7.3-51.1)^43^ library in the R environment (3.5.0)^30^.

Individual features characterizing the firing responses of neurons sampled from multiple rats were tested for statistically significant differences with Bayesian one-way ANOVA (*stan_glm*). Bayesian parameters estimation derives the entire joint posterior distribution of all parameters simultaneously thus circumventing the need to correct for multiple tests on the data^44, 45^.

Bayesian hypothesis testing was performed by analyzing the probability mass of the parameter region in question (estimated by the number of samples drawn from the posterior that fall in this region). This leads to a direct probability measure that defines values inside the 95% of the HDI more credible than outside values^46^. HDI was used to make unbiased decisions on parameter values. Typically, inferences are drawn directly from the comparison of posterior probability distributions between two (or more) contrasts of interests e.g. mean-comparisons testing. For example, an HDI of a credible difference distribution that does not span zero indicates that the model predictions for the two conditions of interest are different from each other. This reallocates evidential support in favor of the alternative hypothesis that the parameters for both populations are unequal. Alternatively, if an HDI of a credible difference distribution spans zero, that indicates the model predictions for the two conditions do not differ. By defining a region of practical equivalence (ROPE) range (values between −0.1 and 0.1), Bayesian analytic techniques afford the ability to accept the null hypothesis. In practice, when the 95% HDI falls completely within the ROPE region we declare the ROPE value accepted and determine that group differences are practically equivalent to zero.

### Bayesian Models

All models were developed in fully hierarchal Bayesian framework with the *rstanarm* package (2.18.1)^42^ in the R environment (3.5.0)^30^. *Rstanarm* implements regression models in *stan*^*47*^, which are fit using Hamiltonian Markov Chain Monte Carlo sampling to compute credible parameter values, e.g. means, standard deviations, regression coefficients, effect sizes^48, 49^. For intercepts and predictors we use Student’s *t-* distribution with mean zero and four degrees of freedom as the prior distribution. The scale of the prior distribution is 10 for the intercept and 2.5 for the predictors. Each model was run with four independent chains for 400 warm-up, 4,000 sampling steps and every second sample (thinning) was discarded. For all parameters, the number of effective (*n_eff*) samples was >500. Convergence was assessed and assumed to have reached the stationary distribution by ensuring that the Gelman–Rubin shrinkage statistic (rhat, *R^*) statistic for all reported parameters was <1.05 (and Monte Carlo standard error of the parameter means was <0.002^50^. We report the expected mean parameter values alongside 95% credible intervals using the highest posterior density approach (HDI) (Extended Data Fig. 19)

### Bayesian Model Validation

Models’ performance were further validated (Extended Data Fig. 19) by computing out-of-sample predictive accuracy using Pareto-smoothed importance sampling (PSIS^51^) to perform leave-one-out cross (LOO) validation. Although alternative validation strategies (WAIC^52^) are asymptotically equal to LOO, PSIS-LOO provides a more accurate and stable estimate of model performance and is more robust in the finite cases with diffuse priors (weakly informative). Model validation was performed in *Rstanarm*^42^ to generate novel posterior draws using the following:

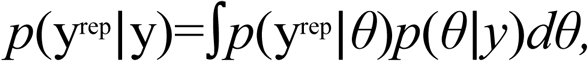

where *y* was observed data and θ were modeled parameters. For each draw of *θ* from the posterior *p*(θ|y) we simulate data *y*^*rep*^ from the posterior predictive distribution *p(y*^*rep*^*|y).* This predictive model simulation generates out-of-sample data (*y*^*rep*^) we can use to make inferences about model performance (Fig. 6). We then obtained standard errors for predictions and for comparison of predictive errors between models to definitively test and rank models based on predictive accuracy (*elpd* = expected log pointwise predictive density)^51^.

### Data availability

Microarray datasets supporting the conclusions of this article are available in the Gene Expression Omnibus (GEO) repository under accession code: GSE126773. Other datasets supporting the conclusions of this article are included within the article and its Supplementary Files or have been deposited in the public repository https://github.com/nickh89/Stephen-N.-Housley-Public-Repository-for-Publication-Code-and-Data

### Code availability

All code and models used can be accessed in the public repository https://github.com/nickh89/Stephen-N.-Housley-Public-Repository-for-Publication-Code-and-Data

### Reporting Summary

Further information on experimental design is available in the Nature Research Reporting Summary linked to this article.

