## Extended Data Figures and Tables for "Cancer Exacerbates Chemotherapy Induced Sensory Neuropathy"

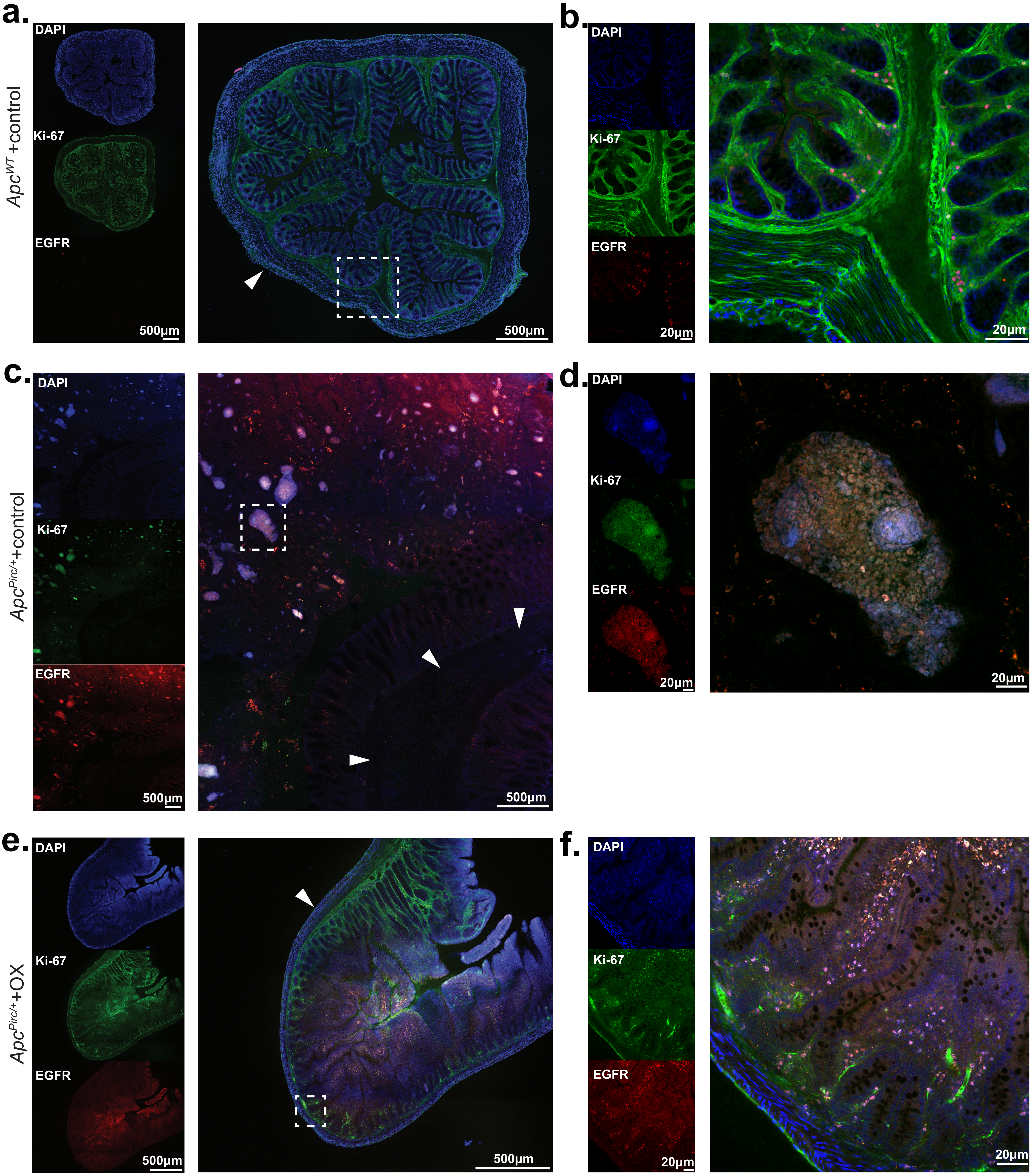

**Extended Data Figure 1. Immunohistochemical and morphological analyses of healthy and tumor bearing colons a-f,** Confocal images of a colon from *Apc*$^{WT}$+control rat (**a, b**) and colon and colon tumors from *Apc*$^{Pirc/+}$+control (**c, d**) and *Apc*$^{Pirc/+}$+OX rats (**e, f**). Immunolabeling shows expression of DAPI and Ki-67, and EGFR a measure proliferative growth associated with tumor progression. White dotted boxes in wide-field images (**a, c, e**) outline region of interest (**b, d, f**). White arrows indicate smooth muscle boundary used as anatomical reference.

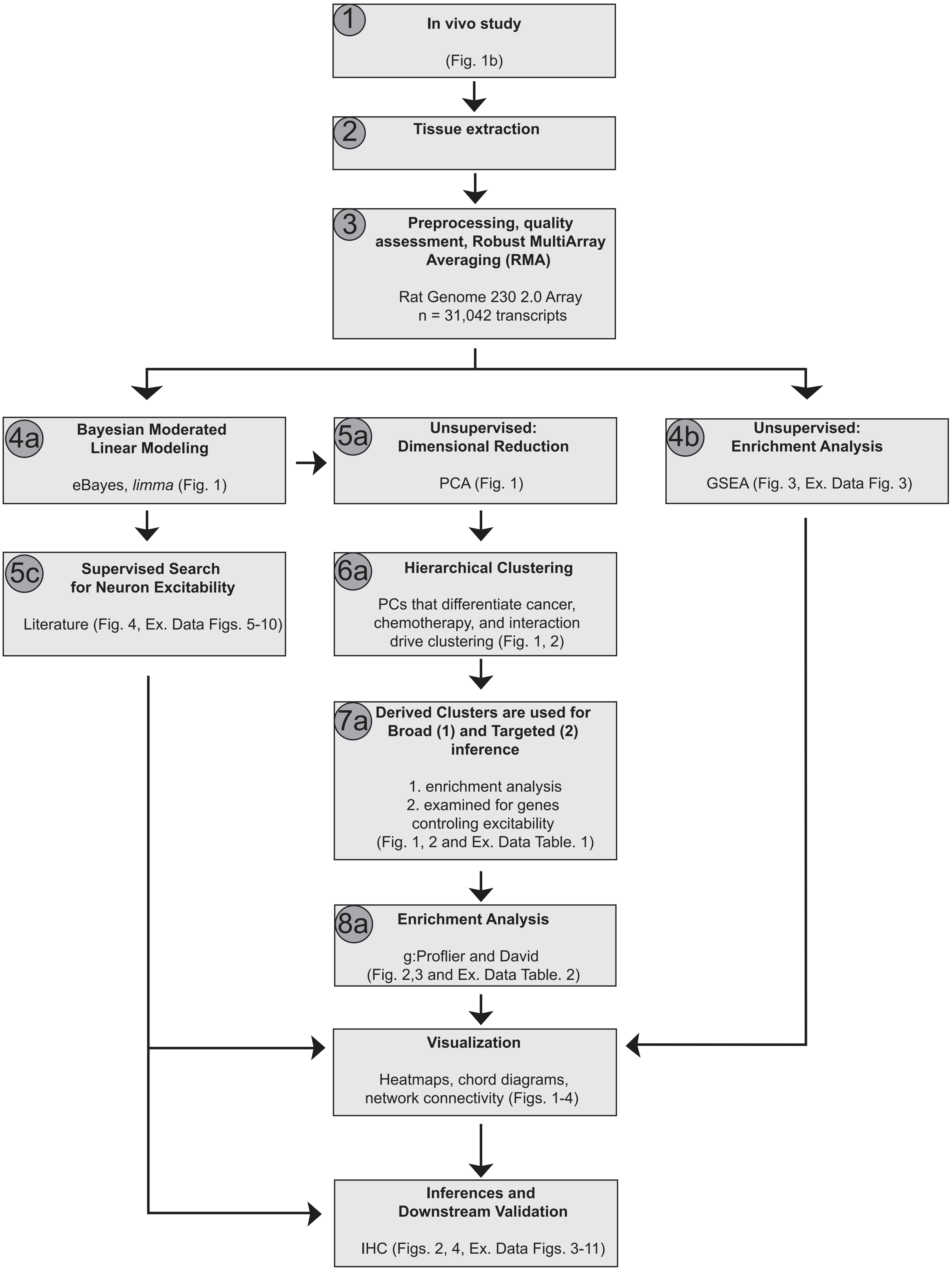

**Extended Data Figure 2. Steps outlining parallel independent transcriptome analyses**, related to Figures 1, 2, 3, 4 and Extended Data Figures 3-11. Data analysis flowchart depicts the multi-step (split into a, b, and c streams) approach utilized for transcriptome analysis, inferences and downstream protein level validation. A detailed description of each step is available in the **Online Methods Sections.** Each box highlights specific analytic techniques, statistical packages (when relevant), and corresponding figures as reference. Principal component analysis, PCA; gene set enrichment analysis, GSEA; principal components, PC; Database for Annotation, Visualization and Integrated Discovery, DAVID; immunohistochemistry, IHC.

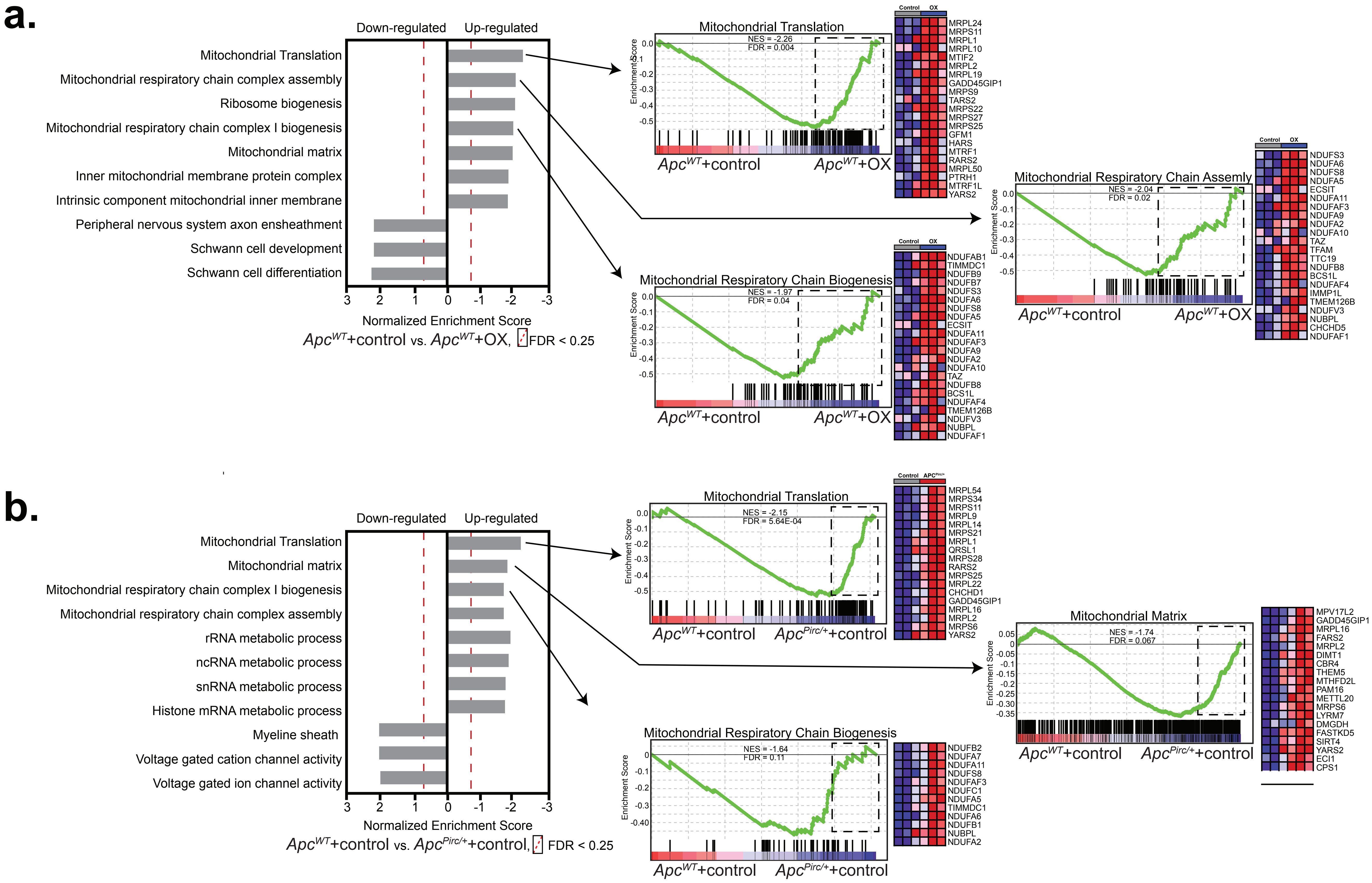

**Extended Data Figure 3. Pathway analysis and GSEA based comparison of the independent effects of cancer and chemotherapy show convergence of mitochondrial dysfunction.** List of enriched gene sets in sensory neurons identified by GSEA (queried against MSigDB, C2-CP: canonical pathways; C3- MIR: microRNA targets; C3- TFT: transcription factor targets; C5-BP: GO biological process; C5- CC: GO cellular component; C5- MF: GO molecular function C7: immunologic signatures gene sets from the comparison of **a,** *Apc*$^{WT}$ +control and *Apc*$^{WT}$+OX and **b,** *Apc*$^{WT}$ +control and *Apc*$^{Pirc/+}$+control groups. Individual frames identify functional relatedgroups of gene sets expressing distinct up- or downregulation as compared with control. Representative enrichment plots are shown on the right. Heat-map representation of leading-edge genes are shown on the right of each enrichment plot highlight key mitochondrion-related proc, such as translation and biogenesis in both *Apc*$^{WT}$+OX and *Apc*$^{Pirc/+}$+control neurons, and key neuron myelinatio-related processes. Statistical significance was determined by permutation testing with normalized enrichment score (NES) and Benjamini–Hochberg false discovery rate (FDR) < 0.25.

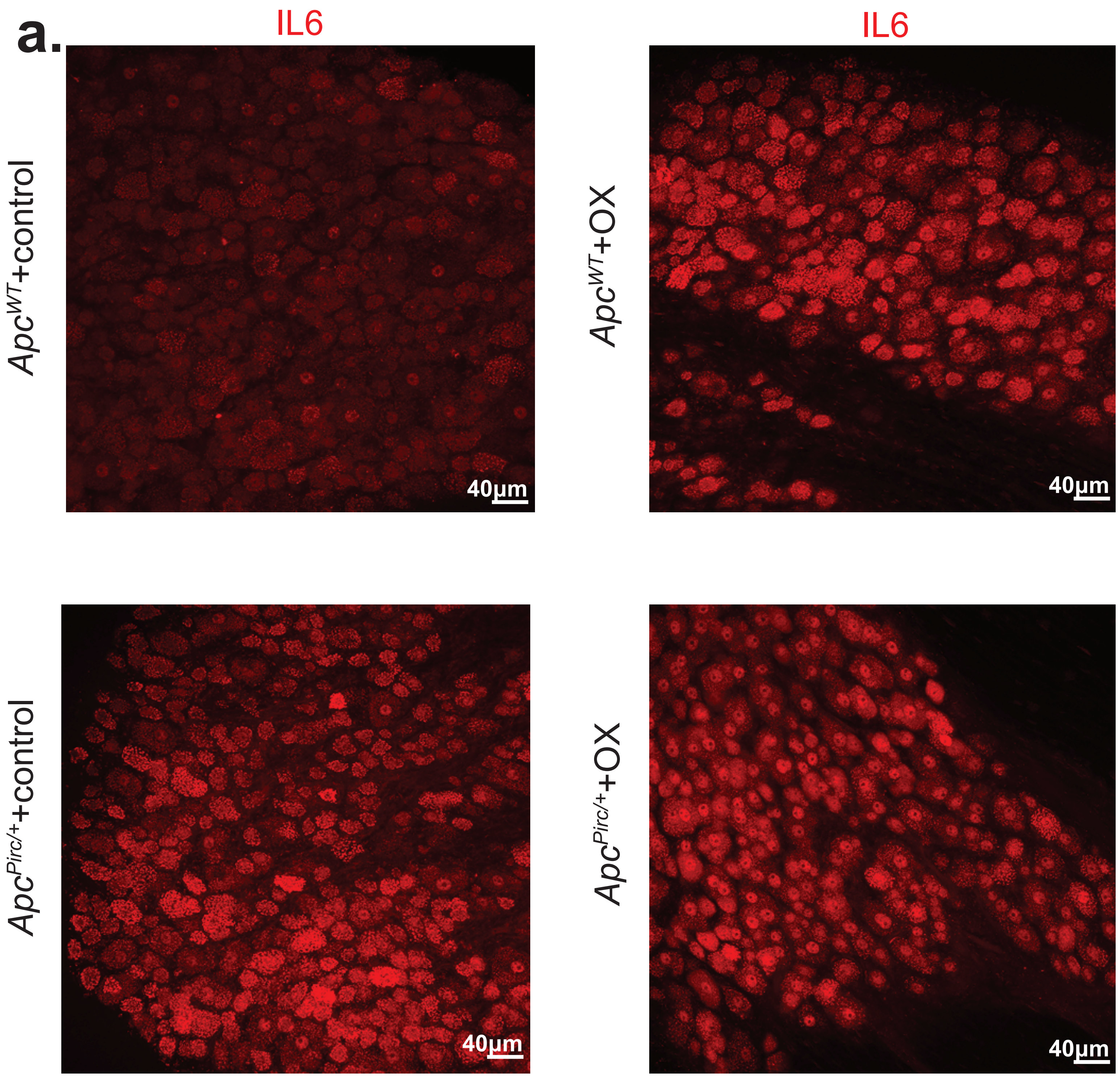

**Extended Data Figure 4. Immunostaining of sensory neurons in dorsal root ganglia (DRG),** related to Fig. 2c and 4c. , Whole-mount staining and confocal axial projections of DRG neurons highlight expression of IL6 (**a**) in *Apc*$^{WT}$+control, *Apc*$^{WT}$+OX, *Apc*$^{Pirc/+}$+control, and *Apc*$^{Pirc/+}$+OX rats

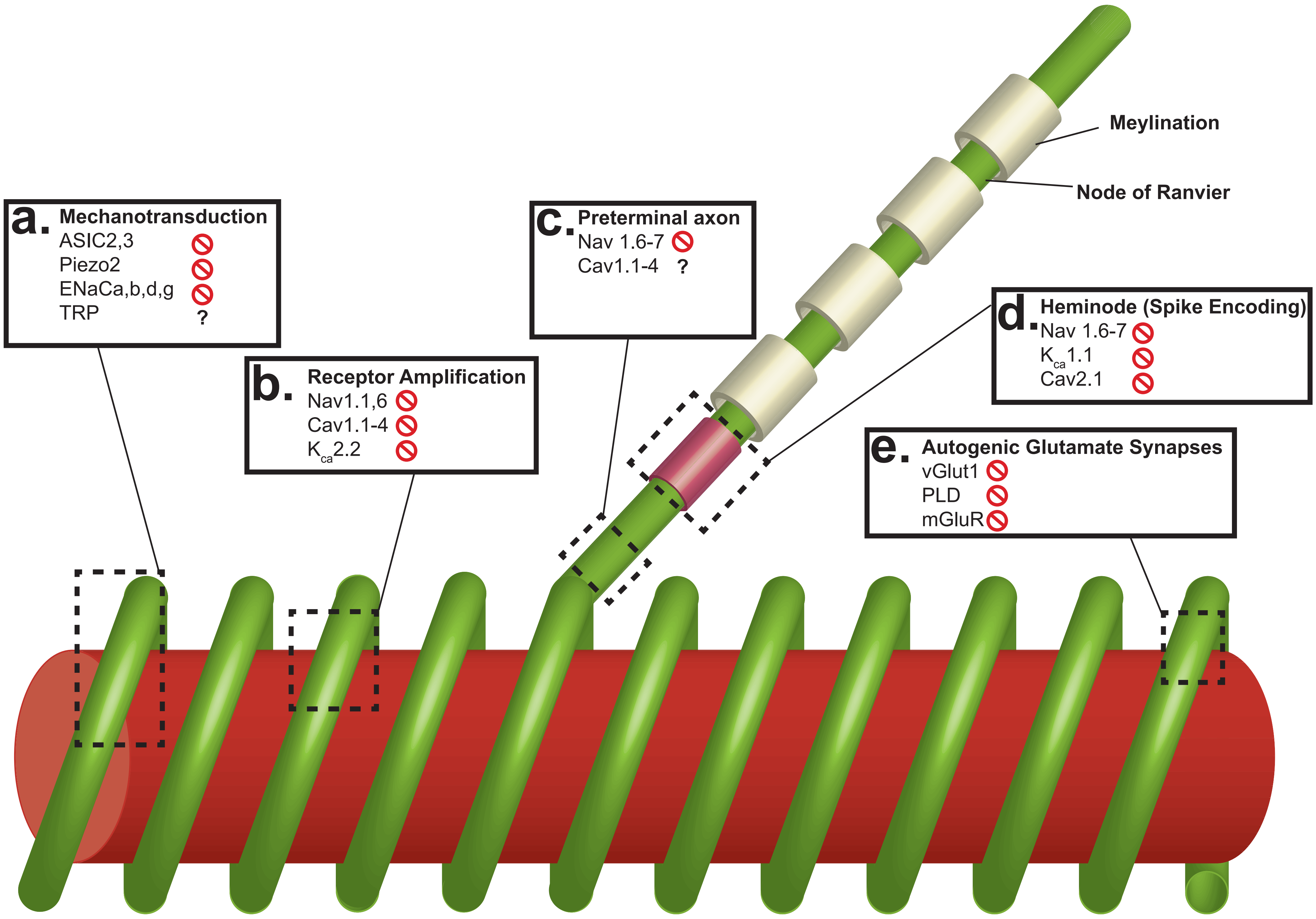

**Extended Data Figure 5. Model of molecular determinants of neuronal signaling of sensory neurons.** Model of mechanically-, voltage- and ligand-gated ion channels and their distribution in muscle spindle Ia primary ending. The intrafusal muscle fiber (red) is wrapped by an annulospiral ending (green) and a preterminal axon that extends unmyelinated from the terminal to the heminode (pink) followed by a myelinated axon. Dotted-boxes represent specific regional distributions believed to underlie specific functional characteristics of neuronal signaling e.g. mechanotransduction, receptor potential amplification, spike encoding, and autogenic feedback. Red circles with slashes indicate no gene and protein level evidential support for involvement in mediating cancer-chemotherapy codepedent neuronal dysfunction. Bold question marks (**?**) indicate protein as a possible candidate due to evidential support from transcriptome, yet protein level confirmation is unavailable. **a,** Deformation of the receptor ending opens mechanotransduction channels that initiate excitatory currents mediated by ASIC2/3 (Simon et al 2010), Piezo2 (Woo et al 2015), ENaCa,b,d,g and to some extend TRP channels (Bewick and Banks 2015). Various Nav (Carrasco et al 2017) and Cav (Bewick and Banks 2015) channels are believed to amplify receptor potentials (**b**) and assist transmission toward encoding in the preterminal axons (**c**). **d**, Recent discovery of specific distribution of Nav channels in the heminodes (Carrasco et al 2017) are believed to underlie encoding of action potential similar to axon initial segments in other systems (Foust et al 2010). **e**, Three components known to regulate autogenic glutamatergic synapses in proprioceptors critical for signaling.

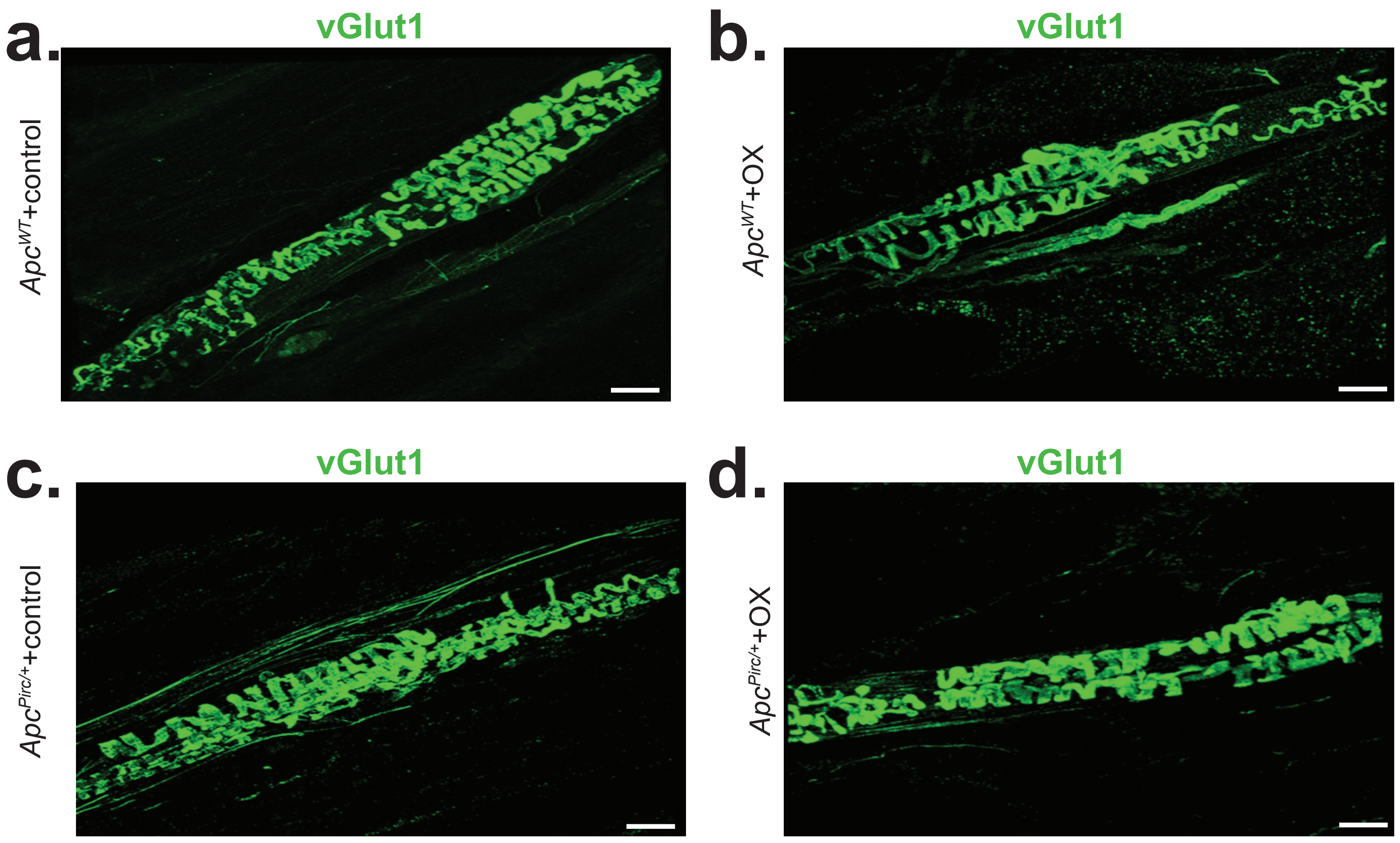

**Extended Data Figure 6. Immunostaining of proprioceptive sensory neurons receptor endings. a-d,** Whole-mount staining and confocal axial projection of the preterminal axons and annulospiral receptor endings of proprioceptive sensory neurons showing single channel images of vGlut1 (green) from: **a,** *Apc*$^{WT}$ +control rat; **b**, *Apc*$^{Pirc/+}$+control; **c,** *Apc*$^{WT}$+OX; **d**, and *Apc*$^{Pirc/+}$+OX rats.

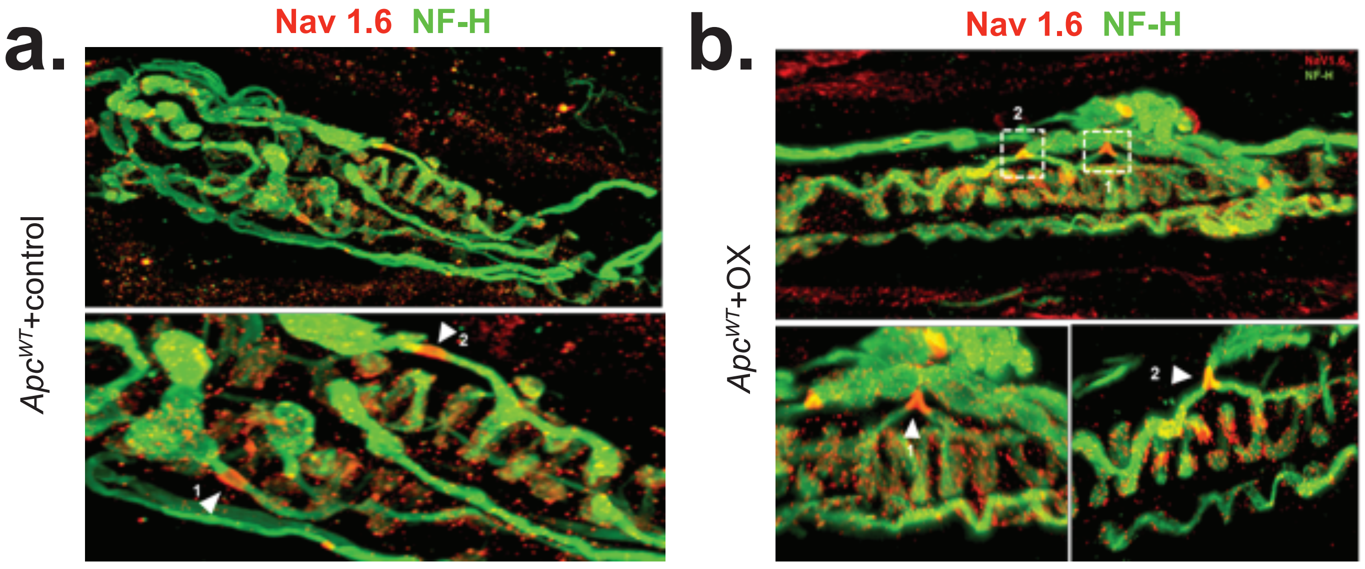

**Extended Data Figure 7. Immunostaining of proprioceptive sensory neurons receptor endings. a-d,** Whole-mount staining and confocal axial projection of the preterminal axons and annulospiral receptor endings of proprioceptive sensory neurons showing merged and single channel images of Nav1.6 (red) and NF-H (green) in top row from: **a,** *Apc*$^{WT}$+control rat; **b**, *Apc*$^{Pirc/+}$+control; **c,** *Apc*$^{WT}$+OX; **d**, and *Apc*$^{Pirc/+}$+OX rats. Bottom rows show Nav1.6 distribution in heminodes where spike encoding is initated.

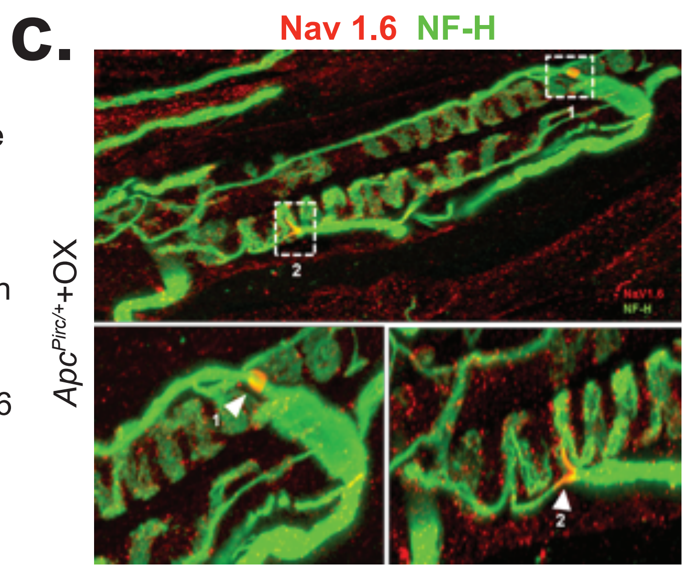

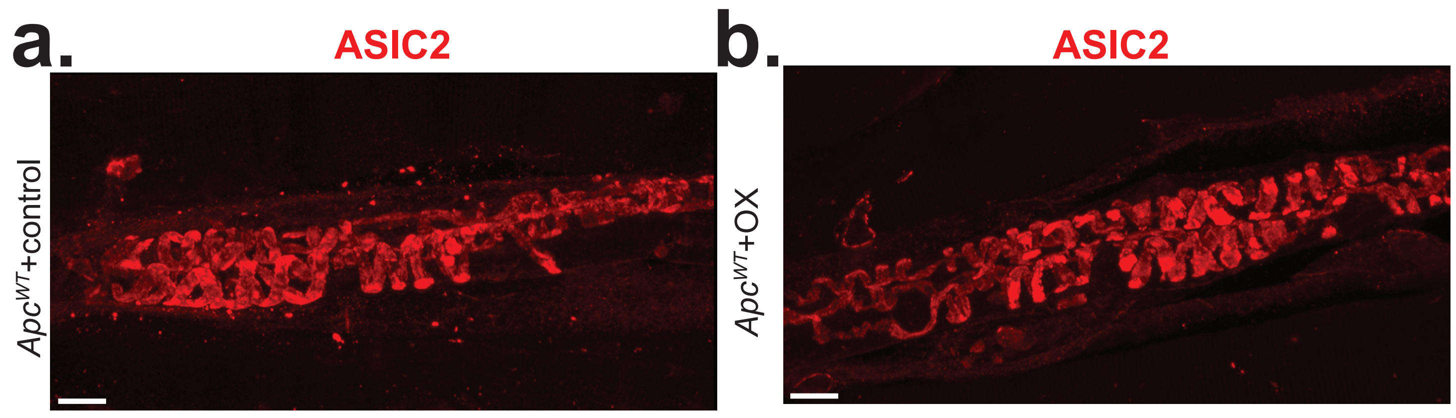

**Extended Data Figure 8. Immunostaining of proprioceptive sensory neurons receptor endings. a-c,** Whole-mount staining and confocal axial projection of the preterminal axons and annulospiral receptor endings of proprioceptive sensory neurons showing merged and single channel images of Asic2 (red) from: **a,** *Apc$^{WT}$* +control rat; **b**, *Apc$^{Pirc/+}$*+control; **c,** *Apc$^{Pirc/+}$*+OX rats.

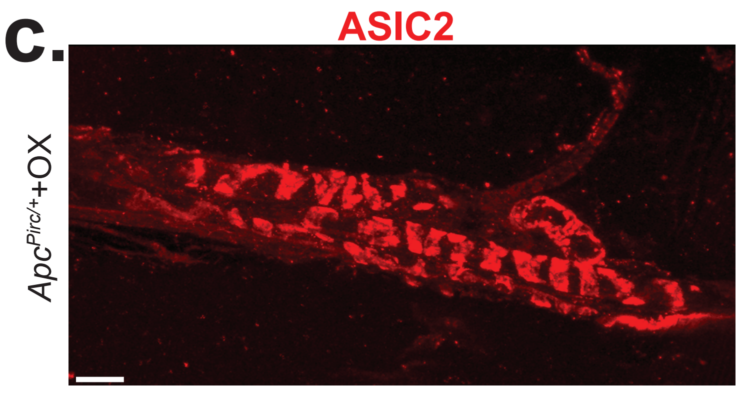

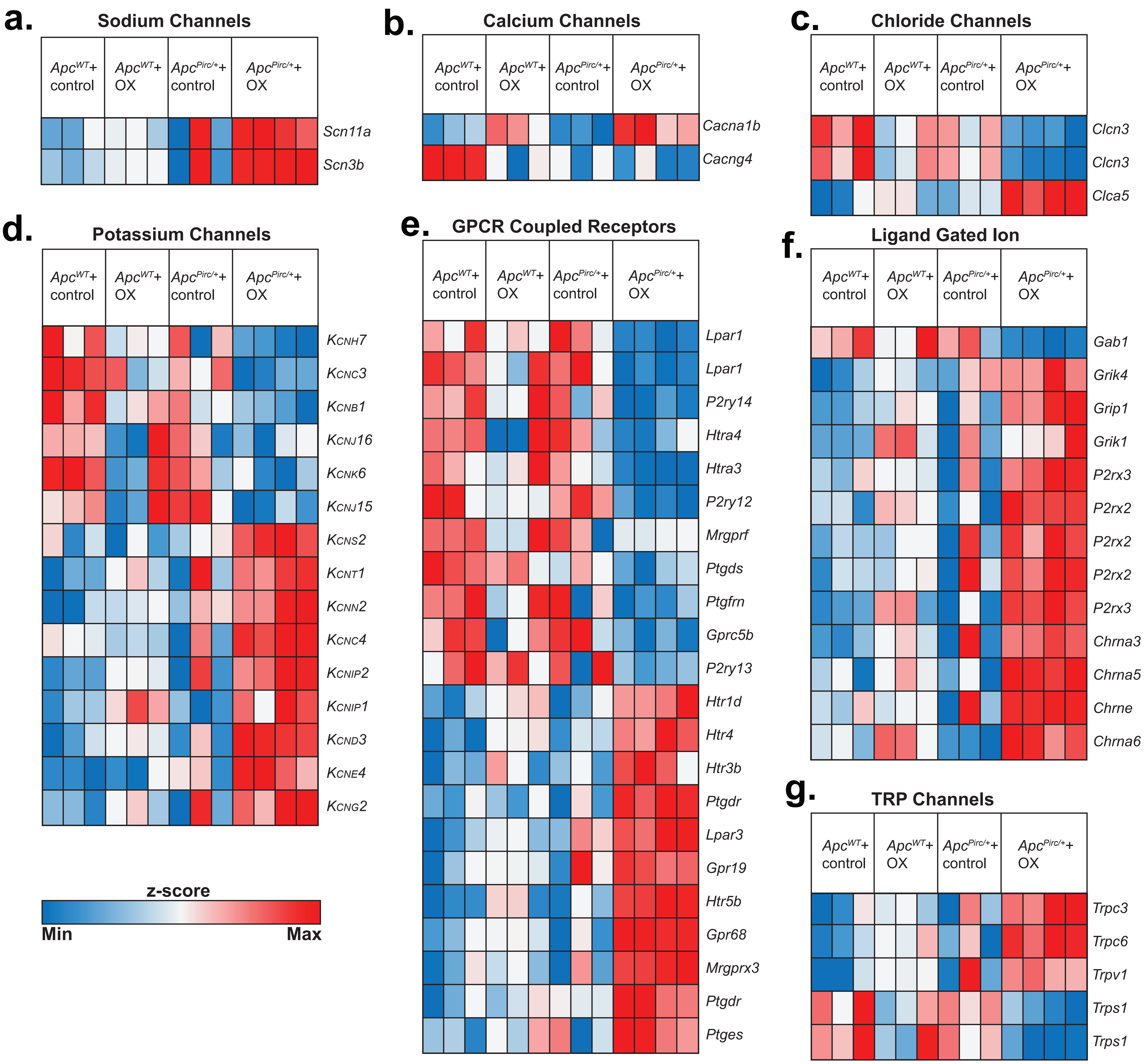

**Extended Data Figure 9. Heat-map distribution of significantly differentially expressed genes specific to neuron neuronal excitability.** Major volatage-gated, mechanically gated, TRP channels and ligand-gated and G-protein coupled receptors across experimental groups. Expression patterns of different sub-types channels and receptors were identified by empirical Bayesian moderated linear fixed effects models (columns are individual samples, heat-maps). (**a**) Sodium channel levels, (**b**) calcium channel levels, (**c**) chloride channel levels (**d**) potassium channel levels, (**e**) G-protein coupled receptors (GPCRs) levels, (**f**) ligand-gated channel levels, and (**g**) transient receptor potential (TRP) channel levels are plotted as heat-maps. Data row standardized (mean centered and s.d. normalized) to specific genes.

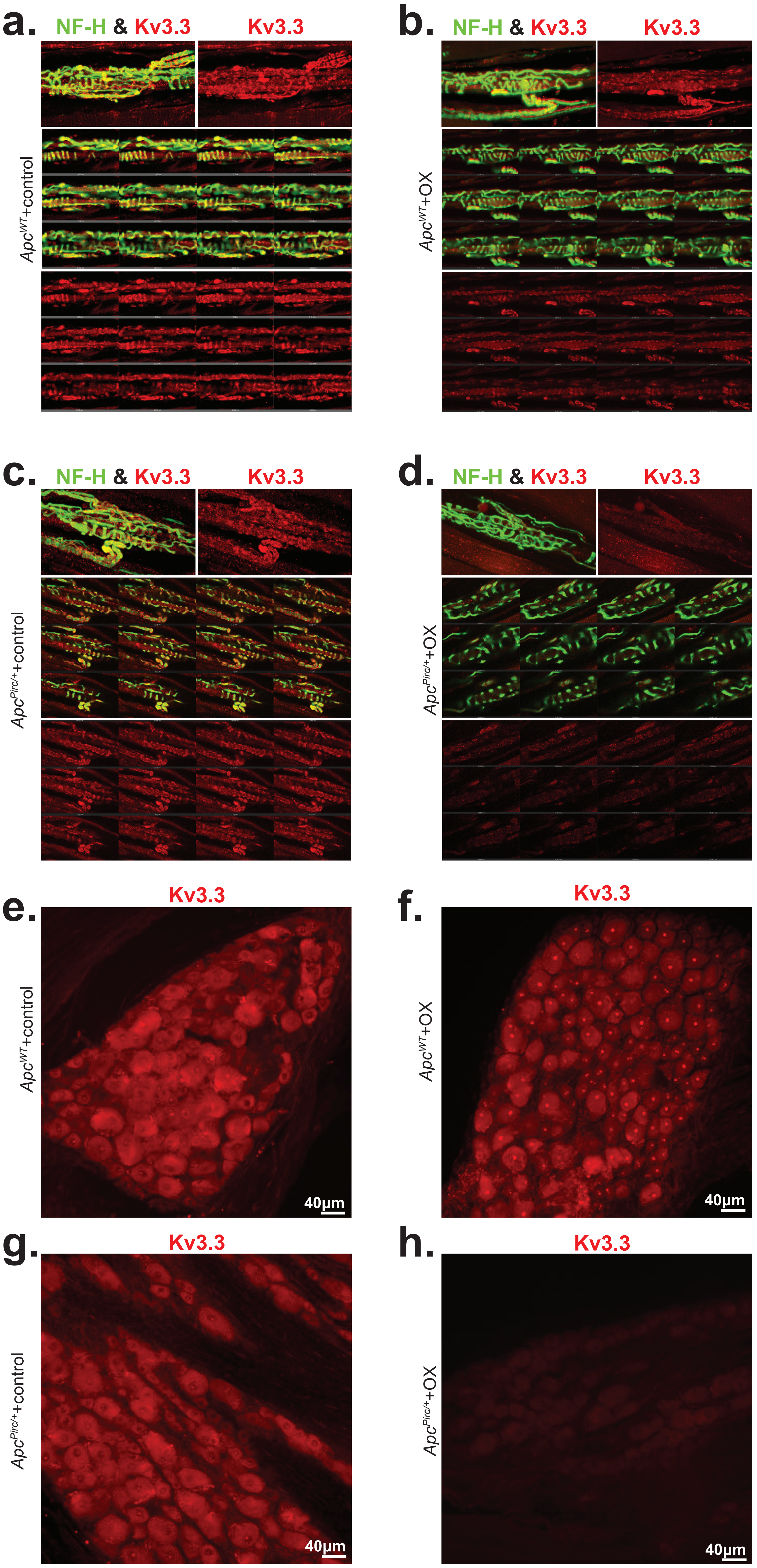

**Extended Data Figure 10. Immunostaining of proprioceptive sensory neurons receptor endings and cell bodies. a-d,** Whole-mount staining and confocal axial projection of the preterminal axons and annulospiral receptor endings of proprioceptive sensory neurons showing merged and single channel images of Kv3.3 (red) and NF-H (green) in top row from: **a,** *Apc*$^{WT}$ +control rat; **b**, *Apc*$^{Pirc/+}$+control; **c,** *Apc*$^{WT}$+OX; **d**, and *Apc*$^{Pirc/+}$+OX rats. Individual optical planes (from z-stack projection) of axial projection highight fine three-dimensional structural distribution of merged and single channel images of Kv3.3 (bottom). Whole-mount staining and confocal axial projections of DRG neurons highlight expression of Kv3.3 in **e**, *Apc*$^{WT}$ +control rat; **f**, *Apc*$^{Pirc/+}$+control; **g**, *Apc*$^{WT}$+OX; **h**, and *Apc*$^{Pirc/+}$+OX rats

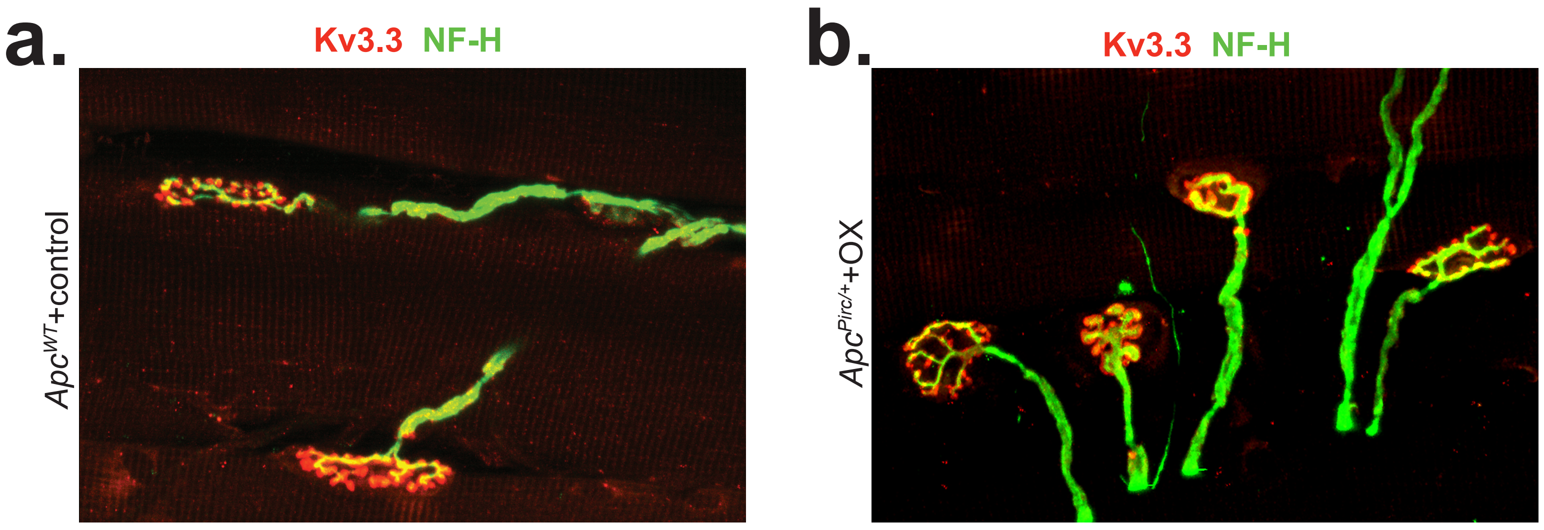

**Extended Data Figure 11. Immunostaining of neuromuscular junction.** Whole-mount staining and confocal axial projection of the neuromuscular junction showing merged images of Kv3.3 (red) and NF-H (green) from: **a,** $Apc^{WT}$ +control and **b**, $Apc^{Pirc/+}$+OX rats.

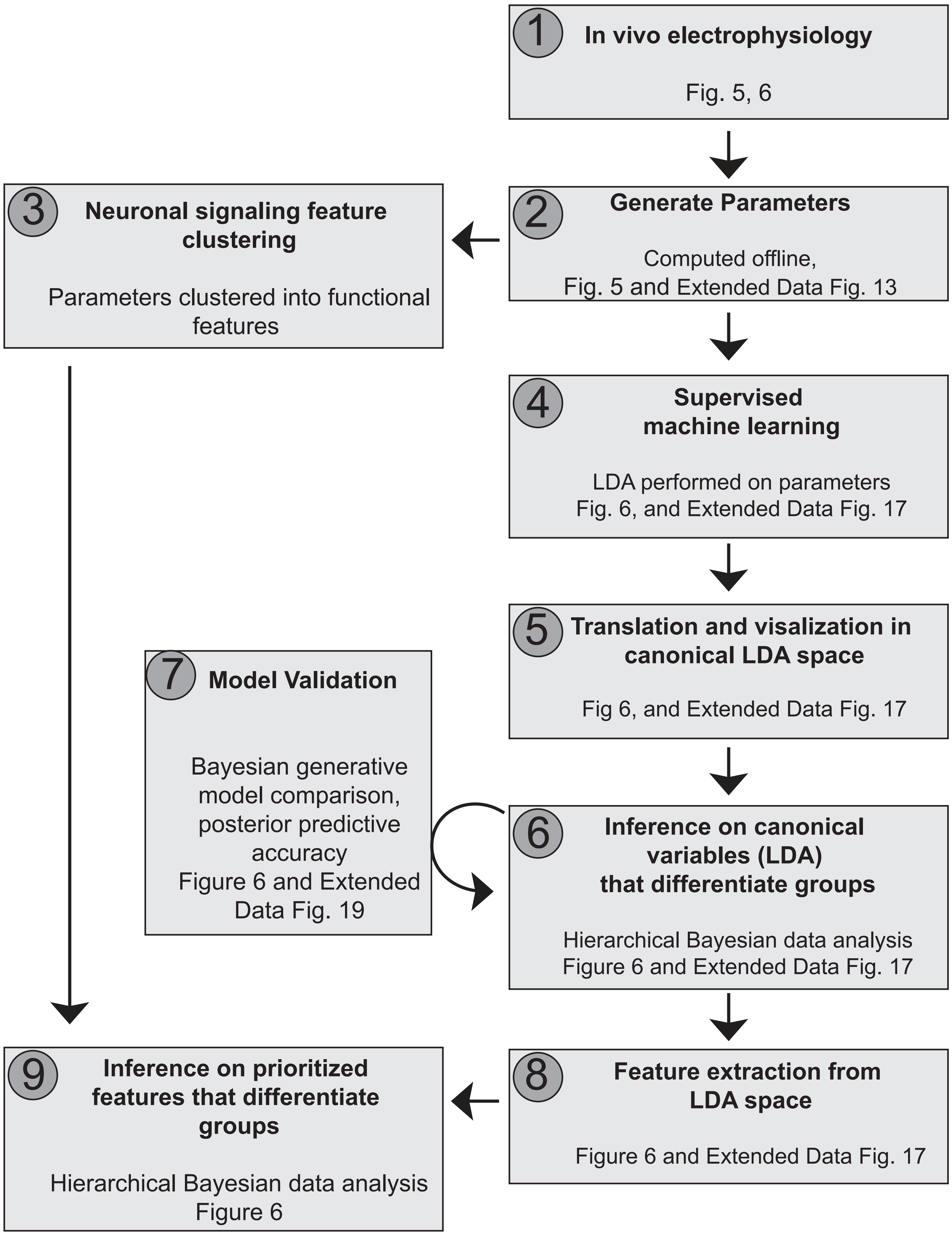

**Extended Data Figure 12. Steps of In Vivo Electrophysiology Analysis**, related to Figures 5, 6 and Extended Data Figures 13-19. Data analysis flowchart depicts the multi-step approach applied for single neuron physiological analyses, inferences and downstream model validation. A detailed description of each step is available in the **Online Methods Sections.** Linear discriminant analysis, LDA.

**a.**

| Sensitivity/Threshold Functional Cluster | | |
|---|---|---|
| 1 | Thr.L3 | Length at which 1st action potential is detected on the 3rd slow ramp (mm) |
| 2 | Thr.L | Length at which 1st action potential is detected on the fast ramp (mm) |
| 3 | Thr.T1 | Time at which 1st action potential is detected on the 1st slow ramp (ms) |
| 4 | Thr.F3 | Muscle tendon force at which 1st action potential is detected on the 3rd slow ramp (dN) |
| 5 | Thr.F1 | Muscle tendon force at which 1st action potential is detected on the 1st slow ramp (dN) |
| 6 | Thr.T | Time at which 1st action potential is detected on the fast ramp (ms) |
| 7 | Thr.F | Muscle tendon force at which 1st action potential is detected on the fast ramp (dN) |
| 8 | Thr.L1 | Length at which 1st action potential is detected on the 1st slow ramp (mm) |
| 9 | Thr.T3 | Time at which 1st action potential is detected on the 3rd slow ramp (ms) |

| Dyamic Signaling Functional Cluster | | |
|---|---|---|
| 10 | Dyn.slp1 | Slope of instantenous firing rate during the 1st slow ramp |
| 11 | Dyn.pfr1 | Peak firing rate achieved during the 1st slow ramp (Hz, pps) |
| 12 | Dyn.spkNum | Number of action potentials (spikes) during the fast ramp (#) |
| 13 | Dyn.DI | Difference between the peak dynamic firing rate and the static firing rate during half hold (Hz) |
| 14 | Dyn.spkNum3 | Number of action potentials (spikes) during the 3rd slow ramp (#) |
| 15 | Dyn.F | |
| 16 | Dyn.slp3 | Slope of instantenous firing rate during the 3rd slow ramp |
| 17 | Dyn.spkNum1 | Number of action potentials (spikes) during the 1st slow ramp (#) |
| 18 | Dyn.IB | Peak instantaneous high-frequency firing rate achieved immediately after stimulus (Hz) |
| 19 | Dyn.slp | Slope of instantenous firing rate during the fast ramp |
| 20 | Dyn.IFRdrop | Drop in instantaneous firing rate after the release on the slow ramps (Hz, pps) |
| 21 | Dyn.pfr3 | Peak firing rate achieved during the 3rd slow ramp (Hz, pps) |
| 22 | Dyn.pfr | Peak firing rate achieved during the fast ramp (Hz, pps) |

| Static Signaling Functional Cluster | | |
|---|---|---|
| 23 | Stat.mSfr | Instantenous firing rate midway through the static hold (Hz, pps) |
| 24 | Stat.afr | Average firing rate during the static hold (Hz, pps) |
| 25 | Stat.HzSTD | The standard deviation of instantaneous firing rate during the static hold |
| 26 | Stat.LSpkF | The muscle force at which the last action potential occurs during the static hold (dN). |
| 27 | Stat.slp | Slope of instantenous firing rate during the static hold. |
| 28 | Stat.LspkT | The time at which the last action potential occurs during the static hold (ms). |
| 29 | Stat.HzEnd | Instantenous firing rate during at the end of the static hold (Hz) |
| 30 | Stat.spkNum | Number of action potentials (spikes) during the static hold (#) |

| History Dependence Functional Cluster | | |
|---|---|---|
| 31 | HxDep.RDR | Difference between the number of action potentials (spikes) in the 1st and 3rd slow ramp |

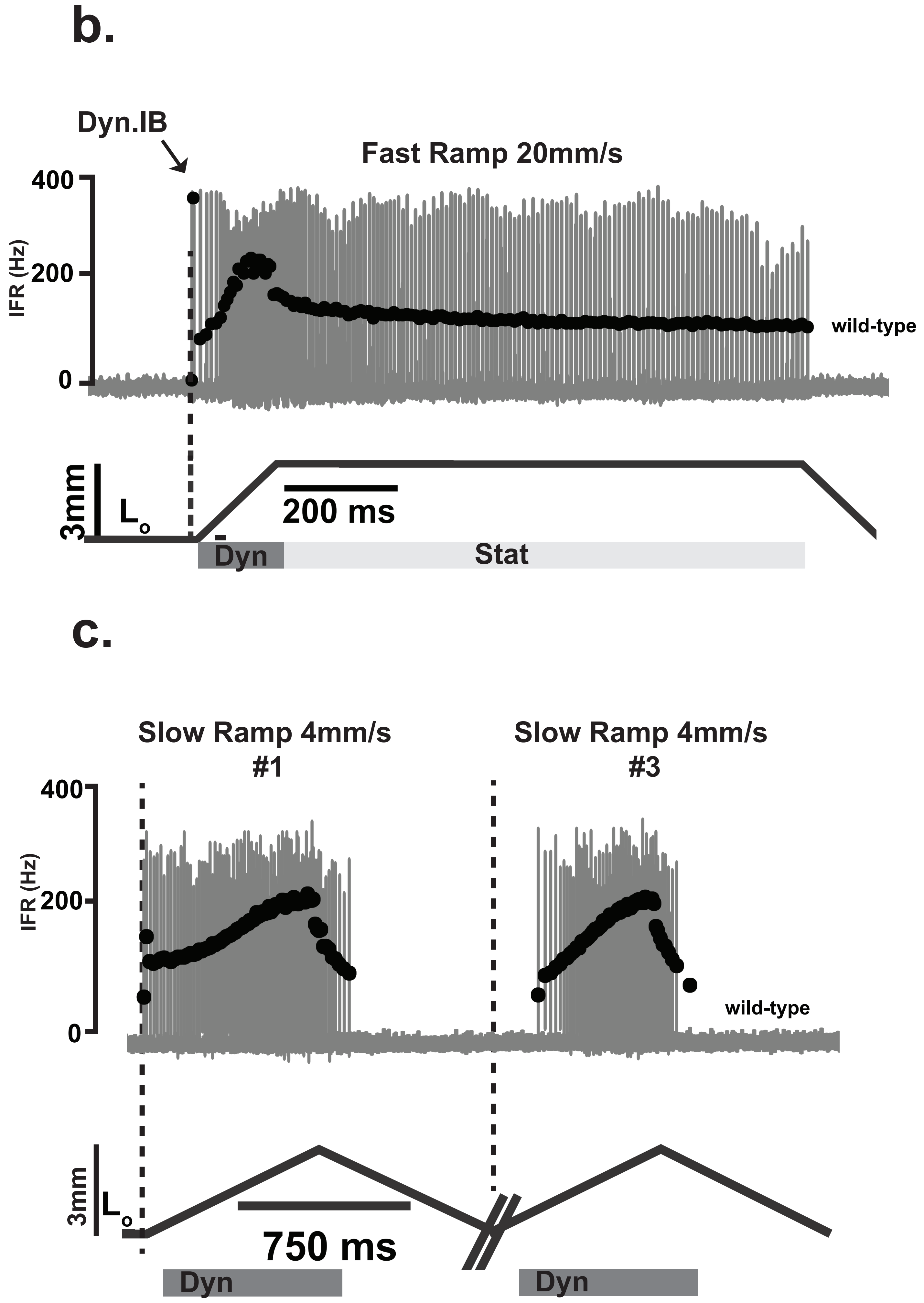

**Extended Data Figure 13. List of Parameters used for Neurophysiological Analyses,** related to Figures 2, 3 and Extended Data Figure 12. **a,** Measured and derived parameters were computed offline using custom written MATLAB scripts. Letters in the left column indicate the specific parameters included in analyses for each functional feature cluster. Thr: threshold (9 parameters); Dyn: dynamic (11 parameters); Stat: static (8 parameters); HxDep: history-dependent (1 parameter). For detailed experimental recording paradigms see Online Methods. **b**, Representative fast ramp-hold-release (3 mm at 20mm/s) trial from a proprioceptive neuron recorded from dorsal roots in *in vivo* electrophysiological experiments of a wild-type rat. **c**, Representative slow repeated ramp (3 mm at 4mm/s) trial from a proprioceptive neuron recorded from dorsal roots in *in vivo* electrophysiological experiments of a wild-type rat. Corresponding action potential trains and overlaid black circles indicate individual action potentials (spikes) and instantaneous firing rates (IFRs) of the responses. Dashed line marks the point of muscle stretch from background length ($L_o$) and indicates the starting point for threshold/sensitivity measurements. Boxes indicate dynamic (dark grey, 150 ms duration after stretch command onset) and static phases for analysis (light grey, 1 s duration after the dynamic phase)

a.

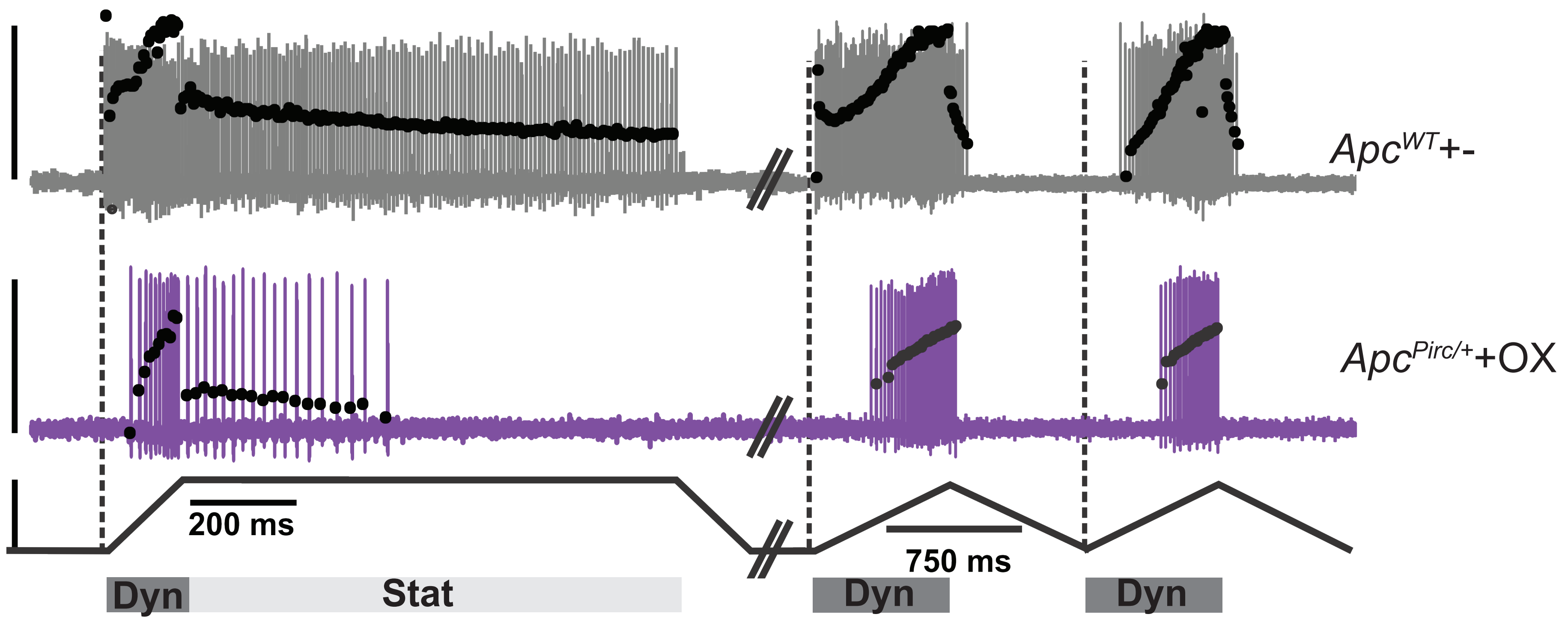

**Extended Data Figure 14. Neuronal signaling responses to slow and fast naturalistic stimulus in *Apc*$^{WT}$+control and *Apc*$^{Pirc/+}$+OX neurons,** related to Figures 5, 6. **a,** Representative fast ramp-hold-release (left, 3 mm at 20mm/s) and slow repeated ramp (right, 3 mm at 4mm/s) trial from a proprioceptive neuron recorded from dorsal roots in *in vivo* electrophysiological experiments of a *Apc*$^{WT}$+control (grey) and *Apc*$^{Pirc/+}$+OX rat (purple). Corresponding action potential trains and overlaid black circles indicate individual action potentials (spikes) and IFRs (instantaneous firing rates) of the responses. Dashed line marks the point of muscle stretch from background length ($L_o$) and indicates the starting point for threshold/sensitivity measurements. Boxes indicate dynamic (dark grey, 150 ms duration after stretch command onset) and static phases for analysis (light grey, 1 s duration after the dynamic phase).

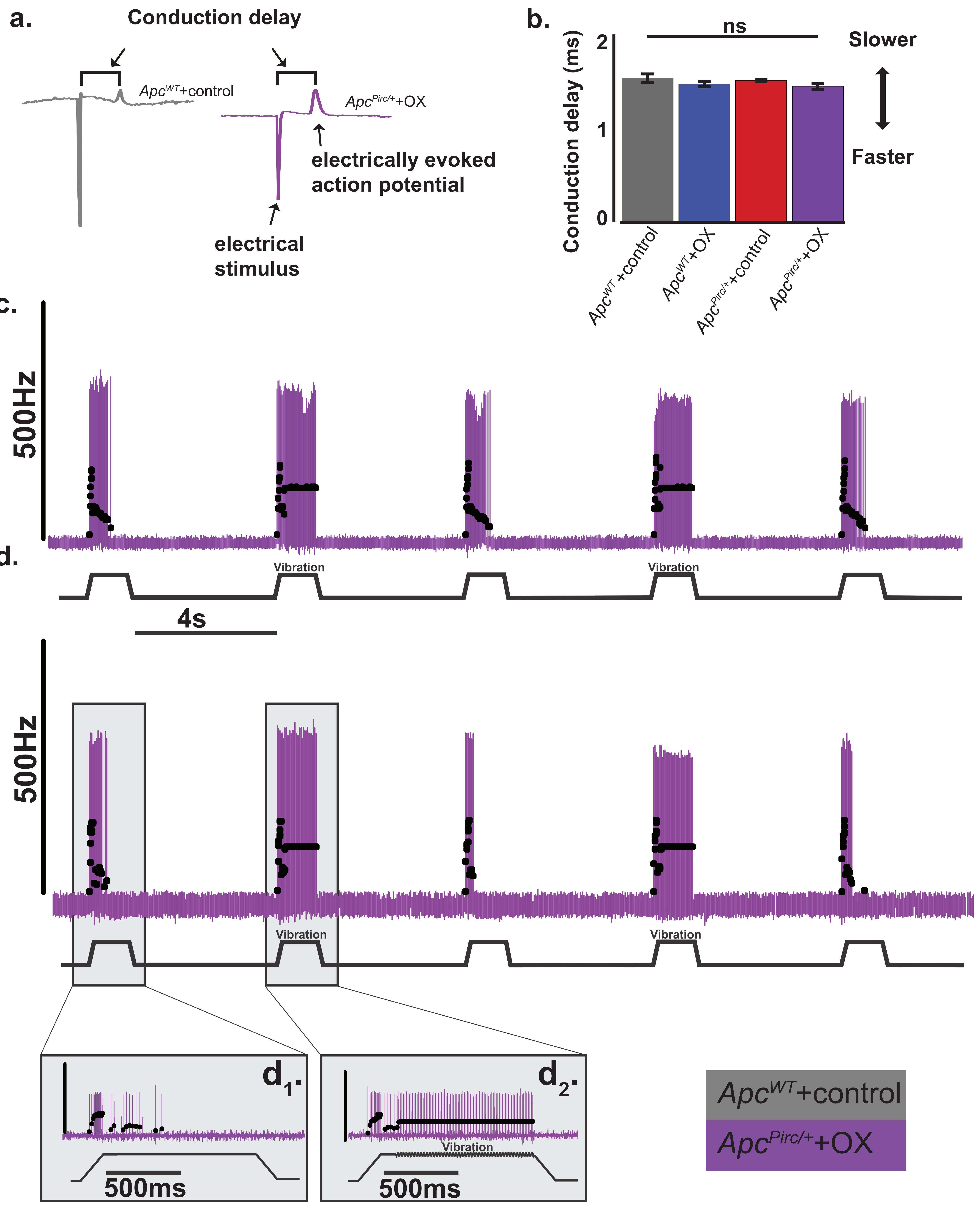

**Extended Data Figure 15. Cancer and chemotherapy interaction does not lead to electrophysiological evidence of degeneration** related to Figures 5. **a,** raw intracellular record from control and APC$^{Pirc/+}$+OX neurons shows derivation of conduction delay (ms). Conduction delay is calculated by subtracting the time from electrical stimulus to electrically evoked action potential. **b,** Mean conduction delay (ms) computed for all neurons as a mesure of axonal demyelination were analyzed from *Apc*$^{WT}$+control (n = 11), *Apc*$^{WT}$+OX (n = 19), *Apc*$^{Pirc/+}$+control(n = 20) and *Apc*$^{Pirc/+}$+OX (n = 10) neurons with hierarchical Bayesian ANOVA model. ns, no significant difference. **c & d,** Two representative stretch evoked responses of two sensory neurons (*Apc*$^{Pirc/+}$+OX ) recorded from dorsal roots in *in vivo* electrophysiological experiments with corresponding action potential trains. Overlayed black circles are IFRs of the responses. Insets show expanded view from **(b)** in a trial without (**b**$_1$) and with (**b**$_2$) superimposed vibration stimulus. **b**$_2$.

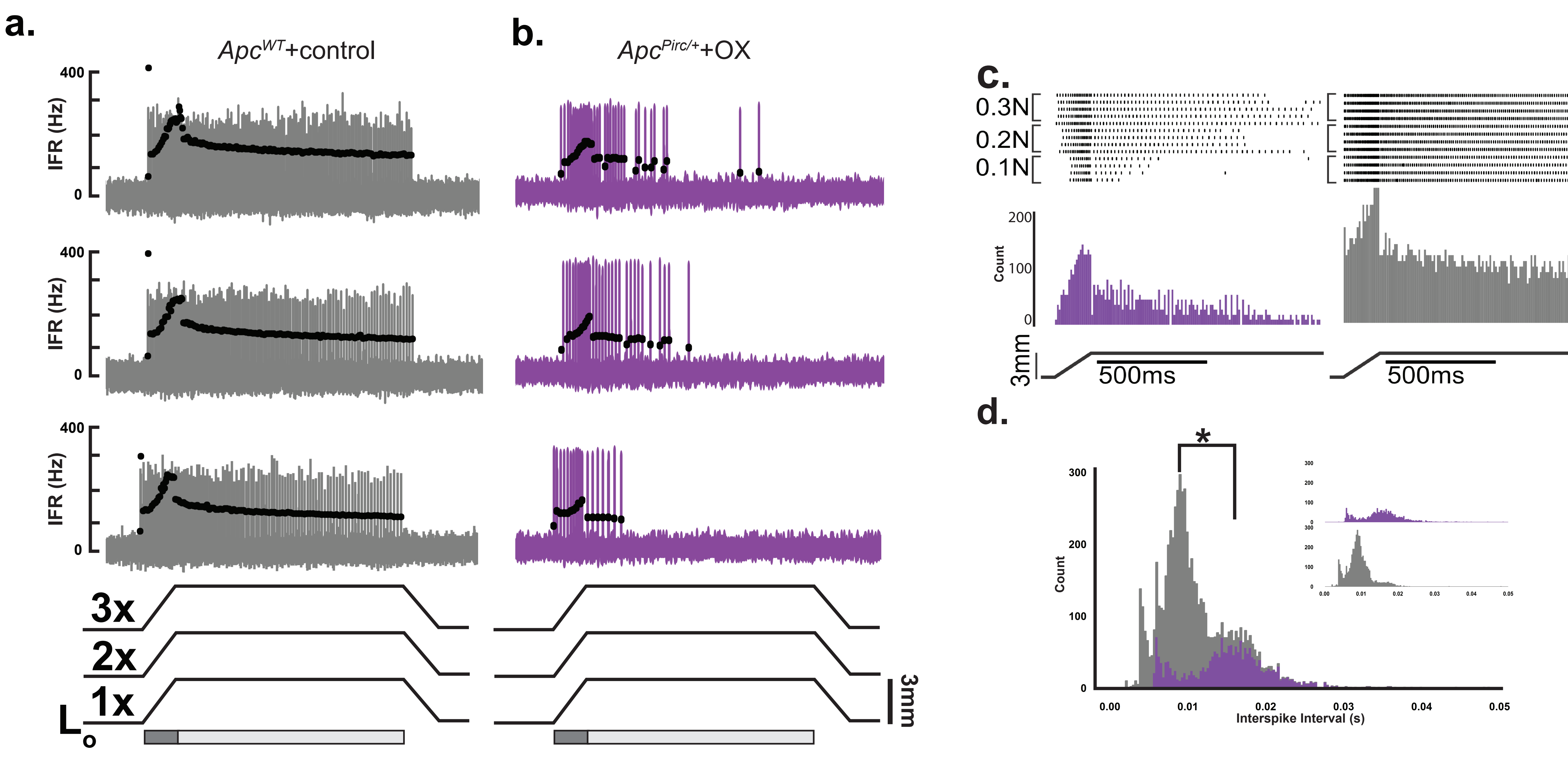

**Extended Data Figure 16. Effects of physiologic compensation in *Apc*$^{WT}$+control and *Apc*$^{Pirc/+}$+OX neurons. a, b,** Representative fast ramp-hold-release (left, 3 mm at 20mm/s) trial from a proprioceptive neuron recorded from dorsal roots in *in vivo* electrophysiological experiments of a *Apc*$^{WT}$+control (grey **a**) and *Apc*$^{Pirc/+}$+OX rat (purple **b**). Corresponding action potential trains and overlaid black circles indicate individual action potentials (spikes) and IFRs (instantaneous firing rates) of the responses. Dashed line marks the point of muscle stretch from background length ($L_o$) and indicates the starting point for threshold/sensitivity measurements. Boxes indicate dynamic (dark grey, 150 ms duration after stretch command onset) and static phases for analysis (light grey, 1 s duration after the dynamic phase). Stretch evoked (3mm) responses recorded from replicate trials at 1, 2, and 3x $L_o$ force for *Apc*$^{WT}$+control (**a**) and *Apc*$^{Pirc/+}$+OX **(b). c,** Raster plots of stretch evoked (3mm) responses from representative neurons in control (right) and APC$^{Pirc/+}$+OX (left) neurons recorded from repeated trials (4 trials) at 1, 2, and 3x background stimulus (Lo strain) intensity. Frequency histograms show cumulative distributions across 4 trials. **d,** Overlaid interval historgram (four trials per neuon at each of three stimulus intensities shows population shift. Inset show non-overlaid interval histograms for clarity. * indicates statistically significant differences between experimental groups as empirically derived from hierarchical Bayesian model (*stan_glm*): 95% highest density inter-vals do not overlap between groupwsie contrasts.

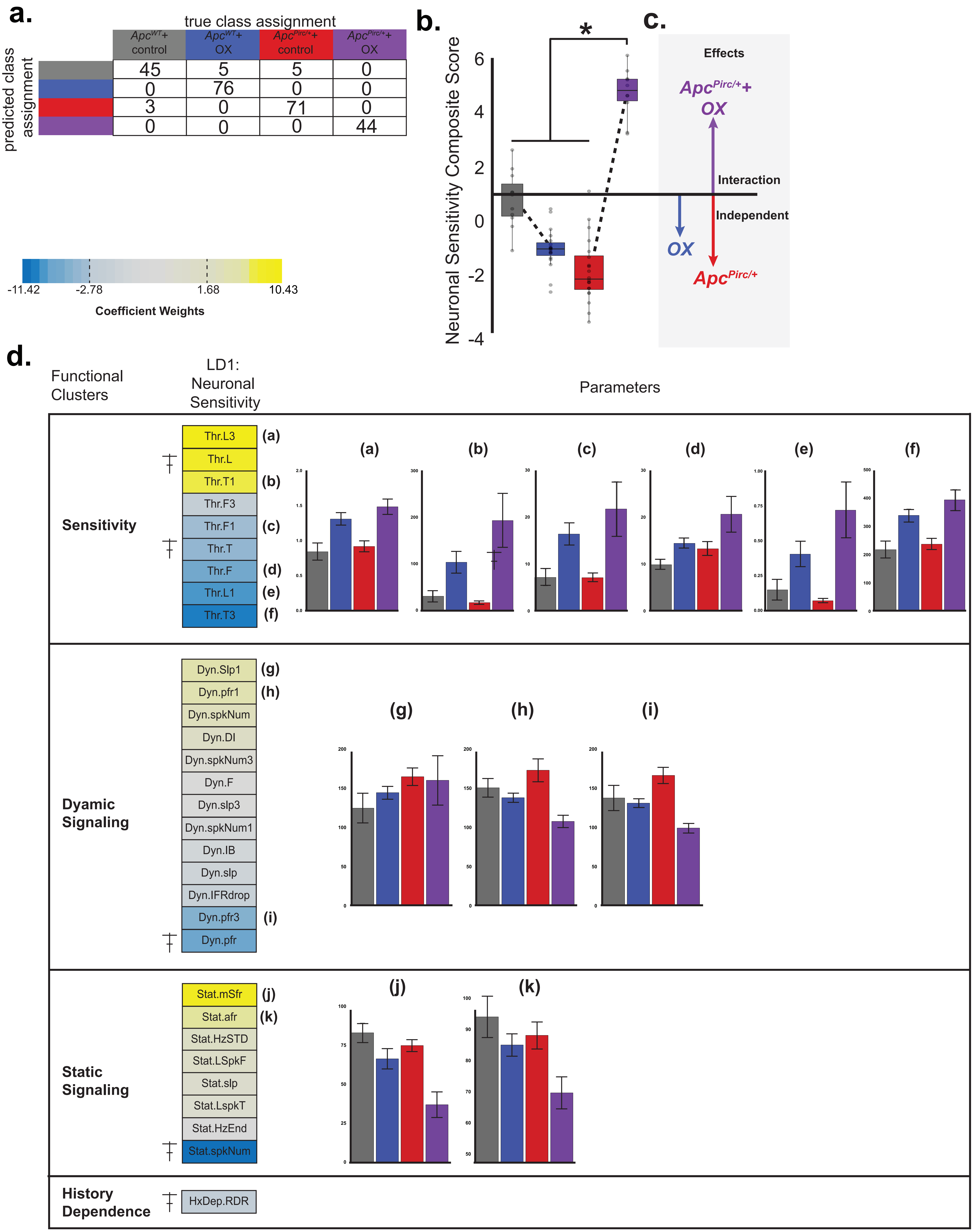

**Extended Data Figure 17. LD1 analysis of all neurons parameters,** related to Figures 6 were analyzed from *Apc*$^{WT}$+control (n = 11), *Apc*$^{WT}$+OX (n = 19), *Apc*$^{Pirc/+}$+control (n = 20) and *Apc*$^{Pirc/+}$+OX (n = 10) neurons. **a,** Confusion matrix illustrates the results of 10-fold cross validation of LDA model performance (repeated holdout method) that achieved overall 94.7% classification accuracy (posterior prediction check). Table values represent the sum of out-of-sample cross validation predictions (row) against true class assignment (column) **b,** Histogram plot, mean values of LD1 scores (Fig. 4) for each group were compared with hierarchical Bayesian ANOVA model. Asterisk indicate statistically significant differences between experimental groups in hierarchical Bayesian ANOVA, (*) indicates 95% highest density intervals (HDI) do not overlap between groupwsie contrasts. **c,** Vectors represent the magnitude and direction of the independent and combinatorial effects of cancer and/or chemotherapy. **d,** Parameters (n =31, middle) and respective discriminant coefficients weights with the interaction-specific LD1, were regrouped into functional feature clusters (left). Histogram plots report mean values for individual parameters (right, a-k) with significant contribution to LD1 loading. Symbol (┼) indicate parameters are included in Fig. 5 and 6.

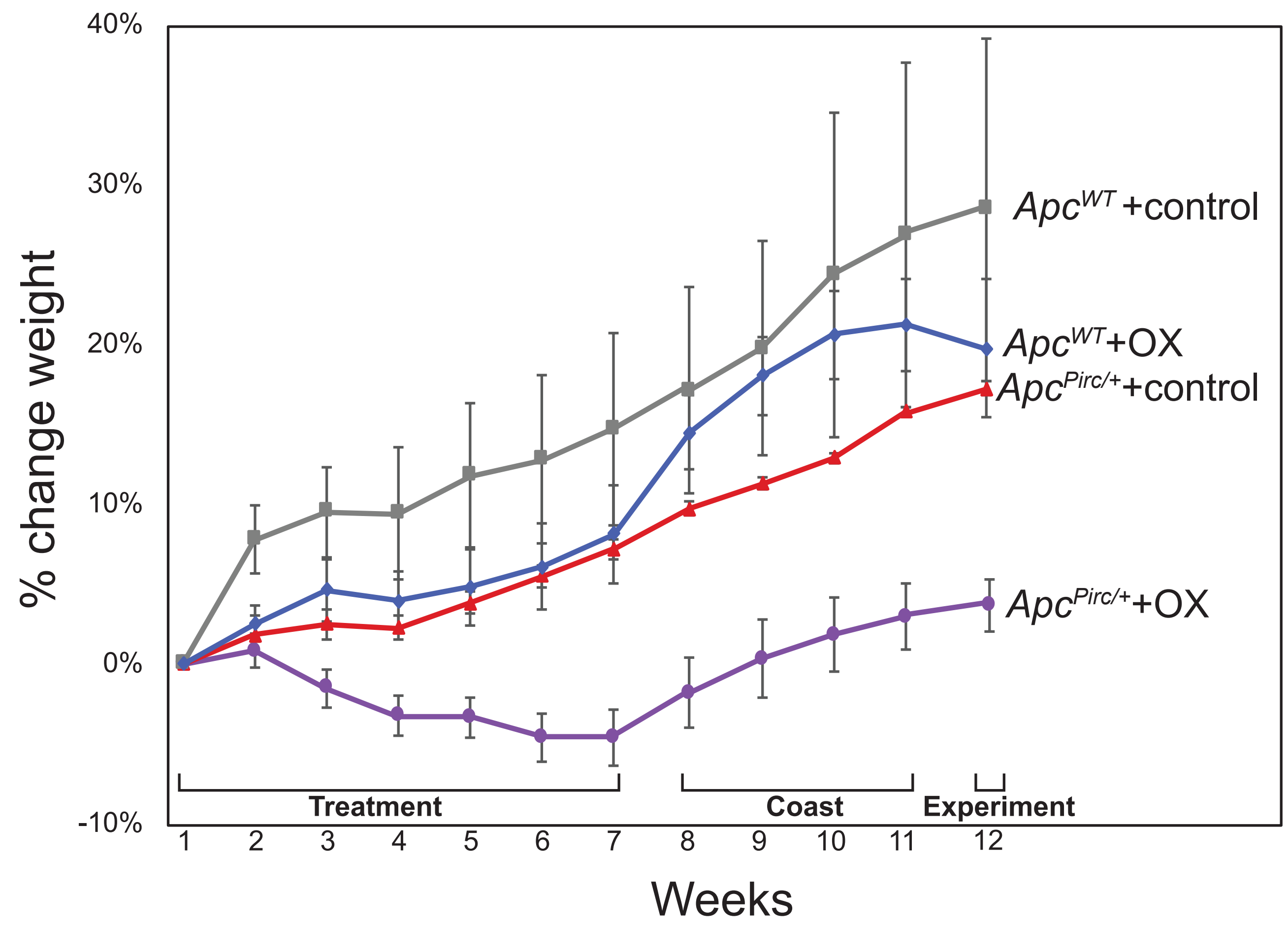

**Extended Data Figure 18. Longitudinal change in animal weight.** Longitudinal study of animal weights reveals systemic non-line ar interaction of cancer and chemotherapy. a, Weekly weight measures tracking systemic effects of cancer and/or chemotherapy. Rats included were used for transcriptional, protein, and physiologic studies. Data are mean±s.d. from 15 $Apc^{WT}$ +control, 10 $Apc^{WT}$+OX, 6 $Apc^{Pirc/+}$+control, and 8 $Apc^{Pirc/+}$+OX .

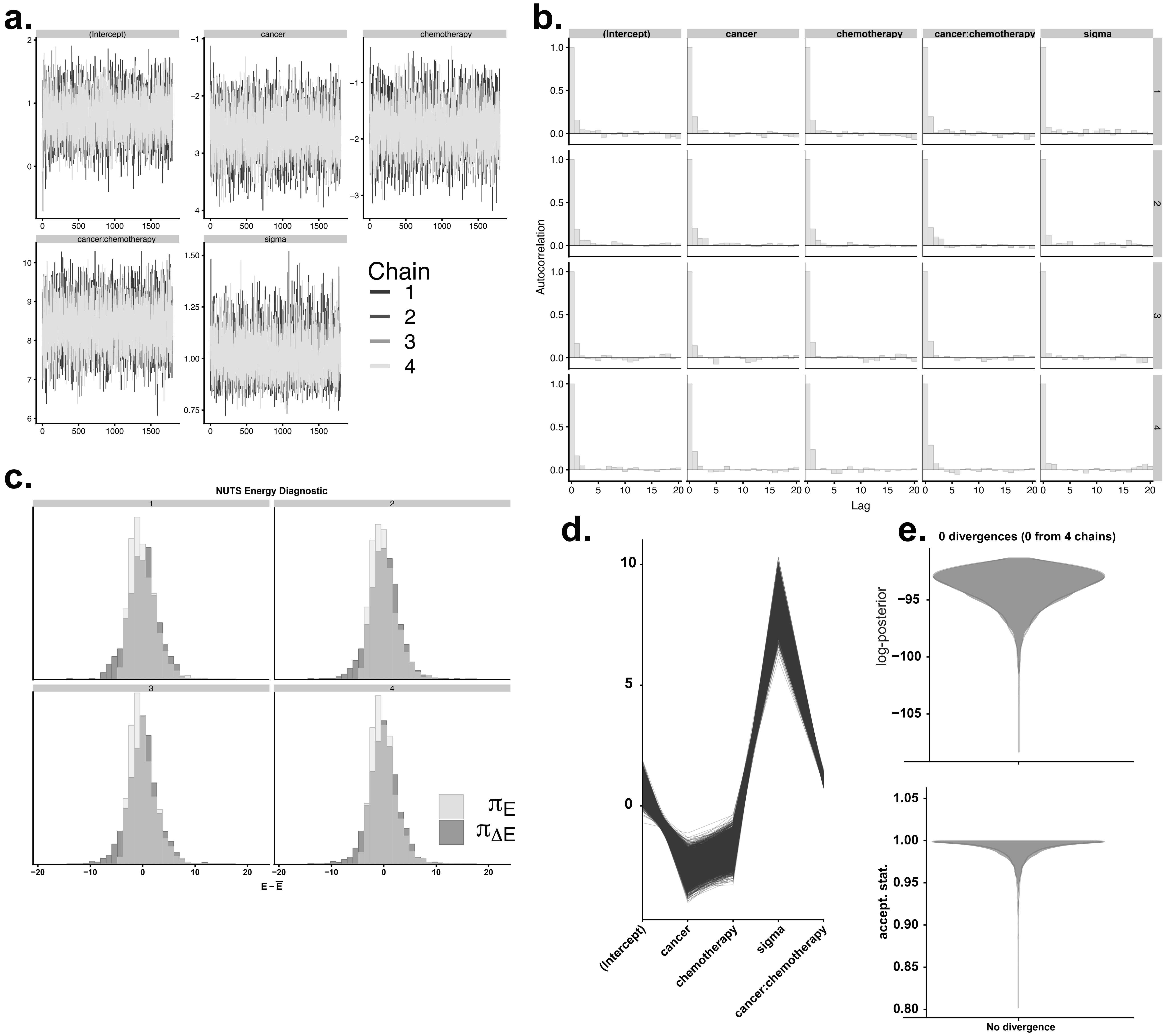

**Extended Data Figure 19. Exemplar Graphical Bayesian Model Validation. a,** Time series trace plot of the Markov chains (n=4) shows the evolution of parameter (n=5) vectors over the all iterations and indicate chains explored the full parameter space. **b,** Autocorrelation of paramters (columns) illustrate the degree of correlation between MCMC samples separated by different lags (x-axis). For example, a lag of 0 represents the degree of correlation between each MCMC sample and itself (correlation of 1). A lag of 1 represents the degree of correlation between each Markov chain Monte Carlo (MCMC) sample and the next sample along the chains (rows). Independent (uncorrelated) samples (autocorrelation of 0) indicate unbiased estimates of parameters. No parameters violate autocorrelation. **c,** Shows overlaid histograms of the (centered) marginal energy distribution $\pi_E$ and the first-differenced distribution $\pi_{\Delta E}$. The MCMC No-U-Turn Sampler (NUTS) Energy diagnostic identifies overly heavy tails that are also challenging for sampling and quantifies the heaviness of the tails of the posterior distribution. The two distributions are well-matched, meaning the random walk will explore the marginal energy distribution extremely efficiently. **d,** Plot shows one line per iteration connecting parameter values at iterations and identifies global patterns to detect divergences if present (divergences will be colored in the plot (by default in red). No divergent transitions were detected. **e,** NUTS divergence plot in the top panel shows the distribution of the log-posterior and indicates no divergences for all chains. This provides evidential support that chains adequately explore all parts of the posterior. The bottom panel shows the distribution of NUTS acceptance statistic approaches 1.

**Extended Data Table 1**

| C1a | C1a |
|---|---|
| 1 | 1 |
| 1389043_at | 1383935_at |
| 1369772_at | 1369550_at |
| 1382074_at | 1377768_at |
| 1367499_at | 1388170_at |
| 1378552_at | 1383354_a_at |
| 1389675_at | 1391582_at |
| 1374806_at | 1376628_at |
| 1367715_at | 1369391_at |
| 1392107_at | 1374911_at |
| 1369814_at | 1373730_at |
| 1370928_at | 1390391_at |
| 1374228_at | 1386611_at |
| 1397916_s_at | 1384063_at |
| 1376737_at | 1377221_at |
| 1371529_at | 1376204_at |
| 1384389_at | 1376125_at |
| 1371697_at | 1398455_at |
| 1385627_at | 1389569_at |
| 1369778_at | 1377688_at |
| 1373219_at | 1387453_at |
| 1368370_at | 1384381_at |
| 1380414_at | 1369738_s_at |
| 1390571_at | 1382404_at |
| 1371781_at | 1374422_at |
| 1390519_at | 1368947_at |
| 1388812_at | 1386181_at |
| 1371550_at | 1368856_at |
| 1387153_at | 1368527_at |
| 1369352_at | 1370141_at |
| 1367825_at | 1373035_at |
| 1390535_at | 1371079_at |
| 1387132_at | 1384507_at |
| 1378005_at | 1373062_at |
| 1371019_at | 1397848_at |
| 1391543_at | 1374113_at |
| 1375866_at | 1388672_at |
| 1369012_at | 1371988_at |
| 1378126_at | 1387995_a_at |
| 1391095_at | 1388482_at |
| 1392267_at | 1371974_at |
| 1374626_at | 1377761_at |
| 1372579_at | 1387959_at |
| 1368489_at | 1388054_a_at |

| C2b | C2b | C2b |
|---|---|---|
| 2 | 2 | 2 |
| 1395359_at | 1377821_at | 1373544_at |
| 1377617_at | 1391848_at | 1386947_at |
| 1385422_at | 1390569_at | 1392929_at |
| 1379338_at | 1389306_at | 1381449_s_at |
| 1393813_at | 1374535_at | 1382454_at |
| 1368939_a_at | 1384792_at | 1368146_at |
| 1396803_at | 1370950_at | 1387450_at |
| 1380144_at | 1390136_at | 1386811_at |
| 1393596_at | 1383240_at | 1376569_at |
| 1382312_at | 1393368_at | 1389974_at |
| 1391689_at | 1391139_at | 1376463_at |
| 1383334_at | 1368993_at | 1395744_at |
| 1390486_at | 1393576_at | 1394891_at |
| 1394972_at | 1390423_at | 1379763_at |
| 1392738_at | 1370831_at | 1377629_at |
| 1380475_at | 1392799_at | 1386064_at |
| 1377906_at | 1372480_at | 1386707_at |
| 1395100_at | 1379755_at | 1379058_at |
| 1390706_at | 1383827_at | 1381175_at |
| 1395923_at | 1385243_at | 1382413_at |
| 1396207_at | 1386979_at | 1390723_at |
| 1383786_at | 1390480_at | 1392555_at |
| 1379615_at | 1388742_at | 1389996_at |
| 1397335_at | 1370907_at | 1398445_at |
| 1381394_at | 1371430_at | 1377781_at |
| 1383849_at | 1373521_at | 1384513_at |
| 1376865_at | 1370302_at | 1376129_at |
| 1394079_at | 1387112_at | 1390351_at |
| 1385393_at | 1374216_at | 1377651_at |
| 1370932_at | 1375726_at | 1375538_at |
| 1395154_at | 1389711_at | 1379506_at |
| 1381115_at | 1376700_at | 1394834_at |
| 1394243_at | 1379444_at | 1381817_at |
| 1388000_at | 1398362_at | 1391552_at |
| 1383066_at | 1375640_at | 1380685_at |
| 1391305_at | 1395201_at | 1385889_at |
| 1395714_at | 1374448_at | 1390942_at |
| 1368858_at | 1368514_at | 1390077_at |
| 1376194_at | 1395794_at | 1394727_at |
| 1383811_at | 1373676_at | 1379511_at |
| 1385431_at | 1398387_at | 1398528_at |
| 1368825_at | 1371518_at | 1381402_at |
| 1370114_a_at | 1398354_at | 1392182_at |

| | |
|---|---|
| 1367721_at | 1371040_at |
| 1371237_a_at | 1389295_at |
| 1388271_at | 1373310_at |
| 1383860_at | 1376102_at |
| 1387558_at | 1388149_at |
| 1369885_at | 1382919_at |
| 1373225_at | 1396163_at |
| 1370023_at | 1376481_at |
| 1374468_at | 1394160_at |
| 1377210_at | 1392881_at |
| 1373984_at | 1387759_s_at |
| 1368353_at | 1370177_at |
| 1372691_at | 1395765_at |
| 1367712_at | 1392946_at |
| 1385869_at | 1371450_at |
| 1369191_at | 1370452_at |
| 1393456_at | 1385827_at |
| 1388944_at | 1388485_at |
| 1393730_at | 1383655_at |
| 1384294_at | 1368015_at |
| 1388722_at | 1393477_at |
| 1388924_at | 1383949_at |
| 1373224_at | 1368254_a_at |
| 1378800_at | 1389380_at |
| 1376151_a_at | 1376648_at |
| 1392818_at | 1369619_at |
| 1371785_at | 1368892_at |
| 1384260_at | 1376976_at |
| 1367973_at | 1385450_at |
| 1377340_at | 1373403_at |
| 1393550_at | 1394047_at |
| 1378925_at | 1393516_at |
| 1379935_at | 1380318_at |
| 1372520_at | 1369973_at |
| 1372729_at | 1370445_at |
| 1380229_at | 1370595_a_at |
| 1397437_at | 1370606_at |
| 1377103_at | 1379312_at |
| 1391643_at | 1391808_at |
| 1379311_at | 1368931_at |
| 1380772_at | 1379500_at |
| 1372211_at | 1387952_a_at |
| 1371194_at | 1393105_at |
| 1388587_at | 1383551_at |
| 1387343_at | 1370624_at |

| | | |
|---|---|---|
| 1375542_at | 1392935_at | 1380777_at |
| 1384648_at | 1372101_at | 1378745_at |
| 1384812_at | 1373151_at | 1376531_at |
| 1374780_at | 1389003_at | 1390743_at |
| 1391598_at | 1388592_at | 1396894_at |
| 1382812_at | 1391556_at | 1395521_at |
| 1382171_at | 1390411_at | 1382020_at |
| 1379412_at | 1387040_at | 1394330_at |
| 1398217_at | 1398258_at | 1380825_at |
| 1378003_at | 1389157_at | 1378548_at |
| 1370260_at | 1373122_at | 1378628_at |
| 1379578_at | 1373130_at | 1398130_at |
| 1373566_at | 1382108_at | 1385156_at |
| 1384182_at | 1392597_at | 1384692_at |
| 1376967_at | 1382732_at | 1376388_at |
| 1371003_at | 1387313_at | 1380701_at |
| 1381021_at | 1370956_at | 1393995_at |
| 1380172_at | 1370895_at | 1390302_at |
| 1394347_at | 1375266_at | 1392311_at |
| 1375523_at | 1368202_a_at | 1395436_at |
| 1369742_at | 1391448_at | 1397120_at |
| 1382064_at | 1388557_at | 1395130_at |
| 1380371_at | 1368671_at | 1396773_at |
| 1394746_at | 1380133_at | 1378486_at |
| 1376725_at | 1371847_at | 1379722_at |
| 1390813_at | 1373538_at | 1392650_at |
| 1383053_x_at | 1391012_at | 1377305_at |
| 1383455_at | 1375709_at | 1382951_at |
| 1370935_at | 1389553_at | 1377212_at |
| 1392249_at | 1384707_at | 1397608_at |
| 1391170_at | 1371394_x_at | 1378682_at |
| 1374803_at | 1385527_at | 1382911_at |
| 1381958_at | 1367971_at | 1397422_at |
| 1384133_at | 1390738_at | 1396877_at |
| 1390983_at | 1367517_at | 1396957_at |
| 1388167_at | 1393808_at | 1394837_at |
| 1370432_at | 1382691_at | 1384890_at |
| 1387146_a_at | 1372110_at | 1396450_at |
| 1368080_at | 1386943_at | 1391523_at |
| 1389214_at | 1368090_at | 1394416_at |
| 1377923_at | 1371310_s_at | 1375468_at |
| 1367860_a_at | 1373911_at | 1382847_at |
| 1373432_at | 1370155_at | 1368242_at |
| 1394574_at | 1388422_at | 1377151_at |
| 1388101_at | 1368861_a_at | 1396952_at |

| | |
|---|---|
| 1382351_at | 1373590_at |
| 1391042_at | 1381923_at |
| 1382379_at | 1369766_at |
| 1396217_at | 1390943_at |
| 1382059_at | 1387389_at |
| 1377187_at | 1368921_a_at |
| 1384147_at | 1393825_at |
| 1387606_at | 1370773_a_at |
| 1385180_at | 1369845_at |
| 1376645_at | 1370406_a_at |
| 1377387_a_at | 1385857_at |
| 1377019_at | 1395789_at |
| 1368473_at | 1398273_at |
| 1393553_at | 1370781_a_at |
| 1375041_at | 1395620_at |
| 1374137_at | 1395840_at |
| 1369035_a_at | 1368864_at |
| 1376425_at | 1382199_at |
| 1377380_at | 1386106_at |
| 1383661_at | 1381611_at |
| 1377310_at | 1371166_at |
| 1392640_at | 1383489_at |
| 1388666_at | 1370047_at |
| 1375901_at | 1386870_at |
| 1384993_at | 1386660_at |
| 1394953_at | 1393331_at |
| 1382275_at | 1382533_at |
| 1388528_at | 1368924_at |
| 1368585_at | 1368853_at |
| 1374857_at | 1387824_at |
| 1367938_at | 1369781_at |
| 1375714_at | 1393480_at |
| 1380972_at | 1383180_at |
| 1371792_at | 1385217_at |
| 1382527_at | 1368980_at |
| 1369055_at | 1384950_at |
| 1371824_at | 1378261_at |
| 1387319_at | 1368941_at |
| 1388858_at | 1395696_at |
| 1374570_at | 1369421_at |
| 1383993_at | 1369069_at |
| 1373499_at | 1388233_at |
| 1382950_at | 1370650_s_at |
| 1372750_at | 1369496_at |
| 1373421_at | 1388686_at |

| | | |
|---|---|---|
| 1372668_at | 1372643_at | 1376742_at |
| 1392166_at | 1379345_at | 1391863_at |
| 1395535_at | 1369955_at | 1390733_at |
| 1386089_at | 1370951_at | 1371776_at |
| 1370948_a_at | 1368989_at | 1390100_s_at |
| 1393702_at | 1395590_at | 1396502_at |
| 1379485_at | 1368388_at | 1380738_at |
| 1382620_at | 1372926_at | 1377589_at |
| 1384573_at | 1393933_at | 1396066_at |
| 1376299_at | 1395157_at | 1380882_at |
| 1387964_a_at | 1385191_at | 1392055_at |
| 1378775_at | 1385493_at | 1393548_at |
| 1367728_at | 1379822_at | 1378161_at |
| 1375396_at | 1392178_at | 1371771_at |
| 1369628_at | 1371186_at | 1383214_at |
| 1380062_at | 1393558_at | 1379089_at |
| 1379555_at | 1385825_at | 1390345_at |
| 1382118_at | 1370693_a_at | 1398020_at |
| 1382265_at | 1379516_at | 1382068_at |
| 1392840_at | 1370032_at | 1393594_at |
| 1397700_x_at | 1386041_a_at | 1395030_at |
| 1398716_at | 1395721_at | 1397453_at |
| 1396141_at | 1388034_at | 1375782_at |
| 1379715_at | 1387367_at | 1393674_at |
| 1369044_a_at | 1397983_at | 1395264_at |
| 1391758_at | 1371069_at | 1381121_at |
| 1368411_a_at | 1387760_a_at | 1397690_at |
| 1380730_at | 1384386_at | 1373972_at |
| 1369678_a_at | 1382561_at | 1389104_s_at |
| 1391838_at | 1396255_at | 1394467_at |
| 1397201_at | 1395251_at | 1380693_at |
| 1384355_at | 1396154_at | 1397999_at |
| 1380063_at | 1380363_at | 1378062_at |
| 1397241_at | 1384639_at | 1391304_at |
| 1395358_at | 1393606_at | 1375577_at |
| 1382565_at | 1371173_a_at | 1395365_at |
| 1394699_at | 1373358_at | 1397786_at |
| 1396676_at | 1369172_at | 1395205_at |
| 1384728_at | 1397525_at | 1376938_at |
| 1387406_at | 1397164_at | 1378814_at |
| 1397505_at | 1377445_at | 1398225_at |
| 1393866_at | 1369248_a_at | 1396775_at |
| 1397573_at | 1385364_at | 1396135_at |
| 1392269_at | 1375721_at | 1381253_at |
| 1382286_at | 1393198_at | 1368841_at |

| | |
|---|---|
| 1368420_at | 1376867_at |
| 1387868_at | 1367795_at |
| 1387037_at | 1378074_at |
| 1367940_at | 1397527_at |
| 1390969_at | 1385753_at |
| 1387202_at | 1387585_at |
| 1389754_at | 1390801_at |
| 1369737_at | 1380054_at |
| 1379910_at | 1369222_at |
| 1398482_at | 1390262_a_at |
| 1371644_at | 1380611_at |
| 1370257_at | 1369577_at |
| 1393728_at | 1385808_at |
| 1376100_at | 1368758_a_at |
| 1367741_at | 1368962_at |
| 1383486_at | 1370757_at |
| 1370080_at | 1369428_a_at |
| 1374474_at | |
| 1378420_at | |
| 1382154_at | |
| 1368266_at | |
| 1387090_a_at | |
| 1372599_at | |
| 1372752_at | |
| 1374758_at | |
| 1392502_at | |
| 1389742_at | |
| 1385568_at | |
| 1387769_a_at | |
| 1371713_at | |
| 1368645_at | |
| 1386888_at | |
| 1378196_at | |
| 1382315_at | |
| 1369871_at | |
| 1370051_at | |
| 1368487_at | |
| 1373623_at | |
| 1376775_at | |
| 1376144_at | |
| 1380110_at | |
| 1368224_at | |
| 1372236_at | |
| 1377049_at | |
| 1381775_at | |

| | | |
|---|---|---|
| 1380195_at | 1379397_at | 1395460_at |
| 1389894_at | 1395799_at | 1392026_at |
| 1389986_at | 1369323_at | 1376667_at |
| 1395092_at | 1382478_at | 1376917_at |
| 1381967_at | 1378666_at | 1394778_at |
| 1369159_at | 1393910_at | 1387350_at |
| 1390277_at | 1387563_at | 1389908_at |
| 1392730_at | 1387998_at | 1380541_at |
| 1390340_a_at | 1368589_at | 1376138_at |
| 1375739_at | 1382524_at | 1376096_a_at |
| 1391285_at | 1374283_at | 1376750_at |
| 1380696_at | 1378544_at | 1382229_at |
| 1397666_at | 1388157_at | 1379779_at |
| 1369810_at | 1375444_at | 1381489_at |
| 1390655_at | 1373759_at | 1385453_at |
| 1398125_at | 1387260_at | 1381985_at |
| 1382920_at | 1397884_at | 1372911_at |
| 1394392_at | 1386935_at | 1375767_at |
| 1381644_at | 1368453_at | 1391209_at |
| 1391906_at | 1379803_at | 1379936_at |
| 1382658_at | 1371104_at | 1384969_at |
| 1393418_at | 1387793_at | 1380168_at |
| 1370955_at | 1375214_at | 1368320_at |
| 1397851_at | 1375531_at | 1383982_at |
| 1392566_at | 1393433_at | 1394714_at |
| 1376770_at | 1393860_at | 1391701_at |
| 1376733_at | 1385641_at | 1397673_at |
| 1394985_at | 1398591_at | 1375449_at |
| 1396612_at | 1398256_at | 1384380_at |
| 1385087_at | 1368290_at | 1382749_at |
| 1376208_at | 1375043_at | 1391160_at |
| 1369093_at | 1372389_at | 1388604_at |
| 1395105_at | 1386995_at | 1377045_at |
| 1371361_at | 1368321_at | 1393662_at |
| 1368087_a_at | 1373401_at | 1384146_at |
| 1370946_at | 1387060_at | 1392704_at |
| 1377753_at | 1374974_at | 1393692_at |
| 1376313_at | 1385202_at | 1380097_at |
| 1383013_at | 1374546_at | 1368887_at |
| 1389441_at | 1389562_at | 1379753_at |
| 1390728_at | 1374171_at | 1398131_at |
| 1367968_at | 1373658_at | 1380154_at |
| 1375857_at | 1375030_at | 1381668_at |
| 1385779_at | 1385367_at | 1379815_at |
| 1383222_at | 1372479_at | 1370991_at |

| |
|---|
| 1376891_at |
| 1373975_at |
| 1387050_s_at |
| 1373286_at |
| 1391026_at |
| 1386552_at |
| 1398706_at |
| 1383574_at |
| 1373989_at |
| 1397304_at |
| 1384968_at |
| 1368332_at |
| 1368014_at |
| 1368361_a_at |
| 1374473_at |
| 1378440_at |
| 1393563_at |
| 1393316_at |
| 1372414_at |
| 1373554_at |
| 1387610_at |

| | | |
|---|---|---|
| 1374241_at | 1387094_at | 1391975_at |
| 1374708_at | 1390159_at | 1386102_at |
| 1374144_at | 1390300_at | 1378099_at |
| 1376170_at | 1379568_at | 1395249_at |
| 1385887_at | 1392157_at | 1393879_at |
| 1387854_at | 1368542_at | 1381125_at |
| 1367563_at | 1382914_at | 1391841_at |
| 1389546_at | 1389763_at | 1397692_at |
| 1373648_at | 1391167_at | 1381646_at |
| 1390404_at | 1372069_at | 1395772_at |
| 1372758_at | 1392785_at | 1397153_at |
| 1375138_at | 1387454_at | 1396254_at |
| 1387024_at | 1384841_at | 1374786_at |
| 1393663_at | 1392587_at | 1378367_at |
| 1387767_a_at | 1386026_at | 1377919_at |
| 1372440_at | 1382134_at | 1381472_at |
| 1390429_at | 1382535_at | 1396791_at |
| 1388392_at | 1368728_at | 1385164_at |
| 1388618_at | 1382599_at | 1381133_at |
| 1387897_at | 1384617_at | 1383479_at |
| 1375862_at | 1389235_at | 1394848_at |

| C2b |
| --- |
| 2 |
| 1376524_at |
| 1393029_at |
| 1385091_at |
| 1397004_at |
| 1379594_at |
| 1375689_at |
| 1392846_at |
| 1396252_at |
| 1396850_at |
| 1385177_at |
| 1378604_at |
| 1373534_at |
| 1378038_at |
| 1395728_at |
| 1394283_at |
| 1397217_at |
| 1381210_at |
| 1379150_at |
| 1377377_at |
| 1380087_at |
| 1374263_at |
| 1392368_at |
| 1395335_at |
| 1390320_at |
| 1396965_at |
| 1397624_at |
| 1382982_at |
| 1392017_at |
| 1390671_at |
| 1391579_at |
| 1375217_at |
| 1377208_at |
| 1391196_at |
| 1394729_at |
| 1380940_at |

| C2a | C2a | C2a | C2a |
| --- | --- | --- | --- |
| 3 | 3 | 3 | 3 |
| 1396348_at | 1383439_at | 1377840_at | 1385047_x_at |
| 1385668_at | 1371530_at | 1371352_at | 1393109_at |
| 1389172_at | 1394678_at | 1380175_at | 1376047_at |
| 1379971_at | 1383920_at | 1376063_at | 1373829_at |
| 1397317_at | 1384081_at | 1368295_at | 1388784_at |
| 1384996_at | 1395126_at | 1379677_at | 1382692_at |
| 1381732_at | 1374403_at | 1380405_at | 1376457_at |
| 1374043_at | 1383058_at | 1387164_at | 1388730_at |
| 1380941_at | 1380387_at | 1375660_at | 1370449_at |
| 1386059_at | 1388569_at | 1374257_at | 1370301_at |
| 1367902_at | 1376344_at | 1373642_at | 1391812_at |
| 1389734_x_at | 1389617_at | 1370334_at | 1387022_at |
| 1389413_at | 1384310_at | 1377943_at | 1370864_at |
| 1391637_at | 1379200_at | 1388511_at | 1387922_at |
| 1382173_at | 1394033_at | 1389475_at | 1386912_at |
| 1373740_at | 1393691_at | 1370959_at | 1389403_at |
| 1398732_at | 1390881_at | 1371924_at | 1391341_at |
| 1397610_at | 1371015_at | 1386899_at | 1378720_at |
| 1373748_at | 1388298_at | 1392672_at | 1390348_at |
| 1385275_at | 1388211_s_at | 1394451_at | 1385978_at |
| 1379766_at | 1368930_at | 1373240_at | 1392265_s_at |
| 1383020_at | 1391383_at | 1372070_at | 1374247_at |
| 1389034_at | 1368381_at | 1387886_at | 1382184_at |
| 1379253_at | 1396316_at | 1395003_at | 1384298_at |
| 1395439_at | 1370024_at | 1368555_at | 1393038_at |
| 1380050_at | 1379374_at | 1380088_at | 1369725_at |
| 1374572_at | 1390964_at | 1380616_at | 1373785_at |
| 1377169_at | 1374475_at | 1384127_at | 1392813_at |
| 1386881_at | 1396001_at | 1373881_at | 1383658_at |
| 1392234_at | 1378324_at | 1375247_at | 1390849_at |
| 1380267_at | 1370823_at | 1387789_at | 1384195_at |
| 1395204_at | 1389622_at | 1368563_at | 1392965_a_at |
| 1388725_at | 1371052_at | 1393259_at | 1374730_at |
| 1393641_at | 1368587_at | 1389490_at | 1385682_at |
| 1392722_at | 1370414_at | 1389059_at | 1391630_at |
| 1389697_at | 1370531_a_at | 1387323_at | 1390510_at |
| 1383008_at | 1397511_at | 1368047_at | 1371913_at |
| 1393552_x_at | 1385051_at | 1371595_at | 1381145_at |
| 1381902_at | 1398250_at | 1387655_at | 1368395_at |
| 1387571_at | 1392118_at | 1382278_at | |
| 1382296_at | 1385692_at | 1381498_at | |
| 1376036_at | 1388385_at | 1392741_at | |
| 1385252_at | 1367887_at | 1396152_s_at | |

| | | |
|---|---|---|
| 1378094_at | 1382525_at | 1388201_at |
| 1379688_at | 1372672_at | 1381179_at |
| 1384392_at | 1372213_at | 1370166_at |
| 1397808_at | 1383743_at | 1370131_at |
| 1390148_a_at | 1377392_at | 1393281_at |
| 1370391_at | 1399019_at | 1389905_at |
| 1384869_at | 1384505_at | 1389423_at |
| 1382027_at | 1381574_at | 1369181_at |
| 1384724_at | 1373903_at | 1387470_at |
| 1373661_a_at | 1370235_at | 1393356_at |
| 1389244_x_at | 1374908_at | 1385248_a_at |
| 1390406_at | 1389221_at | 1383263_at |
| 1376285_at | 1386718_at | 1393210_at |
| 1384515_at | 1398255_at | 1367631_at |
| 1389720_at | 1377375_at | 1372615_at |
| 1385432_at | 1393028_at | 1392510_at |
| 1392521_at | 1368674_at | 1372647_at |
| 1389918_at | 1377455_at | 1373175_at |
| 1374615_at | 1388876_at | 1368380_at |
| 1380312_at | 1392215_at | 1369633_at |
| 1377792_at | 1368554_at | 1372818_at |
| 1383453_at | 1383546_at | 1374273_at |
| 1393806_at | 1382358_at | 1375638_at |
| 1379075_at | 1377878_at | 1384770_at |
| 1384420_at | 1397652_at | 1383946_at |
| 1376263_at | 1383225_at | 1388176_at |
| 1384864_at | 1390283_at | 1376708_at |
| 1382096_at | 1370157_at | 1376937_at |
| 1371643_at | 1374762_at | 1368533_at |
| 1372966_at | 1373908_at | 1388880_at |
| 1398578_at | 1370310_at | 1379379_at |
| 1374871_at | 1389161_at | 1382078_at |
| 1385608_at | 1370068_at | 1377642_at |
| 1374565_at | 1374496_at | 1375962_at |
| 1389232_at | 1390138_at | 1390800_a_at |
| 1378252_at | 1373696_at | 1386261_x_at |
| 1383218_at | 1378320_at | 1389404_at |
| 1384971_at | 1370530_a_at | 1374530_at |
| 1382252_at | 1367811_at | 1367700_at |
| 1373374_at | 1383086_at | 1370960_at |
| 1375654_at | 1376084_a_at | 1376398_at |
| 1376368_at | 1398453_at | 1382905_at |
| 1398360_at | 1384273_at | 1383852_at |
| 1388425_at | 1383721_at | 1388920_at |
| 1375905_at | 1373818_at | 1387348_at |

| | | |
|---|---|---|
| 1371150_at | 1378601_at | 1389662_at |
| 1382072_at | 1379614_at | 1386901_at |
| 1385200_at | 1397343_at | 1370816_at |
| 1393398_at | 1379669_at | 1379057_at |
| 1398661_at | 1379824_at | 1374444_at |
| 1393012_at | 1378480_at | 1375856_at |
| 1379281_at | 1374104_at | 1382050_at |
| 1373210_at | 1375990_a_at | 1396150_at |
| 1392971_at | 1370657_at | 1370019_at |
| 1391911_at | 1375983_at | 1372273_at |
| 1373017_at | 1380577_at | 1368170_at |
| 1393460_at | 1368008_at | 1372111_at |
| 1368154_at | 1372587_at | 1386911_at |
| 1370097_a_at | 1369133_a_at | 1371815_at |
| 1387093_at | 1382276_at | 1373348_at |
| 1370333_a_at | 1377914_at | 1376394_at |
| 1392569_at | 1376779_at | 1370982_at |
| 1388469_at | 1389999_at | 1395184_at |
| 1374016_at | 1392932_at | 1373065_at |
| 1373725_at | 1393491_at | 1372585_at |
| 1373288_at | 1377908_at | 1373596_at |
| 1370048_at | 1389995_at | 1377452_at |
| 1377353_a_at | 1379469_at | 1367939_at |
| 1372042_at | 1367844_at | 1373000_at |
| 1371942_at | 1376516_at | 1397548_at |
| 1368344_at | 1398351_at | 1371475_at |
| 1376688_a_at | 1376355_at | 1393330_at |
| 1368474_at | 1376028_at | 1389873_at |
| 1388426_at | 1379948_at | 1372297_at |
| 1377637_at | 1375369_at | 1377239_at |
| 1393336_at | 1372775_at | 1374544_at |
| 1382613_at | 1382976_at | 1372870_at |
| 1367922_at | 1379300_at | 1386128_at |
| 1377156_at | 1380027_at | 1368558_s_at |
| 1377946_at | 1391117_at | 1390914_at |
| 1395819_at | 1389132_at | 1367850_at |
| 1396833_at | 1390940_at | 1372305_at |
| 1392972_at | 1395353_at | 1373032_at |
| 1390436_at | 1370833_at | 1374527_at |
| 1389586_at | 1378155_at | 1393227_at |
| 1392943_at | 1375305_at | 1389649_at |
| 1394761_at | 1397194_at | 1382825_at |
| 1396009_at | 1394964_at | 1368270_at |
| 1376082_at | 1385981_at | 1383189_at |
| 1389305_at | 1378409_at | 1373523_at |

| | | |
|---|---|---|
| 1389229_at | 1370307_at | 1368482_at |
| 1374874_at | 1398251_a_at | 1370987_at |
| 1372294_at | 1391803_at | 1384837_at |
| 1376105_at | 1382510_at | 1368463_at |
| 1382630_at | 1376771_at | 1385213_at |
| 1378645_at | 1379370_at | 1392515_at |
| 1376868_at | 1383284_at | 1389696_at |
| 1383840_at | 1385006_at | 1398467_at |
| 1384940_at | 1392915_at | 1393387_at |
| 1380091_at | 1384709_at | 1394766_at |
| 1385798_at | 1395000_at | 1388164_at |
| 1375984_at | 1390459_at | 1391067_at |
| 1382084_at | 1397246_at | 1379404_at |
| 1375001_at | 1371363_at | 1398407_at |
| 1379347_at | 1380459_at | 1392655_at |
| 1370420_at | 1377064_at | 1377835_at |
| 1389705_at | 1376045_at | 1385832_s_at |
| 1377342_s_at | 1378878_at | 1373490_at |
| 1367989_at | 1382190_at | 1378193_at |
| 1393228_at | 1370845_at | 1374778_at |
| 1375622_at | 1380563_at | 1373164_at |
| 1391447_at | 1390518_at | 1393249_at |
| 1374652_at | 1369944_at | 1379760_at |
| 1381190_at | 1388939_at | 1397958_at |
| 1388745_at | 1385333_at | 1387113_at |
| 1389648_at | 1368005_at | 1370516_at |
| 1373270_at | 1368322_at | 1381311_at |
| 1384211_at | 1367562_at | 1370382_at |
| 1374746_at | 1380142_at | 1385465_at |
| 1388949_at | 1369135_at | 1389123_at |
| 1373410_at | 1380248_at | 1398304_at |
| 1376575_at | 1388116_at | 1372299_at |
| 1376674_at | 1368612_at | 1387029_at |
| 1378049_at | 1379997_at | 1370883_at |
| 1393345_at | 1386833_at | 1370383_s_at |
| 1393226_at | 1394375_x_at | 1389006_at |
| 1382458_at | 1384178_at | 1373575_at |
| 1391904_at | 1382558_at | 1371033_at |
| 1393575_at | 1394497_at | 1368006_at |
| 1395410_at | 1368703_at | 1370822_at |
| 1376785_at | 1380471_at | 1390420_at |
| 1381131_at | 1380045_at | 1387947_at |
| 1381591_at | 1371429_at | 1370154_at |
| 1383197_at | 1385912_at | 1367679_at |
| 1382882_x_at | 1374558_at | 1370882_at |

| | | |
|---|---|---|
| 1367659_s_at | 1378164_at | 1392074_at |
| 1384535_at | 1376878_at | 1382531_at |
| 1377632_at | 1375658_at | 1379295_at |
| 1372125_at | 1380228_at | 1390798_at |
| 1396383_at | 1375231_a_at | 1367912_at |
| 1395472_at | 1392595_at | 1397736_at |
| 1395644_at | 1378048_at | 1385572_at |
| 1373659_at | 1387111_at | 1383708_at |
| 1368919_at | 1369751_at | 1389783_s_at |
| 1393876_at | 1385876_at | 1393688_at |
| 1389612_at | 1389324_at | 1372200_at |
| 1383414_at | 1371045_at | 1398246_s_at |
| 1378397_at | 1368339_at | 1370202_at |
| 1392928_at | 1376998_a_at | 1367791_at |
| 1380822_at | 1376240_at | 1391977_at |
| 1390037_at | 1374200_at | 1372029_at |
| 1384699_at | 1378802_at | 1368027_at |
| 1380810_at | 1370106_at | 1380894_at |
| 1381208_at | 1391618_at | 1367749_at |
| 1392170_at | 1373161_at | 1373932_at |
| 1373484_at | 1393974_at | 1379344_at |

| C2c | C2c | C2c |
|---|---|---|
| 4 | 4 | 4 |
| 1370949_at | 1387349_at | 1377111_at |
| 1384394_at | 1377857_at | 1379140_at |
| 1382307_at | 1382301_at | 1375119_at |
| 1391625_at | 1385972_at | 1369735_at |
| 1379571_at | 1396696_at | 1397632_at |
| 1393836_at | 1383343_at | 1380503_at |
| 1397362_at | 1383795_at | 1379804_at |
| 1383825_at | 1380783_at | 1397552_at |
| 1371093_at | 1397373_at | 1375453_at |
| 1390398_at | 1368255_at | 1379461_at |
| 1379101_at | 1381063_at | 1391600_at |
| 1373347_at | 1369681_at | 1375378_at |
| 1368725_at | 1395395_at | 1395711_at |
| 1381850_at | 1391549_at | 1396470_at |
| 1375459_at | 1385773_at | 1379103_at |
| 1391430_at | 1385974_at | 1368871_at |
| 1378965_at | 1390797_at | 1375212_at |
| 1393795_at | 1376836_at | 1376533_at |
| 1384376_at | 1370090_at | 1370063_at |
| 1377457_a_at | 1383064_at | 1370512_at |
| 1390710_x_at | 1391711_at | 1392441_at |
| 1383054_at | 1394401_at | 1372812_at |
| 1368401_at | 1392040_at | 1389836_a_at |
| 1369679_a_at | 1395076_at | 1371202_a_at |
| 1375174_at | 1390777_at | 1378325_at |
| 1391719_at | 1393901_at | 1398544_at |
| 1379664_at | 1370043_at | 1370052_at |
| 1393564_at | 1385215_at | 1389957_at |
| 1391416_at | 1384181_at | 1379398_at |
| 1397535_at | 1378509_at | 1391576_at |
| 1387199_a_at | 1391935_at | 1390923_a_at |
| 1383063_a_at | 1379062_at | 1382226_at |
| 1370607_a_at | 1382705_at | 1375699_at |
| 1374463_at | 1380523_at | 1375259_at |
| 1383044_at | 1378962_at | 1392666_at |
| 1382775_at | 1380325_at | 1380143_at |
| 1380069_at | 1384311_at | 1382654_at |
| 1376562_at | 1377172_at | 1377881_at |
| 1394961_at | 1392144_at | 1382953_at |
| 1372542_at | 1385017_at | 1379423_at |
| 1382130_at | 1393804_at | 1382778_at |
| 1367652_at | 1379077_at | 1385888_at |
| 1396036_at | 1375582_at | 1375612_at |

| C1b | C1b |
|---|---|
| 5 | 5 |
| 1376872_at | 1398840_at |
| 1384393_at | 1372642_at |
| 1382682_at | 1376098_a_at |
| 1375999_at | 1376215_at |
| 1393952_at | 1380457_at |
| 1372004_at | 1384564_at |
| 1393157_at | 1388544_at |
| 1385559_at | 1386321_s_at |
| 1389507_at | 1367767_at |
| 1378120_at | 1379907_at |
| 1368358_a_at | 1377739_at |
| 1385099_at | 1390042_at |
| 1393111_at | 1373120_at |
| 1373267_at | 1376754_at |
| 1392206_at | 1378586_at |
| 1372423_at | 1367834_at |
| 1379814_at | 1379401_a_at |
| 1379757_at | 1379477_at |
| 1392999_at | 1385464_at |
| 1381770_at | 1387645_at |
| 1395456_at | 1378536_at |
| 1374367_at | 1390114_at |
| 1368545_at | 1387201_at |
| 1387660_at | 1382319_at |
| 1393283_at | 1394297_at |
| 1393547_at | 1388659_at |
| 1383846_at | 1372851_at |
| 1368059_at | 1370902_at |
| 1370361_at | 1385325_at |
| 1387766_a_at | 1368692_a_at |
| 1390912_at | 1376988_at |
| 1390707_at | 1370694_at |
| 1370450_at | 1370336_at |
| 1388802_at | 1389054_at |
| 1384921_at | 1367774_at |
| 1373085_at | 1379486_at |
| 1385587_at | 1383304_at |
| 1369590_a_at | 1381341_at |
| 1369179_a_at | 1387221_at |
| 1368238_at | 1387818_at |
| 1378855_a_at | 1371259_at |
| 1374649_at | 1368187_at |
| 1396199_at | 1374330_at |

| | | |
|---|---|---|
| 1394504_at | 1372000_at | 1379535_at |
| 1383075_at | 1384542_at | 1390467_at |
| 1392523_at | 1398582_at | 1382291_at |
| 1374593_at | 1378493_at | 1390347_at |
| 1379409_at | 1382614_at | 1370221_at |
| 1390426_at | 1391474_at | 1378453_at |
| 1382952_at | 1383145_at | 1373497_at |
| 1378595_at | 1397779_at | 1393752_at |
| 1391794_at | 1380699_at | 1378740_at |
| 1392854_at | 1377974_at | 1395021_at |
| 1398846_at | 1390866_at | 1379886_at |
| 1392054_at | 1382913_at | 1385044_at |
| 1390704_at | 1384048_at | 1389442_at |
| 1379232_at | 1392717_at | 1386063_at |
| 1394784_at | 1370570_at | 1368933_at |
| 1377816_at | 1369215_a_at | 1382551_at |
| 1377105_at | 1378282_at | 1376654_at |
| 1379130_at | 1392051_at | 1391770_at |
| 1380314_at | 1375785_at | 1394756_at |
| 1376739_at | 1387204_at | 1381471_at |
| 1395616_at | 1384610_at | 1381753_at |
| 1385041_at | 1393505_x_at | 1392675_at |
| 1381809_at | 1383219_at | 1382313_at |
| 1385931_at | 1394483_at | 1385903_at |
| 1384293_at | 1391524_at | 1382258_at |
| 1380981_at | 1375469_at | 1385925_at |
| 1386525_at | 1380763_at | 1379107_at |
| 1398483_at | 1383615_a_at | 1376226_at |
| 1389868_at | 1379221_at | 1379112_at |
| 1392902_at | 1394578_at | 1389840_at |
| 1368867_at | 1382960_at | 1391968_at |
| 1393881_at | 1373773_at | 1370810_at |
| 1382308_at | 1396225_at | 1371005_at |
| 1395836_at | 1394654_at | 1391855_at |
| 1377232_at | 1392245_at | 1375724_at |
| 1377427_at | 1376744_at | 1391687_at |
| 1395064_at | 1391594_at | 1382939_at |
| 1394935_at | 1389444_at | 1378413_at |
| 1378303_at | 1385350_at | 1375426_a_at |
| 1376996_at | 1382584_at | 1390443_at |
| 1376256_at | 1379866_at | 1375303_at |
| 1391852_at | 1382898_at | 1397286_at |
| 1384809_at | 1384164_at | 1378255_at |
| 1384240_at | 1376619_at | 1384609_a_at |
| 1390463_at | 1381296_at | 1396403_at |

| | |
|---|---|
| 1370944_at | 1393441_at |
| 1373606_at | 1368072_at |
| 1376174_at | 1367624_at |
| 1393747_at | 1391121_at |
| 1370936_at | 1374819_at |
| 1368720_at | 1378605_at |
| 1383879_at | 1370695_s_at |
| 1368311_at | 1388261_at |
| 1377453_at | 1372013_at |
| 1383853_at | 1368443_at |
| 1368400_at | 1384444_at |
| 1371536_at | 1392082_a_at |
| 1368622_at | 1368708_a_at |
| 1388663_at | 1367762_at |
| 1368085_at | 1369137_at |
| 1383692_at | 1391074_at |
| 1381276_at | 1383887_at |
| 1374146_at | 1382541_at |
| 1387032_at | 1386679_at |
| 1393124_at | 1384863_at |
| 1368846_at | 1387208_at |
| 1380346_at | 1376410_at |
| 1394070_at | 1368037_at |
| 1367754_s_at | 1387395_at |
| 1368559_at | 1376788_at |
| 1369755_at | 1383220_at |
| 1372832_at | 1393957_at |
| 1371134_at | 1393190_at |
| 1369204_at | 1392077_at |
| 1389500_at | 1369165_at |
| 1387951_at | 1376417_at |
| 1391707_at | 1390473_at |
| 1371162_at | 1374095_at |
| 1378498_at | 1387799_at |
| 1370139_a_at | 1371049_at |
| 1392531_at | 1374818_at |
| 1390828_at | 1375868_at |
| 1380460_at | 1373987_at |
| 1371243_at | 1387065_at |
| 1376829_at | 1369475_x_at |
| 1397850_at | 1387578_a_at |
| 1381626_at | 1388891_at |
| 1383742_at | 1369632_a_at |
| 1377517_at | 1368595_at |
| 1376579_at | 1377176_at |

| | | |
|---|---|---|
| 1384335_at | 1397531_at | 1375215_x_at |
| 1385102_at | 1379872_at | 1384339_s_at |
| 1385101_a_at | 1393622_at | 1367958_at |
| 1394474_at | 1398440_at | 1382382_at |
| 1376490_at | 1393582_at | 1391075_at |
| 1393735_at | 1392864_at | 1381100_at |
| 1395336_at | 1378624_at | 1385240_at |
| 1384484_at | 1376627_at | 1387233_at |
| 1389972_at | 1382489_at | 1376853_at |
| 1388015_at | 1376685_at | 1375703_at |
| 1377556_at | 1390858_at | 1376436_at |
| 1382262_at | 1391757_at | 1376644_at |
| 1381229_at | 1397630_at | 1383296_a_at |
| 1393454_at | 1383726_at | 1378581_at |
| 1384509_s_at | 1377345_at | 1378368_at |
| 1384766_a_at | 1390757_at | 1380406_at |
| 1381410_a_at | 1392806_at | 1389761_at |
| 1381821_at | 1374246_at | 1394493_at |
| 1376843_at | 1395015_at | 1395318_at |
| 1381075_at | 1375763_at | 1380525_at |
| 1395762_at | 1383253_at | 1384804_at |
| 1395136_at | 1392715_at | 1394576_at |
| 1393613_at | 1393643_at | 1383833_at |
| 1383194_a_at | 1385953_at | 1397959_at |
| 1374350_at | 1392592_at | 1394422_at |
| 1385163_at | 1392198_at | 1391405_at |
| 1375486_at | 1384115_at | 1379603_at |
| 1397889_at | 1394425_at | 1383632_at |
| 1381515_at | 1379416_at | 1376523_at |
| 1383829_at | 1376933_at | 1388108_at |
| 1375532_at | 1385227_at | 1385552_at |
| 1385923_at | 1377072_at | 1377513_at |
| 1380744_at | 1392660_at | 1388866_at |
| 1391222_at | 1395986_at | 1388710_at |
| 1393189_at | 1380937_at | 1391436_at |
| 1398595_at | 1379286_at | 1379719_at |
| 1395222_at | 1385519_at | 1377686_at |
| 1391128_at | 1389989_at | 1375343_at |
| 1370387_at | 1371679_at | 1380552_at |
| 1396437_at | 1370830_at | 1378269_at |
| 1383266_at | 1377029_at | 1370089_at |
| 1385157_at | 1397676_at | 1384478_at |
| 1380446_at | 1382268_at | 1393585_at |
| 1377774_at | 1378457_at | 1371024_at |
| 1389554_at | 1384125_at | 1392044_at |

| | |
|---|---|
| 1381064_at | 1373782_a_at |
| 1368313_a_at | 1388099_a_at |
| 1369474_a_at | 1377056_at |
| 1369019_at | 1371908_at |
| 1389092_at | 1382002_at |
| 1390607_at | 1370805_at |
| 1368927_at | 1387063_at |
| 1396176_at | 1370922_at |
| 1378539_at | 1378479_at |
| 1387179_at | 1374724_at |
| 1387269_s_at | 1387360_at |
| 1396206_at | 1379545_at |
| 1389177_at | 1380016_at |
| 1394710_at | 1389461_at |
| 1384034_at | 1382975_at |
| 1368789_at | 1376443_at |
| 1383797_a_at | 1391586_at |
| 1387369_at | 1394522_at |
| 1377168_at | 1385229_at |
| 1388077_a_at | 1367888_at |
| 1367925_at | 1370850_at |
| 1373559_at | 1368667_at |
| 1391106_at | 1393163_at |
| 1368359_a_at | 1391656_at |
| 1394908_at | 1376345_at |
| 1396387_at | 1370964_at |
| 1379363_at | |
| 1393729_at | |
| 1368916_at | |
| 1393706_at | |
| 1372590_at | |
| 1369664_at | |
| 1392948_at | |
| 1387523_at | |
| 1378400_at | |
| 1373783_at | |
| 1394855_at | |
| 1378315_at | |
| 1374948_at | |
| 1398431_at | |
| 1374787_at | |
| 1389350_at | |
| 1388187_at | |
| 1368300_at | |
| 1378470_at | |

| | | |
|---|---|---|
| 1388999_at | 1390266_at | 1394626_at |
| 1383127_at | 1398420_at | 1369654_at |
| 1396550_at | 1391439_at | 1378610_at |
| 1379912_at | 1397179_at | 1396256_at |
| 1385638_at | 1392764_at | 1383552_at |
| 1378389_at | 1383112_at | 1393661_at |
| 1373463_at | 1390506_at | 1398202_at |
| 1368770_at | 1374909_at | 1369404_a_at |
| 1383527_at | 1393981_at | 1379126_at |
| 1376755_at | 1380824_at | 1398739_at |
| 1370678_s_at | 1370085_at | 1369499_at |
| 1383199_at | 1384587_at | 1391569_at |
| 1393672_at | 1390592_at | 1391187_at |
| 1382452_at | 1389998_at | 1377508_at |
| 1392842_at | 1394849_at | 1395546_at |
| 1372814_at | 1380547_at | 1376848_at |
| 1382776_at | 1392453_at | 1384168_at |
| 1379724_at | 1391089_at | 1381167_at |
| 1384250_a_at | 1373494_at | 1394824_at |
| 1391979_at | 1389116_at | 1395361_at |
| 1395333_at | 1368824_at | 1369756_a_at |
| 1375358_at | 1379733_at | 1379492_at |
| 1375011_at | 1395075_at | 1388527_at |
| 1370072_at | 1391297_at | 1369957_at |
| 1370465_at | 1392653_at | 1376623_at |
| 1382211_at | 1370267_at | 1371541_at |
| 1374676_at | 1387165_at | 1393167_at |
| 1393811_at | 1368842_at | 1398649_at |
| 1393777_at | 1377070_at | 1385354_at |
| 1375350_at | 1368958_at | 1371472_at |
| 1398500_at | 1390184_at | 1377675_at |
| 1370145_at | 1391743_at | 1371034_at |
| 1373666_at | 1376419_at | 1395255_at |
| 1376610_a_at | 1375723_at | 1371281_at |
| 1391428_at | 1383052_a_at | 1392295_a_at |
| 1390687_at | 1390871_at | 1392607_at |
| 1370205_at | 1384857_at | 1390097_at |
| 1368221_at | 1382368_at | 1392582_at |
| 1377640_at | 1394436_at | 1378904_at |
| 1390782_at | 1385108_at | |
| 1371951_at | 1394010_at | |
| 1369156_at | 1380545_at | |
| 1389632_at | 1379645_at | |
| 1368691_at | 1375552_at | |
| 1398727_at | 1379544_at | |

| |
|---|
| 1393081_at |
| 1393307_at |
| 1375925_at |
| 1369717_at |
| 1388219_at |
| 1388968_at |
| 1378557_at |
| 1372989_at |
| 1377034_at |
| 1395655_at |
| 1385434_at |
| 1371801_at |
| 1374117_at |
| 1369752_a_at |
| 1379022_at |
| 1389007_at |
| 1368412_a_at |
| 1368355_at |
| 1374672_at |
| 1372844_at |
| 1369050_at |
| 1384667_x_at |
| 1398623_at |
| 1371883_at |
| 1372755_at |
| 1377885_at |
| 1395673_at |
| 1387543_at |
| 1368194_at |
| 1385176_at |
| 1390999_at |
| 1384683_at |
| 1368197_at |
| 1378818_at |
| 1395536_at |
| 1384526_at |
| 1377867_at |
| 1384165_at |
| 1378111_at |
| 1381080_at |
| 1381233_at |
| 1376893_at |
| 1391879_at |
| 1396687_at |
| 1392425_x_at |

| | |
|---|---|
| 1377934_at | 1370035_at |
| 1392231_at | 1367884_at |
| 1377090_at | 1375524_at |
| 1374320_at | 1391438_at |
| 1374779_at | 1390048_at |
| 1380600_at | 1370176_at |
| 1385525_at | 1379509_at |
| 1387154_at | 1374066_at |
| 1379387_at | 1370262_at |
| 1384180_at | 1392180_at |
| 1379055_x_at | 1384759_at |
| 1377051_at | 1381048_at |
| 1383932_at | 1376080_at |
| 1389601_at | 1377526_at |
| 1383644_at | 1379941_at |
| 1371483_at | 1383069_at |
| 1377308_a_at | 1375545_at |
| 1377266_at | 1379313_at |
| 1372002_at | 1384948_at |
| 1383353_at | 1375278_at |
| 1393589_at | 1382303_at |

| |
|---|
| 1381173_at |
| 1372885_at |
| 1393147_at |
| 1379425_at |
| 1378274_at |
| 1393047_at |
| 1392274_at |
| 1375051_at |
| 1374632_at |
| 1372808_at |
| 1376076_at |
| 1393057_at |
| 1384522_at |
| 1382739_at |
| 1373632_at |
| 1389573_at |
| 1375052_at |
| 1376195_at |
| 1388919_at |
| 1384488_at |
| 1379493_at |

### Extended Data Table 2

| Up Regulated in APC^Pirc/++OX in relation to wild-type | | | | | | | | | |
|---|---|---|---|---|---|---|---|---|---|
| NAME | GS<br> follow li | GS DETAILS | SIZE | ES | NES | NOM p-val | FDR q-val | FWER p-val | RANK AT MAX |
| NUCLEOLAR PART | NUCLEOLAR P/ | Details ... | 58 | -0.6006561 | -2.1327953 | 0 | 0.0525942 | 0.055 | 4810 |
| PARTURITION | PARTURITION | Details ... | 18 | -0.76409274 | -2.103174 | 0 | 0.043199193 | 0.088 | 1359 |
| NEGATIVE REGULATION OF HORMONE SECRETION | NEGATIVE REG | Details ... | 70 | -0.5564646 | -2.0809677 | 0 | 0.04139415 | 0.123 | 1315 |
| TRANSLATIONAL ELONGATION | TRANSLATION/ | Details ... | 106 | -0.51414555 | -2.071338 | 0 | 0.03434976 | 0.135 | 5412 |
| REGULATION OF TYROSINE PHOSPHORYLATION OF STAT3 PROTEIN | REGULATION C | Details ... | 38 | -0.61759853 | -2.0574071 | 0 | 0.03441299 | 0.165 | 1359 |
| ACUTE PHASE RESPONSE | ACUTE PHASE | Details ... | 34 | -0.6394826 | -2.0536501 | 0 | 0.02992805 | 0.172 | 976 |
| TRANSLATIONAL TERMINATION | TRANSLATION/ | Details ... | 90 | -0.5214301 | -2.052021 | 0 | 0.026459541 | 0.178 | 5033 |
| CELLULAR RESPONSE TO INTERLEUKIN 6 | CELLULAR RES | Details ... | 20 | -0.71682686 | -2.0428407 | 0 | 0.027400717 | 0.204 | 135 |
| CD4 POSITIVE ALPHA BETA T CELL ACTIVATION | CD4 POSITIVE / | Details ... | 29 | -0.6542219 | -2.0407088 | 0 | 0.025515366 | 0.212 | 1359 |
| NEUROPEPTIDE HORMONE ACTIVITY | NEUROPEPTID | Details ... | 25 | -0.6644572 | -2.0165114 | 0 | 0.030778589 | 0.271 | 3848 |
| RESPONSE TO INTERLEUKIN 6 | RESPONSE TO | Details ... | 24 | -0.6991114 | -2.0083828 | 0 | 0.03079357 | 0.292 | 783 |
| REGULATION OF CIRCADIAN SLEEP WAKE CYCLE | REGULATION C | Details ... | 23 | -0.6778624 | -1.9815817 | 0 | 0.041674998 | 0.401 | 1359 |
| MITOCHONDRIAL TRANSLATION | MITOCHONDR | Details ... | 103 | -0.49125695 | -1.9778397 | 0 | 0.040424347 | 0.416 | 5033 |
| CELLULAR RESPONSE TO CORTICOSTEROID STIMULUS | CELLULAR RES | Details ... | 55 | -0.5542193 | -1.9655665 | 0 | 0.04424591 | 0.477 | 459 |
| RESPONSE TO INTERLEUKIN 1 | RESPONSE TO | Details ... | 91 | -0.49679404 | -1.9585394 | 0 | 0.045131046 | 0.516 | 1171 |
| PRERIBOSOME | PRERIBOSOME | Details ... | 52 | -0.5536905 | -1.9569926 | 0 | 0.04265956 | 0.518 | 5581 |
| MONOCYTE CHEMOTAXIS | MONOCYTE CH | Details ... | 25 | -0.66094345 | -1.9483496 | 0 | 0.04529107 | 0.561 | 470 |
| POSITIVE REGULATION OF T HELPER CELL DIFFERENTIATION | POSITIVE REGU | Details ... | 15 | -0.7229009 | -1.9468211 | 0.002105263 | 0.043553904 | 0.568 | 2 |
| REGULATION OF VASODILATION | REGULATION C | Details ... | 42 | -0.56943715 | -1.9215052 | 0 | 0.057796985 | 0.694 | 2209 |
| PEPTIDE SECRETION | PEPTIDE SECR | Details ... | 55 | -0.53095704 | -1.9176296 | 0 | 0.05707113 | 0.703 | 319 |
| MATURATION OF SSU RRNA | MATURATION ( | Details ... | 39 | -0.5764421 | -1.9105362 | 0 | 0.058608096 | 0.724 | 4894 |
| RIBOSOME BIOGENESIS | RIBOSOME BIO | Details ... | 280 | -0.41458735 | -1.9035401 | 0 | 0.06043072 | 0.753 | 4909 |
| REGULATION OF T HELPER CELL DIFFERENTIATION | REGULATION C | Details ... | 19 | -0.6855839 | -1.9009429 | 0.002227172 | 0.0598399 | 0.764 | 2 |
| POSITIVE REGULATION OF TYROSINE PHOSPHORYLATION OF STAT3 PR | POSITIVE REGU | Details ... | 31 | -0.607626 | -1.899474 | 0 | 0.05859747 | 0.771 | 1359 |
| ARACHIDONIC ACID METABOLIC PROCESS | ARACHIDONIC | Details ... | 30 | -0.6091334 | -1.8987209 | 0 | 0.05666881 | 0.772 | 1316 |
| NEGATIVE REGULATION OF BEHAVIOR | NEGATIVE REG | Details ... | 16 | -0.7276433 | -1.8984941 | 0 | 0.054633893 | 0.773 | 1359 |
| RIBOSOME | RIBOSOME | Details ... | 205 | -0.43139622 | -1.8959061 | 0 | 0.05382831 | 0.783 | 6908 |
| NEUTROPHIL MEDIATED IMMUNITY | NEUTROPHIL M | Details ... | 15 | -0.7247133 | -1.8949344 | 0.002057613 | 0.05227384 | 0.785 | 275 |
| SMALL NUCLEOLAR RIBONUCLEOPROTEIN COMPLEX | SMALL NUCLE( | Details ... | 18 | -0.689783 | -1.8903326 | 0 | 0.053187467 | 0.806 | 3711 |
| REGULATION OF NEUROTRANSMITTER UPTAKE | REGULATION C | Details ... | 15 | -0.7222038 | -1.8896916 | 0 | 0.051758274 | 0.809 | 161 |
| ARGININE METABOLIC PROCESS | ARGININE MET | Details ... | 17 | -0.68985146 | -1.8859904 | 0.002192983 | 0.0527589 | 0.828 | 484 |
| NEUROPEPTIDE SIGNALING PATHWAY | NEUROPEPTID | Details ... | 76 | -0.49999642 | -1.8819045 | 0 | 0.053985182 | 0.843 | 2640 |
| HORMONE ACTIVITY | HORMONE AC | Details ... | 86 | -0.48289585 | -1.8786637 | 0 | 0.05448446 | 0.853 | 1128 |
| RESPONSE TO DEXAMETHASONE | RESPONSE TO | Details ... | 33 | -0.58670306 | -1.8772223 | 0 | 0.05401279 | 0.859 | 1038 |
| DEFENSE RESPONSE TO GRAM POSITIVE BACTERIUM | DEFENSE RES | Details ... | 45 | -0.5426134 | -1.8711301 | 0.002227172 | 0.05619107 | 0.877 | 360 |
| CELLULAR RESPONSE TO DEXAMETHASONE STIMULUS | CELLULAR RES | Details ... | 27 | -0.6205979 | -1.8705033 | 0 | 0.05502473 | 0.878 | 459 |
| PEPTIDE HORMONE RECEPTOR BINDING | PEPTIDE HORM | Details ... | 16 | -0.71249783 | -1.870344 | 0 | 0.05371582 | 0.878 | 1038 |
| NEGATIVE REGULATION OF CYTOKINE BIOSYNTHETIC PROCESS | NEGATIVE REG | Details ... | 23 | -0.6406753 | -1.8698264 | 0 | 0.052722864 | 0.879 | 197 |
| REGULATION OF NEUROLOGICAL SYSTEM PROCESS | REGULATION C | Details ... | 61 | -0.51299596 | -1.8697813 | 0 | 0.051370997 | 0.879 | 1373 |
| RIBOSOME ASSEMBLY | RIBOSOME AS: | Details ... | 47 | -0.54161465 | -1.8631577 | 0.002375297 | 0.053890973 | 0.898 | 4181 |
| RRNA METABOLIC PROCESS | RRNA METABC | Details ... | 232 | -0.42109925 | -1.8628715 | 0 | 0.05273684 | 0.898 | 5924 |

| | | | | | | | | | |
|---|---|---|---|---|---|---|---|---|---|
| ALPHA BETA T CELL ACTIVATION | ALPHA BETA T | Details ... | 42 | -0.551507 | -1.8624436 | 0 | 0.051684488 | 0.9 | 1359 |
| REGULATION OF RENAL SYSTEM PROCESS | REGULATION ( | Details ... | 34 | -0.5793682 | -1.8566442 | 0 | 0.053712733 | 0.909 | 2943 |
| 90S PRERIBOSOME | 90S PRERIBOS | Details ... | 22 | -0.6464757 | -1.8565722 | 0.006564552 | 0.052556306 | 0.909 | 5017 |
| POSITIVE REGULATION OF SMOOTH MUSCLE CONTRACTION | POSITIVE REGU | Details ... | 26 | -0.6096725 | -1.8508208 | 0.002020202 | 0.054939564 | 0.923 | 1023 |
| REGULATION OF NITRIC OXIDE SYNTHASE BIOSYNTHETIC PROCESS | REGULATION ( | Details ... | 16 | -0.6895947 | -1.8404835 | 0.00210084 | 0.060514305 | 0.944 | 1359 |
| POSITIVE REGULATION OF VASODILATION | POSITIVE REGU | Details ... | 29 | -0.59542114 | -1.8394903 | 0 | 0.059689015 | 0.944 | 900 |
| RESPONSE TO COLD | RESPONSE TO | Details ... | 37 | -0.5528324 | -1.8353885 | 0.002192983 | 0.061184622 | 0.951 | 1150 |
| POSITIVE REGULATION OF INFLAMMATORY RESPONSE | POSITIVE REGU | Details ... | 83 | -0.46864662 | -1.8312312 | 0 | 0.0628278 | 0.956 | 917 |
| SMALL SUBUNIT PROCESSOME | SMALL SUBUNI | Details ... | 30 | -0.58187366 | -1.8272423 | 0.002145923 | 0.06401419 | 0.963 | 5765 |
| VASCULAR PROCESS IN CIRCULATORY SYSTEM | VASCULAR PR( | Details ... | 152 | -0.43414718 | -1.8268363 | 0 | 0.06308989 | 0.964 | 2107 |
| INTRINSIC APOPTOTIC SIGNALING PATHWAY IN RESPONSE TO ENDOPL | INTRINSIC APC | Details ... | 29 | -0.57496345 | -1.8192376 | 0.002232143 | 0.06667311 | 0.969 | 2283 |
| NCRNA PROCESSING | NCRNA PROCE | Details ... | 332 | -0.3894126 | -1.8188677 | 0 | 0.065573536 | 0.971 | 5935 |
| POSITIVE REGULATION OF BLOOD CIRCULATION | POSITIVE REGU | Details ... | 87 | -0.47386122 | -1.8163941 | 0 | 0.066034235 | 0.975 | 1023 |
| POSITIVE REGULATION OF IMMUNOGLOBULIN PRODUCTION | POSITIVE REGU | Details ... | 29 | -0.5826723 | -1.8153062 | 0.008810572 | 0.065484665 | 0.975 | 508 |
| PORPHYRIN CONTAINING COMPOUND METABOLIC PROCESS | PORPHYRIN C( | Details ... | 34 | -0.56016403 | -1.81263 | 0.002392344 | 0.06625171 | 0.977 | 3591 |
| ACUTE INFLAMMATORY RESPONSE | ACUTE INFLAM | Details ... | 62 | -0.49246898 | -1.8090931 | 0 | 0.06716453 | 0.98 | 976 |
| POSITIVE REGULATION OF VASOCONSTRICTION | POSITIVE REGU | Details ... | 35 | -0.55808604 | -1.8064594 | 0.004705882 | 0.06786971 | 0.983 | 1005 |
| REGULATION OF BEHAVIOR | REGULATION ( | Details ... | 55 | -0.50086963 | -1.8056107 | 0 | 0.067197666 | 0.983 | 1569 |
| REGULATION OF EXCRETION | REGULATION ( | Details ... | 27 | -0.58789206 | -1.801138 | 0.006741573 | 0.06911023 | 0.986 | 1359 |
| LARGE RIBOSOMAL SUBUNIT | LARGE RIBOSC | Details ... | 85 | -0.46640182 | -1.799376 | 0 | 0.06899603 | 0.988 | 6761 |
| REGULATION OF RENAL SODIUM EXCRETION | REGULATION ( | Details ... | 22 | -0.6254262 | -1.7989463 | 0.010729614 | 0.06820163 | 0.988 | 2338 |
| REGULATION OF SYNAPTIC TRANSMISSION DOPAMINERGIC | REGULATION ( | Details ... | 16 | -0.67172843 | -1.7976642 | 0.006355932 | 0.068061754 | 0.989 | 138 |
| RIBOSOMAL SUBUNIT | RIBOSOMAL SU | Details ... | 149 | -0.41884616 | -1.7883054 | 0 | 0.073387496 | 0.993 | 7124 |
| RRNA TRANSCRIPTION | RRNA TRANSC | Details ... | 16 | -0.6773662 | -1.7872491 | 0.002061856 | 0.07292507 | 0.994 | 1062 |
| SYNAPTIC TRANSMISSION CHOLINERGIC | SYNAPTIC TRA | Details ... | 31 | -0.562881 | -1.7865131 | 0 | 0.07241493 | 0.995 | 4307 |
| NCRNA METABOLIC PROCESS | NCRNA METAE | Details ... | 451 | -0.37245378 | -1.7856072 | 0 | 0.07198153 | 0.995 | 5986 |
| RESPONSE TO SALT | RESPONSE TO | Details ... | 16 | -0.67460734 | -1.7760954 | 0.002087683 | 0.07748433 | 0.998 | 1359 |
| POSITIVE REGULATION OF AMINE TRANSPORT | POSITIVE REGU | Details ... | 32 | -0.55732644 | -1.7756541 | 0.004310345 | 0.0766345 | 0.999 | 386 |
| MATURATION OF SSU RRNA FROM TRICISTRONIC RRNA TRANSCRIPT | MATURATION ( | Details ... | 32 | -0.5586658 | -1.7755531 | 0.006864989 | 0.0756209 | 0.999 | 4894 |
| NEUROPEPTIDE RECEPTOR BINDING | NEUROPEPTID | Details ... | 23 | -0.6144161 | -1.7745613 | 0.002132196 | 0.075324066 | 0.999 | 1038 |
| POSITIVE REGULATION OF NEUROLOGICAL SYSTEM PROCESS | POSITIVE REGU | Details ... | 17 | -0.65589446 | -1.7711015 | 0.004273505 | 0.07657306 | 0.999 | 716 |
| ALPHA BETA T CELL DIFFERENTIATION | ALPHA BETA T | Details ... | 37 | -0.54395115 | -1.7666535 | 0.004672897 | 0.079118565 | 0.999 | 1359 |
| DEATH RECEPTOR BINDING | DEATH RECEP | Details ... | 17 | -0.6517083 | -1.7654413 | 0.00409836 | 0.07893731 | 0.999 | 4370 |
| REGULATION OF RESPONSE TO INTERFERON GAMMA | REGULATION ( | Details ... | 22 | -0.6064194 | -1.7622092 | 0.006410257 | 0.080279015 | 0.999 | 299 |
| T CELL DIFFERENTIATION INVOLVED IN IMMUNE RESPONSE | T CELL DIFFER | Details ... | 24 | -0.5961816 | -1.7616808 | 0.006535948 | 0.07964152 | 0.999 | 140 |
| CELLULAR RESPONSE TO INTERLEUKIN 1 | CELLULAR RES | Details ... | 67 | -0.47468987 | -1.7604797 | 0.002525253 | 0.07932866 | 0.999 | 1359 |
| POSITIVE REGULATION OF MUSCLE CONTRACTION | POSITIVE REGU | Details ... | 39 | -0.5274248 | -1.7587407 | 0.004926108 | 0.079649255 | 0.999 | 400 |
| FERTILIZATION | FERTILIZATION | Details ... | 90 | -0.4426672 | -1.7575744 | 0 | 0.079412885 | 0.999 | 3191 |
| MATURATION OF 5 8S RRNA | MATURATION ( | Details ... | 27 | -0.5572005 | -1.7552909 | 0.004728132 | 0.08021349 | 0.999 | 4716 |
| NEGATIVE REGULATION OF COAGULATION | NEGATIVE REG | Details ... | 41 | -0.5172662 | -1.7550461 | 0.002403846 | 0.07939759 | 0.999 | 1917 |
| ORGAN OR TISSUE SPECIFIC IMMUNE RESPONSE | ORGAN OR TIS | Details ... | 17 | -0.656976 | -1.7540039 | 0.010373444 | 0.079255655 | 0.999 | 44 |
| RESPONSE TO AMINE | RESPONSE TO | Details ... | 45 | -0.51388276 | -1.7515258 | 0.002283105 | 0.08031664 | 0.999 | 739 |
| REACTIVE NITROGEN SPECIES METABOLIC PROCESS | REACTIVE NITI | Details ... | 17 | -0.6334953 | -1.7510895 | 0.008385744 | 0.07971953 | 0.999 | 1839 |

| | | | | | | | | | |
|---|---|---|---|---|---|---|---|---|---|
| NEGATIVE REGULATION OF FAT CELL DIFFERENTIATION | NEGATIVE REG | Details ... | 36 | -0.532201 | -1.7431977 | 0.004444445 | 0.084675536 | 0.999 | 319 |
| REGULATION OF IMMUNOGLOBULIN PRODUCTION | REGULATION ( | Details ... | 44 | -0.50745904 | -1.7398577 | 0.002386635 | 0.086245276 | 1 | 1368 |
| REGULATION OF GLUCONEOGENESIS | REGULATION ( | Details ... | 32 | -0.5385388 | -1.7388082 | 0.006622517 | 0.08608628 | 1 | 52 |
| RESPONSE TO IMMOBILIZATION STRESS | RESPONSE TO | Details ... | 20 | -0.6245355 | -1.7342497 | 0.008849558 | 0.08880805 | 1 | 1401 |
| HEME METABOLIC PROCESS | HEME METABO | Details ... | 28 | -0.5546032 | -1.730616 | 0.004555809 | 0.0910437 | 1 | 3591 |
| RIBONUCLEOPROTEIN COMPLEX BIOGENESIS | RIBONUCLEOP | Details ... | 394 | -0.36640283 | -1.7248359 | 0 | 0.094531156 | 1 | 6143 |
| LEUKOCYTE APOPTOTIC PROCESS | LEUKOCYTE AP | Details ... | 19 | -0.6230631 | -1.7228842 | 0.006479482 | 0.09538452 | 1 | 531 |
| NITRIC OXIDE METABOLIC PROCESS | NITRIC OXIDE | Details ... | 15 | -0.65519947 | -1.7213454 | 0.008752735 | 0.095654845 | 1 | 1839 |
| ORGANELLAR RIBOSOME | ORGANELLAR | Details ... | 67 | -0.46560308 | -1.7185049 | 0.002583979 | 0.09717457 | 1 | 5033 |
| CELLULAR PROTEIN COMPLEX DISASSEMBLY | CELLULAR PRO | Details ... | 116 | -0.41859972 | -1.7181016 | 0.002941177 | 0.09644015 | 1 | 5033 |
| RESPONSE TO MANGANESE ION | RESPONSE TO | Details ... | 15 | -0.6480533 | -1.7172576 | 0.010141988 | 0.09606824 | 1 | 1597 |
| PROSTANOID METABOLIC PROCESS | PROSTANOID M | Details ... | 22 | -0.58687353 | -1.7170697 | 0.006564552 | 0.09528247 | 1 | 1316 |
| POSITIVE REGULATION OF REACTIVE OXYGEN SPECIES BIOSYNTHETIC | POSITIVE REGU | Details ... | 43 | -0.51004577 | -1.715787 | 0.007263923 | 0.0953731 | 1 | 198 |
| QUINONE METABOLIC PROCESS | QUINONE MET | Details ... | 24 | -0.57202625 | -1.7152073 | 0.006564552 | 0.094879024 | 1 | 3141 |
| REGULATION OF COLLATERAL SPROUTING | REGULATION ( | Details ... | 16 | -0.64614403 | -1.7150682 | 0.006263048 | 0.094015524 | 1 | 2374 |
| MATURATION OF 5 8S RRNA FROM TRICISTRONIC RRNA TRANSCRIPT | MATURATION ( | Details ... | 19 | -0.59864265 | -1.713133 | 0.010729614 | 0.09474787 | 1 | 4716 |
| GROWTH FACTOR ACTIVITY | GROWTH FACTOR ACTIVITY | | 138 | -0.40393028 | -1.7130457 | 0 | 0.09385669 | 1 | 1513 |
| MYELOID LEUKOCYTE MEDIATED IMMUNITY | MYELOID LEUKOCYTE MEDIATE | | 33 | -0.54154813 | -1.7119161 | 0.006410257 | 0.093793936 | 1 | 275 |
| OXIDOREDUCTASE ACTIVITY ACTING ON SINGLE DONORS WITH INCOR | OXIDOREDUCTASE ACTIVITY AC | | 22 | -0.59569514 | -1.7105917 | 0.013274336 | 0.09379449 | 1 | 975 |
| REGULATION OF FATTY ACID BIOSYNTHETIC PROCESS | REGULATION OF FATTY ACID BI | | 33 | -0.53827536 | -1.7097352 | 0.015021459 | 0.0936153 | 1 | 2107 |
| FATTY ACID DERIVATIVE METABOLIC PROCESS | FATTY ACID DERIVATIVE METAB | | 60 | -0.47615 | -1.7089449 | 0.005089059 | 0.093277216 | 1 | 316 |
| FATTY ACID DERIVATIVE BIOSYNTHETIC PROCESS | FATTY ACID DERIVATIVE BIOSYN | | 34 | -0.5260888 | -1.7059797 | 0.007281554 | 0.094668284 | 1 | 216 |
| REGULATION OF VASCULAR ENDOTHELIAL GROWTH FACTOR PRODUC | REGULATION OF VASCULAR EN | | 29 | -0.54771656 | -1.7005769 | 0.006818182 | 0.09857468 | 1 | 1150 |
| RESPONSE TO AUDITORY STIMULUS | RESPONSE TO AUDITORY STIMU | | 22 | -0.578965 | -1.6962916 | 0.010706638 | 0.101665884 | 1 | 456 |
| ACETYLCHOLINE RECEPTOR ACTIVITY | ACETYLCHOLINE RECEPTOR AC | | 26 | -0.56859523 | -1.6951098 | 0.010964912 | 0.10175026 | 1 | 3290 |
| QUATERNARY AMMONIUM GROUP BINDING | QUATERNARY AMMONIUM GRO | | 42 | -0.503441 | -1.6921906 | 0.011709602 | 0.103321455 | 1 | 1860 |
| POSITIVE REGULATION OF ADENYLATE CYCLASE ACTIVITY | POSITIVE REGULATION OF ADEN | | 43 | -0.49576235 | -1.6889213 | 0.006976744 | 0.104945175 | 1 | 922 |
| ORGANELLAR LARGE RIBOSOMAL SUBUNIT | ORGANELLAR LARGE RIBOSOMA | | 29 | -0.5487995 | -1.6877773 | 0.011389522 | 0.104930736 | 1 | 5412 |
| PTERIDINE CONTAINING COMPOUND BIOSYNTHETIC PROCESS | PTERIDINE CONTAINING COMPO | | 16 | -0.6346818 | -1.6873835 | 0.012526096 | 0.10433505 | 1 | 3247 |
| MUSCLE CELL CELLULAR HOMEOSTASIS | MUSCLE CELL CELLULAR HOME | | 18 | -0.61325896 | -1.6870475 | 0.013186813 | 0.10376585 | 1 | 16 |
| UNSATURATED FATTY ACID BIOSYNTHETIC PROCESS | UNSATURATED FATTY ACID BIO | | 41 | -0.49552295 | -1.686357 | 0.002427185 | 0.10348446 | 1 | 216 |
| TRNA PROCESSING | TRNA PROCESSING | | 89 | -0.42754707 | -1.6849967 | 0.005434783 | 0.10371124 | 1 | 3736 |
| DEFENSE RESPONSE TO GRAM NEGATIVE BACTERIUM | DEFENSE RESPONSE TO GRAM | | 28 | -0.54311615 | -1.6814405 | 0.013888889 | 0.10608895 | 1 | 349 |
| POSITIVE REGULATION OF LYASE ACTIVITY | POSITIVE REGULATION OF LYAS | | 54 | -0.4726829 | -1.6809183 | 0.007407407 | 0.10574077 | 1 | 922 |
| TRNA SPECIFIC RIBONUCLEASE ACTIVITY | TRNA SPECIFIC RIBONUCLEASE | | 15 | -0.65101033 | -1.6801977 | 0.008230452 | 0.1054935 | 1 | 3590 |
| REGULATION OF TYROSINE PHOSPHORYLATION OF STAT PROTEIN | REGULATION OF TYROSINE PHO | | 58 | -0.4617476 | -1.6759759 | 0.004950495 | 0.10805049 | 1 | 1359 |
| EATING BEHAVIOR | EATING BEHAVIOR | | 27 | -0.54658437 | -1.6738708 | 0.016587678 | 0.109152466 | 1 | 917 |
| CYTOSOLIC LARGE RIBOSOMAL SUBUNIT | CYTOSOLIC LARGE RIBOSOMAL | | 54 | -0.46377632 | -1.6736087 | 0.007125891 | 0.10852884 | 1 | 7455 |
| TRNA METABOLIC PROCESS | TRNA METABOLIC PROCESS | | 146 | -0.39484602 | -1.6662222 | 0 | 0.11458175 | 1 | 3736 |
| CYTOKINE ACTIVITY | CYTOKINE ACTIVITY | | 148 | -0.39194697 | -1.6632221 | 0 | 0.11651573 | 1 | 1088 |
| GLUTATHIONE METABOLIC PROCESS | GLUTATHIONE METABOLIC PRO | | 51 | -0.46978927 | -1.6630397 | 0.004938272 | 0.115741596 | 1 | 4241 |
| REGULATION OF CELLULAR AMINO ACID METABOLIC PROCESS | REGULATION OF CELLULAR AMI | | 61 | -0.45803833 | -1.6625749 | 0.004842615 | 0.115270555 | 1 | 5930 |
| FEEDING BEHAVIOR | FEEDING BEHAVIOR | | 78 | -0.44435227 | -1.662473 | 0 | 0.11442946 | 1 | 917 |

| | | | | | | | | |
|---|---|---|---|---|---|---|---|---|
| NEGATIVE REGULATION OF CIRCADIAN RHYTHM | NEGATIVE REGULATION OF CIR | 15 | -0.6286619 | -1.6611662 | 0.008196721 | 0.11481393 | 1 | 1359 |
| RESPONSE TO DIETARY EXCESS | RESPONSE TO DIETARY EXCES | 20 | -0.5893722 | -1.6533198 | 0.015384615 | 0.12180893 | 1 | 900 |
| SMOOTH MUSCLE CONTRACTION | SMOOTH MUSCLE CONTRACTI | 44 | -0.47343296 | -1.653036 | 0.007194245 | 0.12105345 | 1 | 814 |
| POSITIVE REGULATION OF REACTIVE OXYGEN SPECIES METABOLIC PR | POSITIVE REGULATION OF REA | 75 | -0.43552506 | -1.652058 | 0.002617801 | 0.12109735 | 1 | 598 |
| REGULATION OF NITRIC OXIDE BIOSYNTHETIC PROCESS | REGULATION OF NITRIC OXIDE | 48 | -0.47477174 | -1.6506666 | 0 | 0.121520564 | 1 | 198 |
| CLEAVAGE INVOLVED IN RRNA PROCESSING | CLEAVAGE INVOLVED IN RRNA | 18 | -0.5905455 | -1.6486639 | 0.012765957 | 0.122551434 | 1 | 4716 |
| POSITIVE REGULATION OF CD4 POSITIVE ALPHA BETA T CELL ACTIVATI | POSITIVE REGULATION OF CD4 | 23 | -0.5691609 | -1.645629 | 0.009433962 | 0.12467408 | 1 | 373 |
| BRANCHING INVOLVED IN SALIVARY GLAND MORPHOGENESIS | BRANCHING INVOLVED IN SALIV | 16 | -0.62353224 | -1.6439987 | 0.018367346 | 0.12552686 | 1 | 795 |
| REGULATION OF CD4 POSITIVE ALPHA BETA T CELL ACTIVATION | REGULATION OF CD4 POSITIVE | 30 | -0.52706844 | -1.643717 | 0.01369863 | 0.124831036 | 1 | 373 |
| RIBOSOMAL LARGE SUBUNIT ASSEMBLY | RIBOSOMAL LARGE SUBUNIT AS | 21 | -0.5774823 | -1.6411891 | 0.024096385 | 0.12622117 | 1 | 4181 |
| NEGATIVE REGULATION OF LIPASE ACTIVITY | NEGATIVE REGULATION OF LIPA | 15 | -0.63404745 | -1.6391689 | 0.008714597 | 0.12722947 | 1 | 1385 |
| REGULATION OF AMINE TRANSPORT | REGULATION OF AMINE TRANS | 70 | -0.43591616 | -1.6386166 | 0.005208334 | 0.12688187 | 1 | 1038 |
| REGULATION OF IMMUNOGLOBULIN SECRETION | REGULATION OF IMMUNOGLOB | 16 | -0.62391645 | -1.636391 | 0.01793722 | 0.12847202 | 1 | 691 |
| SPLICEOSOMAL TRI SNRNP COMPLEX | SPLICEOSOMAL TRI SNRNP CO | 25 | -0.53115964 | -1.6363146 | 0.018867925 | 0.12764752 | 1 | 5771 |
| BINDING OF SPERM TO ZONA PELLUCIDA | BINDING OF SPERM TO ZONA P | 23 | -0.5628001 | -1.6347451 | 0.010869565 | 0.12852176 | 1 | 3191 |
| SINGLE FERTILIZATION | SINGLE FERTILIZATION | 67 | -0.4402325 | -1.6337653 | 0.004854369 | 0.12873393 | 1 | 3191 |
| REGULATION OF RESPONSE TO FOOD | REGULATION OF RESPONSE TO | 18 | -0.58248925 | -1.6330615 | 0.025751073 | 0.12855853 | 1 | 310 |
| CELLULAR MODIFIED AMINO ACID BIOSYNTHETIC PROCESS | CELLULAR MODIFIED AMINO AC | 47 | -0.46783745 | -1.6313294 | 0.007246377 | 0.12936743 | 1 | 3247 |
| REGULATION OF MULTICELLULAR ORGANISMAL METABOLIC PROCESS | REGULATION OF MULTICELLUL | 34 | -0.5088341 | -1.6286978 | 0.016470589 | 0.13101038 | 1 | 299 |
| POSITIVE T CELL SELECTION | POSITIVE T CELL SELECTION | 16 | -0.60686964 | -1.6283677 | 0.027600849 | 0.13058516 | 1 | 2 |
| DNA TEMPLATED TRANSCRIPTION ELONGATION | DNA TEMPLATED TRANSCRIPTI | 83 | -0.42352492 | -1.6268209 | 0 | 0.13147178 | 1 | 5690 |
| REGULATION OF TYPE 2 IMMUNE RESPONSE | REGULATION OF TYPE 2 IMMUN | 22 | -0.57637864 | -1.6261991 | 0.013015185 | 0.13114494 | 1 | 1301 |
| AMMONIUM TRANSMEMBRANE TRANSPORTER ACTIVITY | AMMONIUM TRANSMEMBRANE | 21 | -0.56804526 | -1.6256189 | 0.021598272 | 0.13086532 | 1 | 1620 |
| GROWTH FACTOR RECEPTOR BINDING | GROWTH FACTOR RECEPTOR | 112 | -0.39742866 | -1.625026 | 0.005747126 | 0.13072167 | 1 | 1069 |
| REGULATION OF POTASSIUM ION TRANSPORT | REGULATION OF POTASSIUM IO | 79 | -0.4276063 | -1.6245054 | 0 | 0.13035122 | 1 | 2209 |
| HORMONE TRANSPORT | HORMONE TRANSPORT | 70 | -0.43659323 | -1.6206479 | 0 | 0.13350533 | 1 | 319 |
| NIK NF KAPPAB SIGNALING | NIK NF KAPPAB SIGNALING | 73 | -0.42508337 | -1.6174443 | 0.005263158 | 0.13604534 | 1 | 5893 |
| ACETYLCHOLINE BINDING | ACETYLCHOLINE BINDING | 19 | -0.59618646 | -1.6172688 | 0.022779044 | 0.13534378 | 1 | 3290 |
| BLOOD COAGULATION INTRINSIC PATHWAY | BLOOD COAGULATION INTRINS | 16 | -0.6224568 | -1.6152682 | 0.020876827 | 0.13657106 | 1 | 363 |
| CORNIFIED ENVELOPE | CORNIFIED ENVELOPE | 17 | -0.58943444 | -1.6142095 | 0.019480519 | 0.13686062 | 1 | 2297 |
| FIBROBLAST GROWTH FACTOR RECEPTOR BINDING | FIBROBLAST GROWTH FACTOR | 27 | -0.5294301 | -1.6138088 | 0.010845987 | 0.13639387 | 1 | 1286 |
| T CELL ACTIVATION INVOLVED IN IMMUNE RESPONSE | T CELL ACTIVATION INVOLVED I | 37 | -0.50091046 | -1.6131631 | 0 | 0.13616934 | 1 | 287 |
| MYELOID CELL HOMEOSTASIS | MYELOID CELL HOMEOSTASIS | 79 | -0.4252156 | -1.6124978 | 0.011049724 | 0.1360367 | 1 | 887 |
| NEGATIVE REGULATION OF SECRETION | NEGATIVE REGULATION OF SEC | 172 | -0.3727858 | -1.6122775 | 0 | 0.13542643 | 1 | 1359 |
| RIBOSOMAL SMALL SUBUNIT BIOGENESIS | RIBOSOMAL SMALL SUBUNIT BI | 53 | -0.4526313 | -1.610597 | 0.012106538 | 0.1363033 | 1 | 5920 |
| REGULATION OF BLOOD PRESSURE | REGULATION OF BLOOD PRESS | 145 | -0.38046545 | -1.6102413 | 0.002976191 | 0.13581362 | 1 | 900 |
| COFACTOR CATABOLIC PROCESS | COFACTOR CATABOLIC PROCE | 17 | -0.60583204 | -1.607994 | 0.016393442 | 0.13738987 | 1 | 397 |
| U2 SNRNP | U2 SNRNP | 17 | -0.59511024 | -1.6068072 | 0.025695931 | 0.13776828 | 1 | 3660 |
| ER NUCLEUS SIGNALING PATHWAY | ER NUCLEUS SIGNALING PATH | 32 | -0.5070384 | -1.6048927 | 0.019002376 | 0.138942 | 1 | 2310 |
| REGULATION OF REACTIVE OXYGEN SPECIES BIOSYNTHETIC PROCESS | REGULATION OF REACTIVE OX | 59 | -0.45101976 | -1.6048838 | 0.00255102 | 0.1381382 | 1 | 198 |
| POSITIVE REGULATION OF STAT CASCADE | POSITIVE REGULATION OF STAT | 60 | -0.4406852 | -1.6045489 | 0.005037783 | 0.13772956 | 1 | 1359 |
| EXOSOME RNASE COMPLEX | EXOSOME RNASE COMPLEX | 18 | -0.57818687 | -1.6033396 | 0.033783782 | 0.13822304 | 1 | 3708 |
| NUCLEAR TRANSCRIBED MRNA CATABOLIC PROCESS EXONUCLEOLYT | NUCLEAR TRANSCRIBED MRNA | 27 | -0.52313024 | -1.603106 | 0.017699115 | 0.13764825 | 1 | 5285 |

| | | | | | | | | |
|---|---|---|---|---|---|---|---|---|
| NCRNA CATABOLIC PROCESS | NCRNA CATABOLIC PROCESS | 17 | -0.593871 | -1.6019275 | 0.020576132 | 0.1380717 | 1 | 3708 |
| AMMONIUM ION BINDING | AMMONIUM ION BINDING | 60 | -0.43422395 | -1.5990611 | 0.007142857 | 0.14021803 | 1 | 1860 |
| POSITIVE REGULATION OF RESPONSE TO WOUNDING | POSITIVE REGULATION OF RESI | 128 | -0.388692 | -1.5982804 | 0 | 0.14037856 | 1 | 413 |
| REGULATION OF SYSTEM PROCESS | REGULATION OF SYSTEM PROC | 471 | -0.32834715 | -1.5977186 | 0 | 0.14027888 | 1 | 1401 |
| SPERM EGG RECOGNITION | SPERM EGG RECOGNITION | 25 | -0.53195333 | -1.5963802 | 0.027083334 | 0.14095286 | 1 | 5018 |
| RESPONSE TO CADMIUM ION | RESPONSE TO CADMIUM ION | 34 | -0.49306765 | -1.5960038 | 0.009708738 | 0.14052154 | 1 | 1654 |
| MOTILE CILIUM | MOTILE CILIUM | 71 | -0.41637227 | -1.5956029 | 0.007633588 | 0.14020576 | 1 | 3097 |
| RNA POLYMERASE ACTIVITY | RNA POLYMERASE ACTIVITY | 38 | -0.49662656 | -1.5951614 | 0.015189873 | 0.13996711 | 1 | 5690 |
| SNORNA BINDING | SNORNA BINDING | 26 | -0.53463733 | -1.594063 | 0.014799154 | 0.1404181 | 1 | 6050 |
| LOW DENSITY LIPOPROTEIN PARTICLE RECEPTOR BINDING | LOW DENSITY LIPOPROTEIN PA | 16 | -0.59347206 | -1.5927765 | 0.026258206 | 0.14101017 | 1 | 1860 |
| PROSTANOID BIOSYNTHETIC PROCESS | PROSTANOID BIOSYNTHETIC PI | 16 | -0.594672 | -1.5922258 | 0.03285421 | 0.14076108 | 1 | 1316 |
| REGULATION OF SYNAPTIC TRANSMISSION GLUTAMATERGIC | REGULATION OF SYNAPTIC TRA | 47 | -0.4545024 | -1.5918436 | 0.010050251 | 0.14042085 | 1 | 598 |
| OVARIAN FOLLICLE DEVELOPMENT | OVARIAN FOLLICLE DEVELOPME | 51 | -0.45193514 | -1.5914117 | 0.014423077 | 0.14018342 | 1 | 1239 |
| RIBONUCLEASE ACTIVITY | RIBONUCLEASE ACTIVITY | 75 | -0.42091265 | -1.5893621 | 0.005361931 | 0.14168935 | 1 | 4930 |
| POSITIVE REGULATION OF LIGASE ACTIVITY | POSITIVE REGULATION OF LIGA | 101 | -0.39748314 | -1.5874932 | 0 | 0.14287776 | 1 | 5893 |
| POSITIVE REGULATION OF NEUROTRANSMITTER TRANSPORT | POSITIVE REGULATION OF NEU | 15 | -0.60377467 | -1.5867558 | 0.014314928 | 0.14304805 | 1 | 1359 |
| NEGATIVE REGULATION OF POTASSIUM ION TRANSPORT | NEGATIVE REGULATION OF POT | 29 | -0.52382517 | -1.5860679 | 0.034632035 | 0.14311473 | 1 | 1782 |
| CELLULAR RESPONSE TO FLUID SHEAR STRESS | CELLULAR RESPONSE TO FLUID | 18 | -0.5846185 | -1.5859574 | 0.029227557 | 0.14245403 | 1 | 1639 |
| FEMALE GAMETE GENERATION | FEMALE GAMETE GENERATION | 66 | -0.42609072 | -1.5857458 | 0.010282776 | 0.1418895 | 1 | 2444 |
| TRIGLYCERIDE CATABOLIC PROCESS | TRIGLYCERIDE CATABOLIC PRO | 21 | -0.56020814 | -1.5855138 | 0.024229076 | 0.14142999 | 1 | 1860 |
| REGULATION OF URINE VOLUME | REGULATION OF URINE VOLUM | 20 | -0.5639464 | -1.5825925 | 0.020361992 | 0.14379206 | 1 | 1359 |
| REGULATION OF ENDOPLASMIC RETICULUM STRESS INDUCED INTRIN | REGULATION OF ENDOPLASMIC | 26 | -0.5308047 | -1.5815501 | 0.02764977 | 0.14410391 | 1 | 1753 |
| OVULATION CYCLE PROCESS | OVULATION CYCLE PROCESS | 76 | -0.4114712 | -1.5788108 | 0.010526316 | 0.14632742 | 1 | 1239 |
| U12 TYPE SPLICEOSOMAL COMPLEX | U12 TYPE SPLICEOSOMAL COMI | 24 | -0.53203005 | -1.5753808 | 0.019955654 | 0.14938174 | 1 | 3660 |
| RESPONSE TO PAIN | RESPONSE TO PAIN | 26 | -0.52137536 | -1.5750811 | 0.025229357 | 0.14895347 | 1 | 1215 |
| STAT CASCADE | STAT CASCADE | 41 | -0.47006744 | -1.5745685 | 0.01724138 | 0.14872192 | 1 | 1225 |
| G PROTEIN COUPLED AMINE RECEPTOR ACTIVITY | G PROTEIN COUPLED AMINE RE | 33 | -0.49774063 | -1.5713376 | 0.025943397 | 0.15160136 | 1 | 3570 |
| CARBOHYDRATE KINASE ACTIVITY | CARBOHYDRATE KINASE ACTIV | 17 | -0.58471954 | -1.5704219 | 0.025806451 | 0.1517331 | 1 | 3635 |
| MITOCHONDRIAL RESPIRATORY CHAIN COMPLEX ASSEMBLY | MITOCHONDRIAL RESPIRATOR | 59 | -0.43436483 | -1.5685818 | 0.02189781 | 0.15316984 | 1 | 7283 |
| VASODILATION | VASODILATION | 25 | -0.52359414 | -1.5659082 | 0.018099548 | 0.15555234 | 1 | 236 |
| NEGATIVE REGULATION OF LIPID STORAGE | NEGATIVE REGULATION OF LIPI | 17 | -0.58307207 | -1.5657965 | 0.018907564 | 0.1549042 | 1 | 299 |
| CARBOHYDRATE PHOSPHORYLATION | CARBOHYDRATE PHOSPHORYL | 18 | -0.5642845 | -1.5629354 | 0.031712472 | 0.15734911 | 1 | 4008 |
| POSITIVE REGULATION OF BEHAVIOR | POSITIVE REGULATION OF BEH | 20 | -0.55216634 | -1.5621771 | 0.03267974 | 0.15745832 | 1 | 1359 |
| POSITIVE REGULATION OF FATTY ACID BIOSYNTHETIC PROCESS | POSITIVE REGULATION OF FAT | 17 | -0.57834524 | -1.5612104 | 0.03267974 | 0.15776668 | 1 | 2540 |
| POSITIVE REGULATION OF ACUTE INFLAMMATORY RESPONSE | POSITIVE REGULATION OF ACU | 25 | -0.5271728 | -1.5610977 | 0.031042129 | 0.15712583 | 1 | 413 |
| NEGATIVE REGULATION OF PEPTIDASE ACTIVITY | NEGATIVE REGULATION OF PEF | 177 | -0.36526406 | -1.559287 | 0 | 0.15845431 | 1 | 1148 |
| PRERIBOSOME LARGE SUBUNIT PRECURSOR | PRERIBOSOME LARGE SUBUNIT | 17 | -0.5781355 | -1.5587263 | 0.029535865 | 0.15838346 | 1 | 3389 |
| CYTOSOLIC RIBOSOME | CYTOSOLIC RIBOSOME | 101 | -0.3920506 | -1.5578437 | 0.007936508 | 0.15861148 | 1 | 7455 |
| DEFENSE RESPONSE TO BACTERIUM | DEFENSE RESPONSE TO BACTE | 116 | -0.38081437 | -1.5575876 | 0.003003003 | 0.15820447 | 1 | 1673 |
| RRNA BINDING | RRNA BINDING | 53 | -0.43375117 | -1.5560036 | 0.016786572 | 0.15931103 | 1 | 5089 |
| NEUROMUSCULAR SYNAPTIC TRANSMISSION | NEUROMUSCULAR SYNAPTIC T | 26 | -0.51192206 | -1.5529352 | 0.035955057 | 0.16224465 | 1 | 867 |
| NEUTRAL LIPID CATABOLIC PROCESS | NEUTRAL LIPID CATABOLIC PRO | 25 | -0.5175395 | -1.552355 | 0.034136545 | 0.16214025 | 1 | 1860 |
| REGULATION OF HORMONE SECRETION | REGULATION OF HORMONE SE | 239 | -0.34401748 | -1.5522618 | 0 | 0.16145867 | 1 | 2412 |

| | | | | | | | | |
|---|---|---|---|---|---|---|---|---|
| REGULATION OF PROTEIN IMPORT INTO NUCLEUS TRANSLOCATION | REGULATION OF PROTEIN IMPO | 19 | -0.5631081 | -1.5520402 | 0.047210302 | 0.1609801 | 1 | 541 |
| WATER TRANSPORT | WATER TRANSPORT | 18 | -0.5436702 | -1.5511686 | 0.06511628 | 0.16127412 | 1 | 2153 |
| RNA PHOSPHODIESTER BOND HYDROLYSIS | RNA PHOSPHODIESTER BOND H | 97 | -0.39572355 | -1.5497491 | 0.005540166 | 0.16221403 | 1 | 4930 |
| NEGATIVE REGULATION OF INTERFERON GAMMA PRODUCTION | NEGATIVE REGULATION OF INTI | 24 | -0.51790345 | -1.5490801 | 0.027459955 | 0.1623412 | 1 | 2961 |
| RIBOSOMAL LARGE SUBUNIT BIOGENESIS | RIBOSOMAL LARGE SUBUNIT BI | 44 | -0.4594868 | -1.5490752 | 0.013513514 | 0.16160087 | 1 | 5071 |
| PHASIC SMOOTH MUSCLE CONTRACTION | PHASIC SMOOTH MUSCLE CON | 15 | -0.5848662 | -1.5470359 | 0.044444446 | 0.1632779 | 1 | 716 |
| ANAPHASE PROMOTING COMPLEX DEPENDENT CATABOLIC PROCESS | ANAPHASE PROMOTING COMPL | 73 | -0.40845042 | -1.5441308 | 0.00530504 | 0.16619964 | 1 | 7670 |
| PROTEIN HOMOTETRAMERIZATION | PROTEIN HOMOTETRAMERIZAT | 55 | -0.4339742 | -1.5435418 | 0.015306123 | 0.16611451 | 1 | 4241 |
| POSITIVE REGULATION OF FATTY ACID METABOLIC PROCESS | POSITIVE REGULATION OF FATT | 33 | -0.47876945 | -1.5426396 | 0.01978022 | 0.16635132 | 1 | 2107 |
| RESPONSE TO HEAT | RESPONSE TO HEAT | 77 | -0.4095533 | -1.5418661 | 0.016853932 | 0.16659059 | 1 | 1590 |
| CYTOKINE RECEPTOR BINDING | CYTOKINE RECEPTOR BINDING | 202 | -0.34807688 | -1.5379907 | 0 | 0.17050132 | 1 | 1088 |
| SPLICEOSOMAL SNRNP ASSEMBLY | SPLICEOSOMAL SNRNP ASSEMB | 34 | -0.4754643 | -1.5379537 | 0.021413276 | 0.16977246 | 1 | 4322 |
| TETRAPYRROLE METABOLIC PROCESS | TETRAPYRROLE METABOLIC PR | 53 | -0.44046834 | -1.5346752 | 0.016548464 | 0.17310183 | 1 | 3591 |
| NUCLEASE ACTIVITY | NUCLEASE ACTIVITY | 157 | -0.3662824 | -1.5335778 | 0.003030303 | 0.17373148 | 1 | 4957 |
| POSITIVE REGULATION OF INTERLEUKIN 6 PRODUCTION | POSITIVE REGULATION OF INTE | 56 | -0.43621257 | -1.5320773 | 0.014457831 | 0.17493683 | 1 | 319 |
| INNER MITOCHONDRIAL MEMBRANE PROTEIN COMPLEX | INNER MITOCHONDRIAL MEMBI | 93 | -0.38588896 | -1.5320398 | 0.010526316 | 0.17421031 | 1 | 7329 |
| HEME BIOSYNTHETIC PROCESS | HEME BIOSYNTHETIC PROCESS | 20 | -0.54336375 | -1.5315003 | 0.033264033 | 0.17422497 | 1 | 5772 |
| U5 SNRNP | U5 SNRNP | 16 | -0.57524556 | -1.5302283 | 0.046025105 | 0.17492884 | 1 | 3660 |
| ACTIVATION OF ADENYLATE CYCLASE ACTIVITY | ACTIVATION OF ADENYLATE CY | 36 | -0.4672132 | -1.5300931 | 0.029612755 | 0.1743691 | 1 | 2369 |
| RESPONSE TO TUMOR NECROSIS FACTOR | RESPONSE TO TUMOR NECROS | 182 | -0.34968266 | -1.527638 | 0.003257329 | 0.17670485 | 1 | 1341 |
| UNSATURATED FATTY ACID METABOLIC PROCESS | UNSATURATED FATTY ACID ME | 71 | -0.40398467 | -1.5274428 | 0.010471204 | 0.17615965 | 1 | 316 |
| MITOCHONDRIAL RESPIRATORY CHAIN COMPLEX I BIOGENESIS | MITOCHONDRIAL RESPIRATORY | 48 | -0.44593737 | -1.526048 | 0.029339854 | 0.17707443 | 1 | 7283 |
| REGULATION OF APPETITE | REGULATION OF APPETITE | 22 | -0.5292496 | -1.5251846 | 0.046413504 | 0.17741995 | 1 | 310 |
| SIGNAL RELEASE | SIGNAL RELEASE | 154 | -0.36110792 | -1.5248322 | 0 | 0.17714715 | 1 | 1753 |
| POSITIVE REGULATION OF ERK1 AND ERK2 CASCADE | POSITIVE REGULATION OF ERK1 | 133 | -0.366752 | -1.522887 | 0.006116208 | 0.17876211 | 1 | 887 |
| TETRAPYRROLE BIOSYNTHETIC PROCESS | TETRAPYRROLE BIOSYNTHETIC | 26 | -0.5052387 | -1.5222982 | 0.029478459 | 0.17886777 | 1 | 3430 |
| CALCIUM ION REGULATED EXOCYTOSIS OF NEUROTRANSMITTER | CALCIUM ION REGULATED EXOC | 28 | -0.49877366 | -1.5220474 | 0.03524229 | 0.17845209 | 1 | 2246 |
| PROTEASOME COMPLEX | PROTEASOME COMPLEX | 70 | -0.40572596 | -1.5220261 | 0.017676767 | 0.17774692 | 1 | 7481 |
| RESPONSE TO ANTIBIOTIC | RESPONSE TO ANTIBIOTIC | 43 | -0.4493413 | -1.5186414 | 0.009367681 | 0.18140796 | 1 | 316 |
| ESTROUS CYCLE | ESTROUS CYCLE | 18 | -0.5489115 | -1.5183762 | 0.050955415 | 0.18103981 | 1 | 1081 |
| CELL REDOX HOMEOSTASIS | CELL REDOX HOMEOSTASIS | 54 | -0.42965674 | -1.5171634 | 0.02173913 | 0.18186167 | 1 | 352 |
| PEPTIDE TRANSPORT | PEPTIDE TRANSPORT | 66 | -0.4128453 | -1.5167581 | 0.012285012 | 0.18158329 | 1 | 1753 |
| MITOCHONDRIAL PROTEIN COMPLEX | MITOCHONDRIAL PROTEIN COM | 119 | -0.37327826 | -1.5149662 | 0.00617284 | 0.18314807 | 1 | 7369 |
| CELLULAR RESPONSE TO ALCOHOL | CELLULAR RESPONSE TO ALCO | 105 | -0.38075763 | -1.5142857 | 0.008333334 | 0.1834359 | 1 | 1128 |
| TRANSFERASE ACTIVITY TRANSFERRING AMINO ACYL GROUPS | TRANSFERASE ACTIVITY TRANS | 15 | -0.5744686 | -1.5133828 | 0.04 | 0.18381685 | 1 | 884 |
| REGULATION OF DIGESTIVE SYSTEM PROCESS | REGULATION OF DIGESTIVE SYS | 33 | -0.47896248 | -1.513037 | 0.029411765 | 0.18351287 | 1 | 2667 |
| RESPONSE TO MOLECULE OF BACTERIAL ORIGIN | RESPONSE TO MOLECULE OF B | 280 | -0.32937184 | -1.5129856 | 0 | 0.18283932 | 1 | 1153 |
| OXYGEN BINDING | OXYGEN BINDING | 24 | -0.5129608 | -1.5128042 | 0.030042918 | 0.18238054 | 1 | 2184 |
| SENSORY PERCEPTION OF PAIN | SENSORY PERCEPTION OF PAIN | 69 | -0.4025032 | -1.5122916 | 0.026666667 | 0.1823285 | 1 | 1505 |
| ADENYLATE CYCLASE ACTIVATING G PROTEIN COUPLED RECEPTOR SI | ADENYLATE CYCLASE ACTIVATI | 54 | -0.43051577 | -1.5121413 | 0.020512821 | 0.1817751 | 1 | 900 |
| REGULATION OF KERATINOCYTE PROLIFERATION | REGULATION OF KERATINOCYT | 23 | -0.508729 | -1.5119845 | 0.030567685 | 0.18129219 | 1 | 1664 |
| RESPONSE TO HYDROGEN PEROXIDE | RESPONSE TO HYDROGEN PER | 100 | -0.37693194 | -1.5110351 | 0.005617978 | 0.18181239 | 1 | 216 |
| MATERNAL PROCESS INVOLVED IN FEMALE PREGNANCY | MATERNAL PROCESS INVOLVED | 53 | -0.42667273 | -1.5107136 | 0.017632242 | 0.18148367 | 1 | 983 |

| | | | | | | | | |
|---|---|---|---|---|---|---|---|---|
| PSEUDOURIDINE SYNTHESIS | PSEUDOURIDINE SYNTHESIS | 15 | -0.581652 | -1.5093158 | 0.042372882 | 0.18249226 | 1 | 2986 |
| NEGATIVE REGULATION OF NEURON DEATH | NEGATIVE REGULATION OF NEU | 152 | -0.3553655 | -1.5091907 | 0 | 0.1819384 | 1 | 2000 |
| INTRINSIC APOPTOTIC SIGNALING PATHWAY | INTRINSIC APOPTOTIC SIGNALI | 134 | -0.364631 | -1.508536 | 0.005780347 | 0.18210009 | 1 | 1597 |
| RNA PHOSPHODIESTER BOND HYDROLYSIS ENDONUCLEOLYTIC | RNA PHOSPHODIESTER BOND H | 52 | -0.42352653 | -1.5078399 | 0.024271844 | 0.18224119 | 1 | 4909 |
| REGULATION OF AMINO ACID TRANSPORT | REGULATION OF AMINO ACID T | 25 | -0.4915542 | -1.5073988 | 0.05463183 | 0.18216921 | 1 | 917 |
| REGULATION OF CATECHOLAMINE SECRETION | REGULATION OF CATECHOLAM | 42 | -0.4397539 | -1.5073323 | 0.025171624 | 0.18156698 | 1 | 1315 |
| HISTONE MRNA METABOLIC PROCESS | HISTONE MRNA METABOLIC PR | 25 | -0.5023693 | -1.5062847 | 0.0247191 | 0.1821372 | 1 | 5633 |
| REGULATION OF INFLAMMATORY RESPONSE | REGULATION OF INFLAMMATOR | 240 | -0.33373713 | -1.5062556 | 0.003533569 | 0.18148285 | 1 | 1359 |
| PROTEIN TARGETING TO MITOCHONDRION | PROTEIN TARGETING TO MITOC | 44 | -0.43842366 | -1.5050302 | 0.025906736 | 0.18225285 | 1 | 6509 |
| INTRINSIC COMPONENT OF MITOCHONDRIAL INNER MEMBRANE | INTRINSIC COMPONENT OF MIT | 16 | -0.5573384 | -1.5046825 | 0.06508876 | 0.18203111 | 1 | 4339 |
| CELLULAR RESPONSE TO KETONE | CELLULAR RESPONSE TO KETO | 69 | -0.4005081 | -1.5040661 | 0.012953368 | 0.18211387 | 1 | 459 |
| LIGASE ACTIVITY FORMING CARBON NITROGEN BONDS | LIGASE ACTIVITY FORMING CAR | 45 | -0.4421545 | -1.5025651 | 0.023316063 | 0.18338753 | 1 | 3552 |
| REGULATION OF VASOCONSTRICTION | REGULATION OF VASOCONSTR | 61 | -0.41895628 | -1.5004456 | 0.025773196 | 0.18529692 | 1 | 2883 |
| CLATHRIN BINDING | CLATHRIN BINDING | 52 | -0.42702314 | -1.5000782 | 0.029339854 | 0.18510163 | 1 | 1247 |
| POSITIVE REGULATION OF HORMONE SECRETION | POSITIVE REGULATION OF HOR | 107 | -0.37145925 | -1.498013 | 0.008902078 | 0.18706751 | 1 | 2412 |
| OXIDOREDUCTASE ACTIVITY ACTING ON NAD P H QUINONE OR SIMILAI | OXIDOREDUCTASE ACTIVITY AC | 46 | -0.4393343 | -1.4960595 | 0.042128604 | 0.18885854 | 1 | 7283 |
| NCRNA 3 END PROCESSING | NCRNA 3 END PROCESSING | 18 | -0.5428576 | -1.4953035 | 0.054545455 | 0.1891 | 1 | 5579 |
| POSITIVE REGULATION OF ANION TRANSPORT | POSITIVE REGULATION OF ANIC | 53 | -0.41383368 | -1.4939908 | 0.02238806 | 0.1901258 | 1 | 413 |
| NEGATIVE REGULATION OF CYSTEINE TYPE ENDOPEPTIDASE ACTIVITY | NEGATIVE REGULATION OF CYS | 76 | -0.392046 | -1.4939059 | 0.010752688 | 0.18951656 | 1 | 1310 |
| THREONINE TYPE PEPTIDASE ACTIVITY | THREONINE TYPE PEPTIDASE A | 18 | -0.5419048 | -1.492682 | 0.057815846 | 0.19043736 | 1 | 5887 |
| REGULATION OF MACROPHAGE DIFFERENTIATION | REGULATION OF MACROPHAGE | 19 | -0.5425801 | -1.4913406 | 0.051282052 | 0.1915715 | 1 | 621 |
| LYMPHOCYTE ACTIVATION INVOLVED IN IMMUNE RESPONSE | LYMPHOCYTE ACTIVATION INVO | 70 | -0.40071315 | -1.4903527 | 0.013227513 | 0.1921314 | 1 | 1359 |
| OVULATION | OVULATION | 16 | -0.5681397 | -1.4900314 | 0.06431536 | 0.19183323 | 1 | 908 |
| SALIVARY GLAND DEVELOPMENT | SALIVARY GLAND DEVELOPMEN | 31 | -0.4819838 | -1.4893404 | 0.03803132 | 0.19208884 | 1 | 937 |
| GAMMA TUBULIN COMPLEX | GAMMA TUBULIN COMPLEX | 15 | -0.57429147 | -1.4893016 | 0.043392505 | 0.19148552 | 1 | 4428 |
| CIRCULATORY SYSTEM PROCESS | CIRCULATORY SYSTEM PROCE | 326 | -0.32090357 | -1.4887723 | 0 | 0.19147106 | 1 | 2338 |
| RESPONSE TO INTERLEUKIN 4 | RESPONSE TO INTERLEUKIN 4 | 28 | -0.48238257 | -1.4878048 | 0.049676027 | 0.1919718 | 1 | 465 |
| REGULATION OF REACTIVE OXYGEN SPECIES METABOLIC PROCESS | REGULATION OF REACTIVE OXY | 132 | -0.35306308 | -1.4878005 | 0.016666668 | 0.19130246 | 1 | 1658 |
| PALMITOYLTRANSFERASE ACTIVITY | PALMITOYLTRANSFERASE ACTI | 28 | -0.48619598 | -1.487645 | 0.041942604 | 0.1908322 | 1 | 2869 |
| SNARE BINDING | SNARE BINDING | 110 | -0.36425754 | -1.4865952 | 0.018766755 | 0.19155277 | 1 | 2389 |
| RESPONSE TO ATP | RESPONSE TO ATP | 29 | -0.47075677 | -1.486306 | 0.039337475 | 0.19125162 | 1 | 716 |
| U4 U6 X U5 TRI SNRNP COMPLEX | U4 U6 X U5 TRI SNRNP COMPLE | 20 | -0.5397503 | -1.485856 | 0.059196617 | 0.19129172 | 1 | 5633 |
| SYMPATHETIC NERVOUS SYSTEM DEVELOPMENT | SYMPATHETIC NERVOUS SYSTE | 18 | -0.55113745 | -1.4858037 | 0.06198347 | 0.19068505 | 1 | 4342 |
| LEUKOCYTE HOMEOSTASIS | LEUKOCYTE HOMEOSTASIS | 52 | -0.4181489 | -1.4855887 | 0.030075189 | 0.19029953 | 1 | 2552 |
| ALDITOL METABOLIC PROCESS | ALDITOL METABOLIC PROCESS | 16 | -0.5502645 | -1.4855599 | 0.05788423 | 0.18970095 | 1 | 2980 |
| OVULATION CYCLE | OVULATION CYCLE | 99 | -0.37328175 | -1.4852479 | 0.0078125 | 0.18949583 | 1 | 1239 |
| STRUCTURAL CONSTITUENT OF RIBOSOME | STRUCTURAL CONSTITUENT O | 192 | -0.34091878 | -1.4844387 | 0 | 0.18993105 | 1 | 7457 |
| REGULATION OF TRANSFORMING GROWTH FACTOR BETA PRODUCTI | REGULATION OF TRANSFORMI | 21 | -0.52256995 | -1.482731 | 0.04112554 | 0.19133961 | 1 | 4265 |
| TRANSCRIPTION FACTOR TFIID COMPLEX | TRANSCRIPTION FACTOR TFIID | 19 | -0.53149444 | -1.4818634 | 0.047817048 | 0.19184008 | 1 | 3023 |
| POSITIVE REGULATION OF IMMUNE EFFECTOR PROCESS | POSITIVE REGULATION OF IMMU | 130 | -0.3567113 | -1.4795928 | 0.002976191 | 0.19405459 | 1 | 771 |
| ENDOCRINE PROCESS | ENDOCRINE PROCESS | 38 | -0.44895583 | -1.4793258 | 0.049411766 | 0.19377719 | 1 | 2174 |
| REGULATION OF SYSTEMIC ARTERIAL BLOOD PRESSURE BY HORMONI | REGULATION OF SYSTEMIC ART | 30 | -0.4736552 | -1.47694 | 0.048034936 | 0.1962737 | 1 | 2174 |
| DECIDUALIZATION | DECIDUALIZATION | 20 | -0.5194038 | -1.4752 | 0.07594936 | 0.1978949 | 1 | 621 |

| | | | | | | | | |
|---|---|---|---|---|---|---|---|---|
| CARBOHYDRATE TRANSPORT | CARBOHYDRATE TRANSPORT | 80 | -0.37704593 | -1.4748414 | 0.018324608 | 0.1976951 | 1 | 2681 |
| NEUTRAL LIPID METABOLIC PROCESS | NEUTRAL LIPID METABOLIC PRO | 76 | -0.38658246 | -1.4742677 | 0.015957447 | 0.19774294 | 1 | 2010 |
| POSITIVE REGULATION OF SMOOTH MUSCLE CELL PROLIFERATION | POSITIVE REGULATION OF SMO | 56 | -0.41435722 | -1.4736433 | 0.035885166 | 0.19787605 | 1 | 7 |
| AMIDE TRANSPORT | AMIDE TRANSPORT | 86 | -0.37916154 | -1.4723507 | 0.026954178 | 0.19893293 | 1 | 1753 |
| POSITIVE REGULATION OF PRODUCTION OF MOLECULAR MEDIATOR O | POSITIVE REGULATION OF PRO | 55 | -0.4116909 | -1.4718888 | 0.021791767 | 0.1988569 | 1 | 1444 |
| POSITIVE REGULATION OF SYNAPTIC TRANSMISSION | POSITIVE REGULATION OF SYN | 101 | -0.36916587 | -1.4706763 | 0.008474576 | 0.19986254 | 1 | 1578 |
| REGULATION OF SYSTEMIC ARTERIAL BLOOD PRESSURE MEDIATED BY | REGULATION OF SYSTEMIC ART | 40 | -0.44337496 | -1.4702408 | 0.0647482 | 0.1998278 | 1 | 5670 |
| ENDONUCLEASE ACTIVITY | ENDONUCLEASE ACTIVITY | 95 | -0.37676018 | -1.4698186 | 0.012658228 | 0.19981 | 1 | 4756 |
| REGULATION OF CIRCADIAN RHYTHM | REGULATION OF CIRCADIAN RH | 92 | -0.37869564 | -1.4671913 | 0.015625 | 0.20259586 | 1 | 1407 |
| TETRAPYRROLE BINDING | TETRAPYRROLE BINDING | 84 | -0.3793695 | -1.4671406 | 0.015228426 | 0.20200098 | 1 | 883 |
| CYTOSOLIC PART | CYTOSOLIC PART | 186 | -0.33493024 | -1.4666361 | 0 | 0.201992 | 1 | 7481 |
| DNA DIRECTED RNA POLYMERASE III COMPLEX | DNA DIRECTED RNA POLYMERA | 16 | -0.5478966 | -1.4656448 | 0.05567452 | 0.20272966 | 1 | 5690 |
| RESPONSE TO TEMPERATURE STIMULUS | RESPONSE TO TEMPERATURE S | 128 | -0.35419306 | -1.4652139 | 0.016574586 | 0.20258796 | 1 | 1590 |
| NEGATIVE REGULATION OF CARBOHYDRATE METABOLIC PROCESS | NEGATIVE REGULATION OF CAR | 41 | -0.4416111 | -1.464601 | 0.03456221 | 0.20272787 | 1 | 1315 |
| MULTICELLULAR ORGANISMAL RESPONSE TO STRESS | MULTICELLULAR ORGANISMAL R | 64 | -0.39967018 | -1.4625052 | 0.024509804 | 0.20488754 | 1 | 1215 |
| INFLAMMATORY RESPONSE | INFLAMMATORY RESPONSE | 339 | -0.30894756 | -1.462265 | 0 | 0.20454146 | 1 | 1084 |
| REGULATION OF NEUROTRANSMITTER TRANSPORT | REGULATION OF NEUROTRANS | 58 | -0.39720213 | -1.461888 | 0.02 | 0.2044358 | 1 | 2332 |
| PROTEASOME ACCESSORY COMPLEX | PROTEASOME ACCESSORY CO | 23 | -0.49786693 | -1.4602864 | 0.06696428 | 0.20589502 | 1 | 7613 |
| ENDORIBONUCLEASE COMPLEX | ENDORIBONUCLEASE COMPLEX | 17 | -0.5443269 | -1.4595349 | 0.057395145 | 0.2062938 | 1 | 3691 |
| BLOOD COAGULATION FIBRIN CLOT FORMATION | BLOOD COAGULATION FIBRIN C | 22 | -0.5074394 | -1.4568717 | 0.057522126 | 0.20938872 | 1 | 363 |
| REGULATION OF CYTOKINE BIOSYNTHETIC PROCESS | REGULATION OF CYTOKINE BIO | 77 | -0.38490415 | -1.4566895 | 0.029333333 | 0.209007 | 1 | 215 |
| REGULATION OF PRODUCTION OF MOLECULAR MEDIATOR OF IMMUNE | REGULATION OF PRODUCTION | 85 | -0.3711027 | -1.4553424 | 0.009925558 | 0.2101326 | 1 | 1444 |
| CYTOPLASMIC TRANSLATION | CYTOPLASMIC TRANSLATION | 38 | -0.4432526 | -1.4531518 | 0.03837472 | 0.21257405 | 1 | 6994 |
| RRNA MODIFICATION | RRNA MODIFICATION | 22 | -0.507285 | -1.4518151 | 0.06521739 | 0.21370053 | 1 | 3881 |
| REGULATION OF ADENYLATE CYCLASE ACTIVITY | REGULATION OF ADENYLATE CY | 64 | -0.3960506 | -1.45106 | 0.029023746 | 0.21404397 | 1 | 1467 |
| INTERMEDIATE FILAMENT ORGANIZATION | INTERMEDIATE FILAMENT ORGA | 19 | -0.52850926 | -1.4508499 | 0.05882353 | 0.2136798 | 1 | 715 |
| SNAP RECEPTOR ACTIVITY | SNAP RECEPTOR ACTIVITY | 33 | -0.45146918 | -1.4497415 | 0.046838406 | 0.2145873 | 1 | 3297 |
| CELLULAR RESPONSE TO HYDROGEN PEROXIDE | CELLULAR RESPONSE TO HYDR | 55 | -0.41078296 | -1.4493854 | 0.04379562 | 0.2145949 | 1 | 216 |
| RESPONSE TO BACTERIUM | RESPONSE TO BACTERIUM | 370 | -0.30817643 | -1.4477462 | 0 | 0.21622553 | 1 | 1153 |
| EXOCRINE SYSTEM DEVELOPMENT | EXOCRINE SYSTEM DEVELOPM | 43 | -0.41932932 | -1.4467467 | 0.040302265 | 0.2168338 | 1 | 937 |
| RESPONSE TO PROTOZOAN | RESPONSE TO PROTOZOAN | 18 | -0.5235125 | -1.446707 | 0.07392197 | 0.21622217 | 1 | 140 |
| AMINO ACID BETAINE METABOLIC PROCESS | AMINO ACID BETAINE METABOL | 18 | -0.54725134 | -1.4463378 | 0.07949791 | 0.21608591 | 1 | 3055 |
| MYELOID LEUKOCYTE MIGRATION | MYELOID LEUKOCYTE MIGRATIO | 73 | -0.38488442 | -1.4457092 | 0.02917772 | 0.21628095 | 1 | 1194 |
| RESPONSE TO EXTRACELLULAR STIMULUS | RESPONSE TO EXTRACELLULAR | 410 | -0.30303368 | -1.442221 | 0 | 0.22055115 | 1 | 1712 |
| REGULATION OF NECROTIC CELL DEATH | REGULATION OF NECROTIC CE | 23 | -0.49624252 | -1.4419107 | 0.07522124 | 0.22035748 | 1 | 1468 |
| REGULATION OF ENDOCRINE PROCESS | REGULATION OF ENDOCRINE P | 44 | -0.42229626 | -1.4413791 | 0.02739726 | 0.22047465 | 1 | 1038 |
| POSITIVE REGULATION OF SECRETION | POSITIVE REGULATION OF SECR | 316 | -0.310219 | -1.441378 | 0 | 0.21982126 | 1 | 2412 |
| RNA CAPPING | RNA CAPPING | 34 | -0.44613087 | -1.4404988 | 0.058252428 | 0.22034411 | 1 | 5723 |
| REGULATION OF LYASE ACTIVITY | REGULATION OF LYASE ACTIVIT | 78 | -0.3792947 | -1.4400505 | 0.031088082 | 0.22040416 | 1 | 922 |
| MULTI MULTICELLULAR ORGANISM PROCESS | MULTI MULTICELLULAR ORGAN | 178 | -0.3349211 | -1.4370419 | 0.006230529 | 0.22397721 | 1 | 1639 |
| NEGATIVE REGULATION OF CYTOKINE SECRETION | NEGATIVE REGULATION OF CYT | 31 | -0.45390552 | -1.4363214 | 0.058441557 | 0.22433162 | 1 | 1359 |
| CELLULAR RESPONSE TO GLUCOSE STARVATION | CELLULAR RESPONSE TO GLUC | 30 | -0.45526192 | -1.4361544 | 0.05491991 | 0.22391376 | 1 | 3972 |
| OXIDOREDUCTASE ACTIVITY ACTING ON PEROXIDE AS ACCEPTOR | OXIDOREDUCTASE ACTIVITY AC | 35 | -0.4403891 | -1.43615 | 0.049763035 | 0.2232646 | 1 | 40 |

| | | | | | | | | | |
|---|---|---|---|---|---|---|---|---|---|
| POSITIVE REGULATION OF NF KAPPAB TRANSCRIPTION FACTOR ACTIV | POSITIVE REGULATION OF NF K | | 112 | -0.354142 | -1.435459 | 0.016 | 0.22362961 | 1 | 1468 |
| POSITIVE REGULATION OF ALPHA BETA T CELL DIFFERENTIATION | POSITIVE REGULATION OF ALPH | | 31 | -0.45109162 | -1.4343241 | 0.058956917 | 0.22453584 | 1 | 2 |
| NEGATIVE REGULATION OF PROTEOLYSIS | NEGATIVE REGULATION OF PRO | | 253 | -0.3157989 | -1.4331231 | 0 | 0.22563861 | 1 | 1310 |
| REGULATION OF ALPHA BETA T CELL DIFFERENTIATION | REGULATION OF ALPHA BETA T | | 37 | -0.43442369 | -1.4322191 | 0.057208236 | 0.22635579 | 1 | 2 |
| POSITIVE REGULATION OF PEPTIDE SECRETION | POSITIVE REGULATION OF PEPT | | 82 | -0.36715126 | -1.4312605 | 0.021447722 | 0.22703332 | 1 | 2412 |
| REGULATION OF DOPAMINE SECRETION | REGULATION OF DOPAMINE SEC | | 22 | -0.49248195 | -1.430936 | 0.06048387 | 0.22685581 | 1 | 867 |
| OXIDOREDUCTASE ACTIVITY ACTING ON PAIRED DONORS WITH INCOR | OXIDOREDUCTASE ACTIVITY AC | | 26 | -0.47041622 | -1.4301897 | 0.06593407 | 0.22718418 | 1 | 1206 |
| MULTI ORGANISM METABOLIC PROCESS | MULTI ORGANISM METABOLIC P | | 128 | -0.34645948 | -1.4299927 | 0.009345794 | 0.22683115 | 1 | 7455 |
| CELLULAR METABOLIC COMPOUND SALVAGE | CELLULAR METABOLIC COMPOU | | 31 | -0.46260047 | -1.4293532 | 0.06418219 | 0.22725613 | 1 | 6576 |
| DIGESTION | DIGESTION | | 100 | -0.35716358 | -1.4273509 | 0.021917809 | 0.22956927 | 1 | 1486 |
| NUCLEOBASE BIOSYNTHETIC PROCESS | NUCLEOBASE BIOSYNTHETIC P | | 17 | -0.5214006 | -1.424214 | 0.07805907 | 0.23370193 | 1 | 3055 |
| REGULATION OF LIGASE ACTIVITY | REGULATION OF LIGASE ACTIVIT | | 119 | -0.3451833 | -1.4221232 | 0.012618297 | 0.23615907 | 1 | 5893 |
| NEGATIVE REGULATION OF CALCIUM ION TRANSPORT | NEGATIVE REGULATION OF CAL | | 44 | -0.421213 | -1.421 | 0.035799522 | 0.23723716 | 1 | 400 |
| INTERMEDIATE FILAMENT BASED PROCESS | INTERMEDIATE FILAMENT BASE | | 37 | -0.42383933 | -1.4203533 | 0.049881235 | 0.2375629 | 1 | 2001 |
| NEGATIVE REGULATION OF BLOOD PRESSURE | NEGATIVE REGULATION OF BLO | | 41 | -0.41764176 | -1.4201684 | 0.055309735 | 0.23712546 | 1 | 3922 |
| REGULATION OF RESPONSE TO WOUNDING | REGULATION OF RESPONSE TO | | 344 | -0.30643508 | -1.4168161 | 0 | 0.24156949 | 1 | 1311 |
| ENDORIBONUCLEASE ACTIVITY PRODUCING 5 PHOSPHOMONOESTERS | ENDORIBONUCLEASE ACTIVITY | | 25 | -0.47995478 | -1.4150153 | 0.0787037 | 0.24379633 | 1 | 4909 |
| RESPONSE TO TRANSITION METAL NANOPARTICLE | RESPONSE TO TRANSITION MET | | 130 | -0.33859673 | -1.4145788 | 0.009118541 | 0.2437639 | 1 | 1277 |
| NUCLEIC ACID PHOSPHODIESTER BOND HYDROLYSIS | NUCLEIC ACID PHOSPHODIESTI | | 208 | -0.32388794 | -1.4141859 | 0 | 0.24365608 | 1 | 5057 |
| EXCRETION | EXCRETION | | 42 | -0.41379026 | -1.4121912 | 0.0520362 | 0.24602704 | 1 | 1706 |
| POSITIVE REGULATION OF DNA REPLICATION | POSITIVE REGULATION OF DNA | | 78 | -0.3629849 | -1.4110427 | 0.025252525 | 0.24713914 | 1 | 229 |
| FOLIC ACID METABOLIC PROCESS | FOLIC ACID METABOLIC PROCE | | 17 | -0.51644874 | -1.4109027 | 0.08898305 | 0.24661696 | 1 | 5705 |
| GLYCOSYL COMPOUND CATABOLIC PROCESS | GLYCOSYL COMPOUND CATABO | | 31 | -0.45353988 | -1.4098713 | 0.073068894 | 0.24746238 | 1 | 3179 |
| LEUKOCYTE CHEMOTAXIS | LEUKOCYTE CHEMOTAXIS | | 91 | -0.35307142 | -1.4085457 | 0.02406417 | 0.24878691 | 1 | 1201 |
| PIGMENT METABOLIC PROCESS | PIGMENT METABOLIC PROCESS | | 52 | -0.4023851 | -1.4080186 | 0.06367925 | 0.24888144 | 1 | 3591 |
| PROTEIN LOCALIZATION TO MITOCHONDRION | PROTEIN LOCALIZATION TO MIT | | 59 | -0.38216123 | -1.4079753 | 0.035545025 | 0.2482795 | 1 | 7082 |
| METAL CLUSTER BINDING | METAL CLUSTER BINDING | | 55 | -0.39306262 | -1.4076259 | 0.066831686 | 0.2480896 | 1 | 7443 |
| POSITIVE REGULATION OF CHEMOKINE PRODUCTION | POSITIVE REGULATION OF CHE | | 40 | -0.42287114 | -1.4073389 | 0.061926607 | 0.2478202 | 1 | 319 |
| POSITIVE REGULATION OF INSULIN SECRETION INVOLVED IN CELLULAR | POSITIVE REGULATION OF INSU | | 23 | -0.488199 | -1.4059256 | 0.09684685 | 0.24928598 | 1 | 4886 |

| Down Regulated in APC^Pirc/++OX in relation to wild-type | | | | | | | | | |
|---|---|---|---|---|---|---|---|---|---|
| NAME | GS<br> follow li | GS DETAILS | SIZE | ES | NES | NOM p-val | FDR q-val | FWER p-val | RANK AT MAX |
| EXTRACELLULAR MATRIX COMPONENT | EXTRACELLUL | Details ... | 111 | 0.6992233 | 2.5978312 | 0 | 0 | 0 | 2354 |
| ENSHEATHMENT OF NEURONS | ENSHEATHME | Details ... | 83 | 0.6816524 | 2.414752 | 0 | 0 | 0 | 1680 |
| PROTEINACEOUS EXTRACELLULAR MATRIX | PROTEINACEO | Details ... | 292 | 0.5758632 | 2.3865523 | 0 | 0 | 0 | 1716 |
| EXTRACELLULAR MATRIX BINDING | EXTRACELLUL | Details ... | 41 | 0.739514 | 2.3541222 | 0 | 0 | 0 | 1697 |
| EXTRACELLULAR MATRIX | EXTRACELLUL | Details ... | 351 | 0.55319816 | 2.3176136 | 0 | 0 | 0 | 2452 |
| COLLAGEN BINDING | COLLAGEN BIN | Details ... | 50 | 0.708326 | 2.3174088 | 0 | 0 | 0 | 1620 |
| REGULATION OF CARTILAGE DEVELOPMENT | REGULATION C | Details ... | 51 | 0.70255125 | 2.3109586 | 0 | 0 | 0 | 2601 |
| COLLAGEN TRIMER | COLLAGEN TR | Details ... | 67 | 0.66677797 | 2.3083568 | 0 | 0 | 0 | 1679 |
| REGULATION OF CHONDROCYTE DIFFERENTIATION | REGULATION C | Details ... | 37 | 0.76406735 | 2.3068252 | 0 | 0 | 0 | 2601 |
| BASEMENT MEMBRANE | BASEMENT ME | Details ... | 81 | 0.6520201 | 2.280848 | 0 | 0 | 0 | 2354 |
| COMPLEX OF COLLAGEN TRIMERS | COMPLEX OF C | Details ... | 22 | 0.83910173 | 2.278998 | 0 | 0 | 0 | 1679 |

| | | | | | | | | | |
|---|---|---|---|---|---|---|---|---|---|
| BONE DEVELOPMENT | BONE DEVELO | Details ... | 141 | 0.5865296 | 2.270413 | 0 | 0 | 0 | 2185 |
| CELL MATRIX ADHESION | CELL MATRIX A | Details ... | 103 | 0.61496246 | 2.2502763 | 0 | 0 | 0 | 2437 |
| EXTRACELLULAR STRUCTURE ORGANIZATION | EXTRACELLUL | Details ... | 260 | 0.5402207 | 2.222778 | 0 | 1.49E-04 | 0.002 | 2473 |
| CELL SUBSTRATE ADHESION | CELL SUBSTRA | Details ... | 144 | 0.5697678 | 2.1820848 | 0 | 3.47E-04 | 0.005 | 2437 |
| CHONDROCYTE DIFFERENTIATION | CHONDROCYT | Details ... | 53 | 0.66284615 | 2.1816912 | 0 | 3.25E-04 | 0.005 | 2139 |
| NEGATIVE REGULATION OF TRANSFORMING GROWTH FACTOR BETA R | NEGATIVE REG | Details ... | 59 | 0.6378764 | 2.155715 | 0 | 4.90E-04 | 0.008 | 1105 |
| SCHWANN CELL DIFFERENTIATION | SCHWANN CE | Details ... | 30 | 0.73784286 | 2.151092 | 0 | 5.81E-04 | 0.01 | 2575 |
| CANONICAL WNT SIGNALING PATHWAY | CANONICAL W | Details ... | 75 | 0.6160004 | 2.1480687 | 0 | 6.04E-04 | 0.011 | 1589 |
| SKELETAL SYSTEM DEVELOPMENT | SKELETAL SYS | Details ... | 396 | 0.49951655 | 2.126146 | 0 | 0.001146836 | 0.021 | 2185 |
| ENDOCHONDRAL BONE MORPHOGENESIS | ENDOCHONDR | Details ... | 41 | 0.6743561 | 2.1220813 | 0 | 0.001192407 | 0.023 | 2161 |
| BONE MORPHOGENESIS | BONE MORPHO | Details ... | 69 | 0.61362857 | 2.1189167 | 0 | 0.001327559 | 0.027 | 2161 |
| CONNECTIVE TISSUE DEVELOPMENT | CONNECTIVE T | Details ... | 166 | 0.5346225 | 2.1020133 | 0 | 0.001723111 | 0.036 | 2185 |
| CARTILAGE DEVELOPMENT | CARTILAGE DE | Details ... | 128 | 0.55716497 | 2.1002471 | 0 | 0.001651315 | 0.036 | 2185 |
| POSITIVE REGULATION OF STEM CELL DIFFERENTIATION | POSITIVE REGU | Details ... | 43 | 0.65838027 | 2.0897753 | 0 | 0.001878162 | 0.043 | 1063 |
| REGULATION OF STEM CELL DIFFERENTIATION | REGULATION O | Details ... | 95 | 0.5726009 | 2.0874763 | 0 | 0.001886995 | 0.045 | 1221 |
| REGULATION OF CELLULAR RESPONSE TO TRANSFORMING GROWTH F | REGULATION O | Details ... | 87 | 0.5764726 | 2.0872536 | 0 | 0.001817106 | 0.045 | 1105 |
| SKELETAL SYSTEM MORPHOGENESIS | SKELETAL SYS | Details ... | 166 | 0.5279275 | 2.0839016 | 0 | 0.001939769 | 0.05 | 2161 |
| PERIPHERAL NERVOUS SYSTEM AXON ENSHEATHMENT | PERIPHERAL N | Details ... | 21 | 0.7728995 | 2.0830863 | 0 | 0.001979958 | 0.053 | 1182 |
| COLLAGEN FIBRIL ORGANIZATION | COLLAGEN FIB | Details ... | 34 | 0.6993348 | 2.0652502 | 0 | 0.002575385 | 0.071 | 2608 |
| MEMBRANE PROTEIN PROTEOLYSIS | MEMBRANE PR | Details ... | 33 | 0.69261014 | 2.0591502 | 0 | 0.002794588 | 0.078 | 1947 |
| SCHWANN CELL DEVELOPMENT | SCHWANN CE | Details ... | 25 | 0.73166883 | 2.0539007 | 0 | 0.003031134 | 0.087 | 1867 |
| REGULATION OF CELLULAR RESPONSE TO GROWTH FACTOR STIMULU | REGULATION O | Details ... | 197 | 0.5097253 | 2.051497 | 0 | 0.003001672 | 0.089 | 1206 |
| SMAD BINDING | SMAD BINDING | Details ... | 62 | 0.6049491 | 2.0484452 | 0 | 0.002975029 | 0.09 | 2476 |
| BETA CATENIN BINDING | BETA CATENIN | Details ... | 73 | 0.5839476 | 2.0358965 | 0 | 0.003485941 | 0.109 | 1734 |
| EXTRACELLULAR MATRIX STRUCTURAL CONSTITUENT | EXTRACELLUL | Details ... | 63 | 0.59945196 | 2.0261571 | 0 | 0.003822845 | 0.123 | 1679 |
| APPENDAGE DEVELOPMENT | APPENDAGE D | Details ... | 133 | 0.5279114 | 2.0248644 | 0 | 0.003775477 | 0.125 | 2802 |
| REGULATION OF SMALL GTPASE MEDIATED SIGNAL TRANSDUCTION | REGULATION O | Details ... | 248 | 0.49366313 | 2.024741 | 0 | 0.003676122 | 0.125 | 2584 |
| GLIAL CELL DIFFERENTIATION | GLIAL CELL DIF | Details ... | 123 | 0.535029 | 2.015629 | 0 | 0.004114042 | 0.138 | 1989 |
| REGULATION OF LAMELLIPODIUM ORGANIZATION | REGULATION O | Details ... | 32 | 0.68065745 | 2.0149183 | 0 | 0.004011191 | 0.138 | 2160 |
| EMBRYONIC EYE MORPHOGENESIS | EMBRYONIC E | Details ... | 31 | 0.677109 | 2.01426 | 0.001795332 | 0.003913357 | 0.138 | 1025 |
| ENDODERM DEVELOPMENT | ENDODERM D | Details ... | 60 | 0.5972082 | 2.0071712 | 0 | 0.004468182 | 0.159 | 1366 |
| REGULATION OF EPITHELIAL TO MESENCHYMAL TRANSITION | REGULATION O | Details ... | 58 | 0.6034728 | 2.0067313 | 0 | 0.004364271 | 0.159 | 1221 |
| SENSORY ORGAN MORPHOGENESIS | SENSORY ORG | Details ... | 206 | 0.49415582 | 2.0053625 | 0 | 0.004525335 | 0.17 | 2409 |
| LAMELLIPODIUM MEMBRANE | LAMELLIPODIU | Details ... | 19 | 0.7451145 | 2.0038557 | 0 | 0.004563333 | 0.175 | 1858 |
| NEGATIVE REGULATION OF CELL JUNCTION ASSEMBLY | NEGATIVE REG | Details ... | 18 | 0.76398754 | 2.0000577 | 0 | 0.004893971 | 0.193 | 2160 |
| MUSCLE ORGAN DEVELOPMENT | MUSCLE ORGA | Details ... | 230 | 0.49222553 | 1.9988596 | 0 | 0.0049667 | 0.201 | 1537 |
| EAR DEVELOPMENT | EAR DEVELOPM | Details ... | 159 | 0.5118388 | 1.9982015 | 0 | 0.004905887 | 0.202 | 2049 |
| TRABECULA FORMATION | TRABECULA FO | Details ... | 20 | 0.7541097 | 1.9968579 | 0 | 0.004869513 | 0.205 | 1946 |
| POSITIVE REGULATION OF CARTILAGE DEVELOPMENT | POSITIVE REGU | Details ... | 21 | 0.7382876 | 1.9934283 | 0 | 0.005042414 | 0.216 | 2846 |
| NEGATIVE REGULATION OF CELLULAR RESPONSE TO GROWTH FACTO | NEGATIVE REG | Details ... | 104 | 0.53408754 | 1.9905947 | 0 | 0.005230209 | 0.226 | 1206 |
